# From Genomics to Integrative Taxonomy? The Case Study of *Pocillopora* Corals

**DOI:** 10.1101/2022.10.04.510617

**Authors:** Nicolas Oury, Cyril Noël, Stefano Mona, Didier Aurelle, Helene Magalon

## Abstract

With the advent of genomics, sequencing thousands of loci from hundreds of individuals now appears feasible at reasonable costs, allowing complex phylogenies to be resolved. This is particularly relevant for cnidarians, for which insufficient data due to the small number of currently available markers, coupled with difficulties in inferring gene trees and morphological incongruences, encrypts species boundaries, thereby blurring the study and conservation of these organisms. Yet, can genomics alone be used to delimit species in an integrative taxonomic context? Here, focusing on the coral genus *Pocillopora*, which plays key roles in Indo-Pacific reef ecosystems but has challenged taxonomists for decades, we explored and discussed the usefulness of multiple criteria (genetics, morphology, biogeography and symbiosis ecology) to delimit species of this genus. Phylogenetic inferences, clustering approaches and species delimitation methods based on genome-wide single-nucleotide polymorphisms (SNPs) were first used to resolve *Pocillopora* phylogeny and propose genomic species hypotheses from 356 colonies sampled across the Indo-Pacific (western Indian Ocean, tropical southwestern Pacific and south-east Polynesia). These species hypotheses were then compared to previous genetic evidences, as well as to evidences based on morphology, biogeography and symbiosis. Genomics allowed to delimit 21 species hypotheses where only seven are currently recognised based on current taxonomy. Moreover, 13 species were strongly supported by all approaches, either confirming their currently recognised species status, or supporting the presence of new species that need to be formally described. Some of the other genomic species hypotheses were supported by biogeographic or symbiosis evidences, but additional investigations are needed to state on their species status. Altogether, our results support (1) the obsolescence of macromorphology (i.e., overall colony and branches shape) but the relevance of micromorphology (i.e., corallite structures) to refine *Pocillopora* species limits, (2) the need to identify molecularly species prior to their study, as morphology can blur species identification on the field, (3) the relevance of the mtORF (coupled with other markers in some cases) as a diagnostic marker of most species, and (4) the need for a taxonomical revision in the *Pocillopora* genus. These results give new insights into the usefulness of multiple criteria for resolving *Pocillopora* species limits and will ultimately provide helpful insights for the conservation of the species from this scleractinian genus. [biogeography; cryptic species delimitation; Indo-Pacific; microsatellites; morphology; phylogenetics; single-nucleotide polymorphism (SNP); Symbiodiniaceae]

---

Efficiently protecting species implies knowing their life history traits and functioning. This requires accurately defining species limits, something that may sound trivial but has long been debated (e.g., Mayden 1997; De Queiroz 2007). Indeed, several species concepts, based on more or less compatible criteria, have previously been proposed (reviewed in De Queiroz 2007). Each concept has its own advantages, but also its own approximations of the biological truth, so it now appears evident to integrate multiple criteria and go towards a unified species concept (De Queiroz 2005). However, it is not always obvious how these criteria should be combined, and some may be more informative than others or give contradictory insights, depending on organisms.

Accurately delimiting species is particularly essential for scleractinian corals, the cornerstone of coral reefs, which are experiencing critical decline worldwide (Hughes et al. 2017, 2018, 2019; Heron et al. 2018), attributable both to local (e.g., coastal development, over-fishing, pollution) and global (e.g., climate change) pressures. Coral taxonomy initially relied on skeleton morphological traits (i.e., *corallum* macromorphology and corallite microstructure; Vaughan and Wells 1943; Wells 1956; Chevalier 1971; Veron 2000), but phenotypic plasticity hampers reliable species delimitation on this basis (see Todd 2008). With the advent of genetics, molecular approaches have been used to explore species boundaries, revealing incongruences of conventional systematics within many scleractinian genera (e.g., Keshavmurthy et al. 2013; Schmidt-Roach et al. 2014; Gélin et al. 2017b; Cunha et al. 2019; Arrigoni et al. 2020). Nuclear internal transcribed spacers (ITS) and mitochondrial markers have been extensively used in phylogenetic inferences (e.g., Benzoni et al. 2007; Gélin et al. 2017b; Nakajima et al. 2017). However, intra-individual and intra-specific variations for the formers (van Oppen et al. 2000; Chen et al. 2004; Vollmer and Palumbi 2004), and relatively slow evolutionary rates for the latter (van Oppen et al. 1999; Shearer et al. 2002; Hellberg 2006), make these markers usually not informative for species delimitation in most genera (e.g., Forsman et al. 2009; Terraneo et al. 2016). Additionally, the small number of currently available markers, coupled with hybridisation (Willis et al. 2006; Combosch et al. 2008; Richards et al. 2008), introgression (Combosch and Vollmer 2015; Hellberg et al. 2016) and incomplete lineage sorting in gene trees (van Oppen et al. 2001; Fukami et al. 2008) blur phylogenetic relationships between taxa.

The recent development of high-throughput sequencing technologies now enables the cost-effective target of large numbers of loci from hundreds of individuals from virtually any species (Metzker 2010). These methods appear particularly promising to resolve complex phylogenies such as those involving scleractinian corals (e.g., Forsman et al. 2017; Cunha et al. 2019; Arrigoni et al. 2020). In particular, restriction-site associated DNA sequencing (RADseq; Baird et al. 2008) and sequence capture (also called target enrichment; Hodges et al. 2007; Gnirke et al. 2009) are increasingly used, from population genetics to phylogenetic studies (see Narum et al. 2013 for a review). While RADseq typically generates datasets of anonymous loci, sequence capture enables the deep sequencing of previously identified loci of interest, but needs existing genomic resources to design probes (Davey et al. 2011; Harvey et al. 2016). When such genomic resources are unavailable for the species of interest, probes from genomic regions that are conserved across divergent taxa [e.g., ultraconserved elements (UCEs); https://www.ultraconserved.org/] can be used (Faircloth et al. 2012, 2013; McCormack et al. 2012).

The coral genus *Pocillopora* Lamarck, 1816 (Scleractinia, Pocilloporidae) represents a key component of coral reef ecosystems from the Indo-Pacific and the Red Sea (Veron 2000), as its branching colonies are abundant and sometimes the main bio-constructors (e.g., Benzoni et al. 2003). However, its taxonomy remains challenging, and the extraordinary range of morphological diversity among its colonies has led to the coining of more than 40 species names (Hoeksema and Cairns 2022). Defining morphospecies based on morphological characters (shape and organisation of branches and verrucae), Veron (2000) recognised only 17 of them. Recent genetic studies identified several cryptic species and lineages within those morphospecies (see Gélin et al. 2017b for a review). As an illustration, the so-called *P. damicornis* (Linnaeus, 1758) was disentangled in five genetic lineages: *P. damicornis* types *a, ß, δ, γ*, and *ε* (Schmidt-Roach et al. 2012), *a posteriori* defined as five distinct species and named *P. damicornis* (Linnaeus, 1758), *P. acuta* Lamarck, 1816, *P. aliciae* Schmidt-Roach, Miller & Andreakis 2013, *P. verrucosa* (Ellis & Solander, 1786), and *P. brevicornis* Lamarck, 1816, respectively (Schmidt-Roach et al. 2014). Following this taxonomical revision of the genus, 21 valid *Pocillopora* species are currently accepted (Hoeksema and Cairns 2022). Besides, using species delimitation methods based on sequence data from colonies sampled in three marine provinces (western Indian Ocean, tropical southwestern Pacific and south-east Polynesia), Gélin et al. (2017b) defined within the *Pocillopora* genus 16 primary species hypotheses (PSHs *sensu* Pante et al. 2015). Some of these PSHs correspond to currently accepted species, but others do not and would therefore represent undescribed species. Additionally, using microsatellites, some PSHs were partitioned into several secondary species hypotheses (SSHs *sensu* Pante et al. 2015), themselves partitioned into several divergent but sympatric genetic clusters (Gélin et al. 2017a, 2017b, 2018a, 2018b; Oury et al. 2020a, 2021, 2022). This genetic partitioning questions species limits and shelves taxonomic uncertainties for which traditional genetic markers appear not enough resolutive. So far, only two studies (Johnston et al. 2017, 2022) have inferred phylogenetic relationships among species of the *Pocillopora* genus using high-throughput sequencing data (ezRAD; Toonen et al. 2013). In both cases, they resolved clear monophyletic groups that coincide with previously published mitochondrial clades based on the so-called open reading frame marker (mtORF; a putative protein-coding region of unknown function; Flot and Tillier 2007). However, their samplings were relatively concise (13 and 55 samples from seven morphospecies) and restricted to the Pacific, missing a huge part of the high diversity of this genus.

Here, considering a subset of 356 *Pocillopora* colonies from the same sampling set as in Gélin et al. (2017b), representing the totality of the PSHs, SSHs and clusters previously identified (see Gélin et al. 2017a, 2017b, 2018a, 2018b; Oury et al. 2020a, 2021, 2022), as well as all morphotypes sampled, we used sequence capture of UCEs and exon loci to collect single-nucleotide polymorphisms (SNPs). Maximum-likelihood and Bayesian phylogenetic inferences, clustering approaches and species delimitation methods based on SNP data were applied to resolve the *Pocillopora* phylogeny and define genomic species hypotheses, which were compared to previous genetic partitionings of the genus (i.e., the PSHs, SSHs and clusters previously defined based on the mtORF marker and microsatellites). Genetic evidences were then confronted to other criteria (macro- and micromorphology, biogeography and associated Symbiodiniaceae communities), to propose species delimitation of *Pocillopora* in an integrative taxonomic context. The usefulness of each criterion and its integration were then discussed.

## Materials and Methods

Detailed materials and methods, including sampling, sequencing and analytical methods, are available in Appendices 1-4.

### Sampling

The sampling was the same as in Gélin et al. (2017b) and comprised ca. 9,000 *Pocillopora* colonies from various habitats and morphotypes, from three marine provinces: the western Indian Ocean (WIO), the tropical southwestern Pacific (TSP) and the south-east Polynesia (SEP). All colonies were previously genotyped with 13 microsatellites and for a subset, we also sequenced the mitochondrial ORF locus (mtORF; see Gélin et al. 2017b for more details). Each colony was thus assigned beforehand a primary and a secondary species hypothesis (PSH and SSH, respectively; *sensu* Gélin et al. 2017b), and a cluster when appropriate, based on these genetic data (see, for example, Oury et al. 2021). From now, to simplify the reading, PSHs that were not subdivided into several SSHs are designated SSHs, keeping their corresponding number (e.g., PSH01 switches to SSH01). These SSHs remain easily recognisable as no lowercase letter follows the number.

In this study, a subset of 356 *Pocillopora* colonies (Table S1 & Fig. S1 in Appendix 1), covering the totality of the localities and morphotypes sampled, as well as all SSHs and clusters, was considered to maximise the genetic diversity explored. Four *Seriatopora hystrix* and four *Stylophora pistillata* colonies were also included as outgroups.

### Molecular Analyses

#### Sequencing and bioinformatics processing

All 364 colonies, plus eight sequencing replicates, were sequenced following a target enrichment protocol of 1,248 ultraconserved elements (UCEs) and 1,385 exon loci (Quattrini et al. 2018; see Appendix 2 for more details). The bioinformatics pipeline, from demultiplexed reads to final SNP datasets, is detailed in Appendix 2. Three individuals were discarded due to too many missing data (> 60%).

#### Phylogenomic analyses

All following analyses (detailed in Appendix 2) were performed on two datasets, one keeping all filtered SNPs and the other keeping one randomly chosen SNP per locus to reduce the effect of linkage disequilibrium. Available *Pocillopora* genomes [i.e., *P. acuta* (Vidal-Dupiol et al. 2019), *P. damicornis* (Cunning et al. 2018) and *P. verrucosa* (Buitrago-López et al. 2020)] were also included by retrieving the genotypes of the SNPs corresponding to each dataset. Phylogenetic relationships were investigated using maximum likelihood (ML) and Bayesian inferences with RAxML-NG v0.9.0 (Kozlov et al. 2019) and BEAST v2.6.3 (Bouckaert et al. 2019), respectively, both using the GTR+G model. To support the phylogenomic analyses and further explore the genetic partitioning of the datasets, several clustering approaches were used. First, assignment tests were performed with Structure v2.3.4 (Pritchard et al. 2000), sNMF (Frichot et al. 2014) and discriminant analyses of principal components (dAPC; Jombart et al. 2010). Signals of admixture were further investigated with NewHybrids v1.1 (Anderson and Thompson 2002). Second, Nei (1972) individual genetic distances were computed with the R v4.0.4 (R Core Team 2021) library *‘StAMPP’* (Pembleton et al. 2013), and then used to build a minimum spanning tree (MST) and an unrooted equal-angle split network using EDENetworks v2.18 (Kivelä et al. 2015) and SplitsTree v4.15.1 (Huson and Bryant 2006), respectively. Finally, SSH and cluster assignments issued from microsatellite data (Gélin et al. 2017a, 2017b, 2018a, 2018b; Oury et al. 2020a, 2021, 2022) were compared to the groups identified with all above analyses, named hereafter genomic species hypotheses (GSHs). *F_ST_* (Weir and Cockerham 1984) were computed with *‘StAMPP’* (Pembleton et al. 2013) for each pair of GSHs, and the resulting matrix was clustered using the *heatmap.2* function from the R library *‘gplots ‘* (Warnes et al. 2020).

As some GSHs did not include any individual whose mtORF had previously been sequenced, and in order to retrieve the correspondence with previous studies, we completed the set of mtORF-sequenced colonies following Gélin et al. (2017b), and further sequenced a subset of colonies for the PocHistone, a recently discovered marker partly mapped to partial histone 3 genes from other cnidarians, and allowing to identify *P. grandis* (the senior synonym of *P. eydouxi*) colonies (Johnston et al. 2018). The same laboratory protocol and analyses as for the mtORF in Gélin et al. (2017b) were used (Appendix 2).

#### Species delimitation analyses and divergence time estimation

To confirm GSHs, Bayes factor delimitation with genomic data (BFD*; Leaché et al. 2014) was used to test several possible species delimitation models, using the SNAPP package (Bryant et al. 2012) implemented in BEAST v2.6.3 (Bouckaert et al. 2019; details in Appendix 2). To deal with computationally intensive demands from SNAPP, we first tested a batch of species delimitation scenarios to confirm the four main clades from the phylogeny (arbitrarily defined as monophyletic groups of individuals separated by nucleotide substitution per site distances of at least 0.4 on the ML tree). Then, several possible species delimitation models were tested within each clade separately, from one single species to the number of GSHs found for each clade (Table S5 in Appendix 2). Models were compared and ranked using their marginal likelihood estimate (MLE) and by calculating the Bayes factor (BF; Kass and Raftery 1995).

For each of the best-supported model (except for Clade 1, constituted of a single GSH), a coalescent-based tree was calculated with SNAPP. DensiTree v2.2.7 (Bouckaert 2010) was used to visualise the posterior distribution of topologies as cladograms, hence allowing for a clear depiction of uncertainties in the topology. Finally, GSH divergence times were estimated with the BEAST package SNAPPER v1.0.1 (Stoltz et al. 2021; Appendix 2). The divergence between *Pocillopora* and outgroups was constrained to the middle-end Paleogene (28.4-42.7 Ma; Simpson et al. 2011).

### Macro- and Micromorphological Analyses

In order to compare previously described morphospecies with GSHs (defined above), each colony was attributed a morphotype (or several when morphology was unclear), determined only by its *corallum* macromorphology [branch shape and thickness, size and uniformity of verrucae, and overall growth form as described in Veron (2000) and Schmidt-Roach et al. (2014)].

A subset of 10 colonies per GSH were also randomly selected for micromorphological observations of the bleached skeletons (particularly of the corallite structures) using scanning electron microscopy (SEM). A collection of skeleton images was thus obtained for each specimen, and multiple measurements of seven quantitative variables (e.g., corallite and columella diameters; see Appendix 3 for details) were done with ImageJ2 (Rueden et al. 2017; https://imagej.nih.gov/ij/). A non-parametric permutational multivariate anova (PERMANOVA) was then performed using the R library *‘RVAideMemoire’* (Hervé 2021) with the GSHs as factor. Each metric was analysed separately using a non-parametric permutational anova. Two additional categorical variables were also considered, and a factorial analysis of mixed data (FAMD) was performed for all nine variables using the R library *‘FactoMineR’* (Lê et al. 2008). A reference specimen representative of each species enclosed in the latest *Pocillopora* taxonomic revision (Schmidt-Roach et al. 2014) was included by measuring the variables on the images incorporated.

### Characterisation of Associated Symbiodiniaceae

Symbiodiniaceae communities were characterised for a subset of colonies (ca. 15 per GSH, when available; including three replicates) by high-throughput sequencing the ribosomal RNA internal transcribed spacer 2 (ITS2; see Appendix 4 for details). Reads were processed with the SAMBA v3.0.1 workflow (https://github.com/ifremer-bioinformatics/samba). Resulting operational taxonomic units (OTUs) were taxonomically assigned by querying a custom reference database of Symbiodiniaceae ITS2 adapted from the one available in SymPortal (downloaded on 13/01/2022; Hume et al. 2019). Taxonomic affiliations of the OTUs were confirmed by reconstructing the phylogenetic relationships among them using MAFFT v7.713 (Katoh and Standley 2013) and FastTree v2.1.11 (GTR+CAT model; Price et al. 2009). OTUs and individuals with less than 10 and 500 sequences, respectively, were then removed to reduce possible sequencing errors. Alpha diversity metrics (Chao1 and Shannon) were computed at the OTU level with the R library *‘vegan’* (Oksanen et al. 2020) and compared using non-parametric permutational ANOVA performed with the R library *‘RVAideMemoire’* (Hervé 2021), with the GSHs or the localities as factor. Finally, a nonmetric multidimensional scaling (NMDS) using Bray and Curtis (1957) dissimilarity index was performed to assess community similarity.

Species hypotheses delimited with each criterion (genomics, genetics, macro- and micromorphology, and symbiosis ecology) were then compared in an integrative species delimitation context. Sampling sites were also integrated to identify sympatric or allopatric GSHs. We then discussed the usefulness of each criterion for the delimitation of *Pocillopora* species.

## Results

### Molecular Analyses

#### Sequencing and bioinformatics processing

A total of 1.6 × 10^9^ reads (2.5 × 10^11^ bp) were produced with a highly variable number of reads per individual [varying from 9.1 × 10^3^ to 8.2 × 10^6^ reads; mean ± s.e. = (4.4 ± 0.1) × 10^6^ reads], but only three individuals (*a posteriori* removed) had less than a million reads. Quality controls and adapter trims then led to the removal of 3.0% of the bases. From the resulting trimmed reads, between 41.0% and 86.2% reads per individual were successfully mapped on the reference sequences (mean ± s.e. = 78.3 ± 0.4%), with a mean coverage depth (± s.e.) of 60.2× (± 0.1). Finally, SNPs calling and filtering (Table S4 in Appendix 2) led to two datasets: one including all SNPs (361 individuals × 17,465 SNPs; 5.8% missing data) and the other keeping randomly one SNP per locus (361 individuals × 1,559 SNPs; 6.0% missing data), with mean SNP coverage depths (± s.e.) of 85.8× (± 0.4) and 76.1× (± 1.3), respectively.

#### Phylogenomic analyses

All results were very consistent between both datasets (i.e., with one or all SNPs per locus). Thus, only results with one SNP per locus are presented below, but results keeping all SNPs are provided in Appendix 2. The phylogenetic trees inferred both with RAxML and BEAST gave similar tree topologies and recovered four strongly supported clades (Clades 1-4; i.e., monophyletic groups of individuals chosen here as separated by at least 0.4 nucleotide substitution per site on the ML tree; Fig. 1), themselves split (except Clade 1) into a total of 21 genomic species hypotheses (GSHs). Each GSH (except three) was restricted to a single marine province (Table 1 & Fig. 1) and several GSHs were thus sympatric (12 in the WIO, 11 in the TSP and 2 in the SEP; Table S6 in Appendix 2), supporting evolutionary rather than geographic reproductive isolations. Moreover, most of the GSHs (see below for the exceptions) roughly corresponded to previously defined secondary species hypotheses on the basis of microsatellites (SSHs *sensu* Gélin et al. 2017b). Therefore, to avoid introducing a new nomenclature and to ease correspondence with earlier works, the GSHs were named according to the corresponding SSHs (e.g., the GSH corresponding to SSH01 was named GSH01; Table 1 & Fig. 1).

**Fig. 1.**
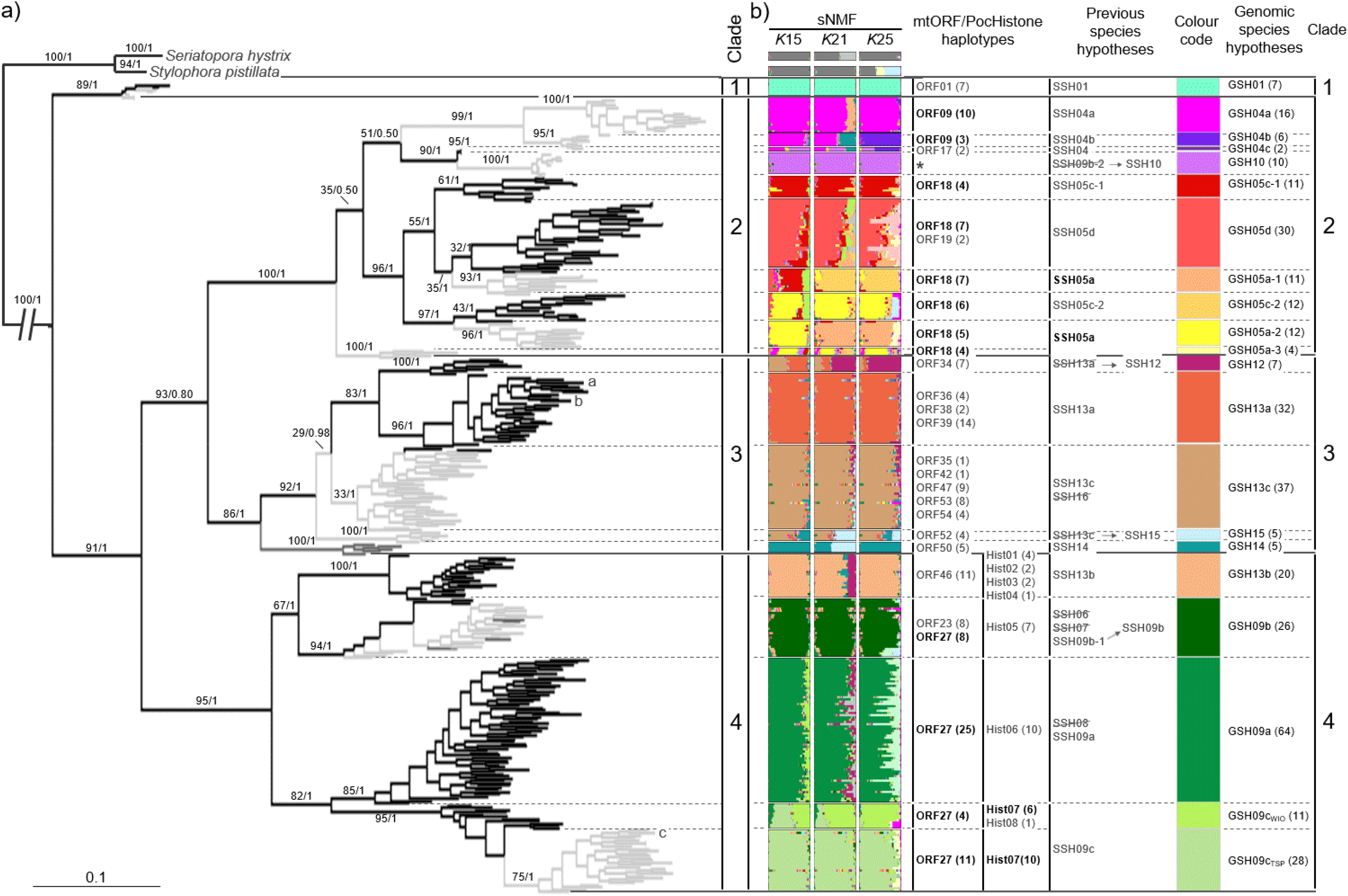
*Pocillopora* phylogeny reconstructed with one SNP per locus (361 individuals × 1,559 SNPs). (a) maximum likelihood (ML) phylogenetic tree. Branches are coloured according to marine provinces [black: western Indian Ocean (WIO); light grey: tropical southwestern Pacific (TSP); dark grey: south-east Polynesia (SEP)], and branch support, based on ML bootstrap analyses (first number) and Bayesian posterior probabilities (second number), is indicated for branches supporting the genomic species hypotheses (GSHs; delimited by dashed lines; full lines delimit the clades indicated alongside). Published genomes are indicated by lowercase letters [a: *P. verrucosa* (Buitrago-López et al. 2020); b: *P. acuta* (Vidal-Dupiol et al. 2019); c: *P. damicornis* (Cunning et al. 2018)]. (b) sNMF assignments at *K* = 15, *K* = 21 and *K* = 25, mitochondrial open reading frame (mtORF) and PocHistone haplotypes repartition [number of colonies in parentheses; haplotypes in bold are found in several GSHs; *: ORF30 (3) & ORF31(4)], corresponding secondary species hypotheses (SSHs) and clusters (defined in Gélin et al. 2017a, 2017b, 2018a, 2018b; Oury et al. 2020a, 2021, 2022), genomic species hypotheses (GSHs; number of colonies in parentheses), clades and colour code retained throughout this study.

**Table 1.**
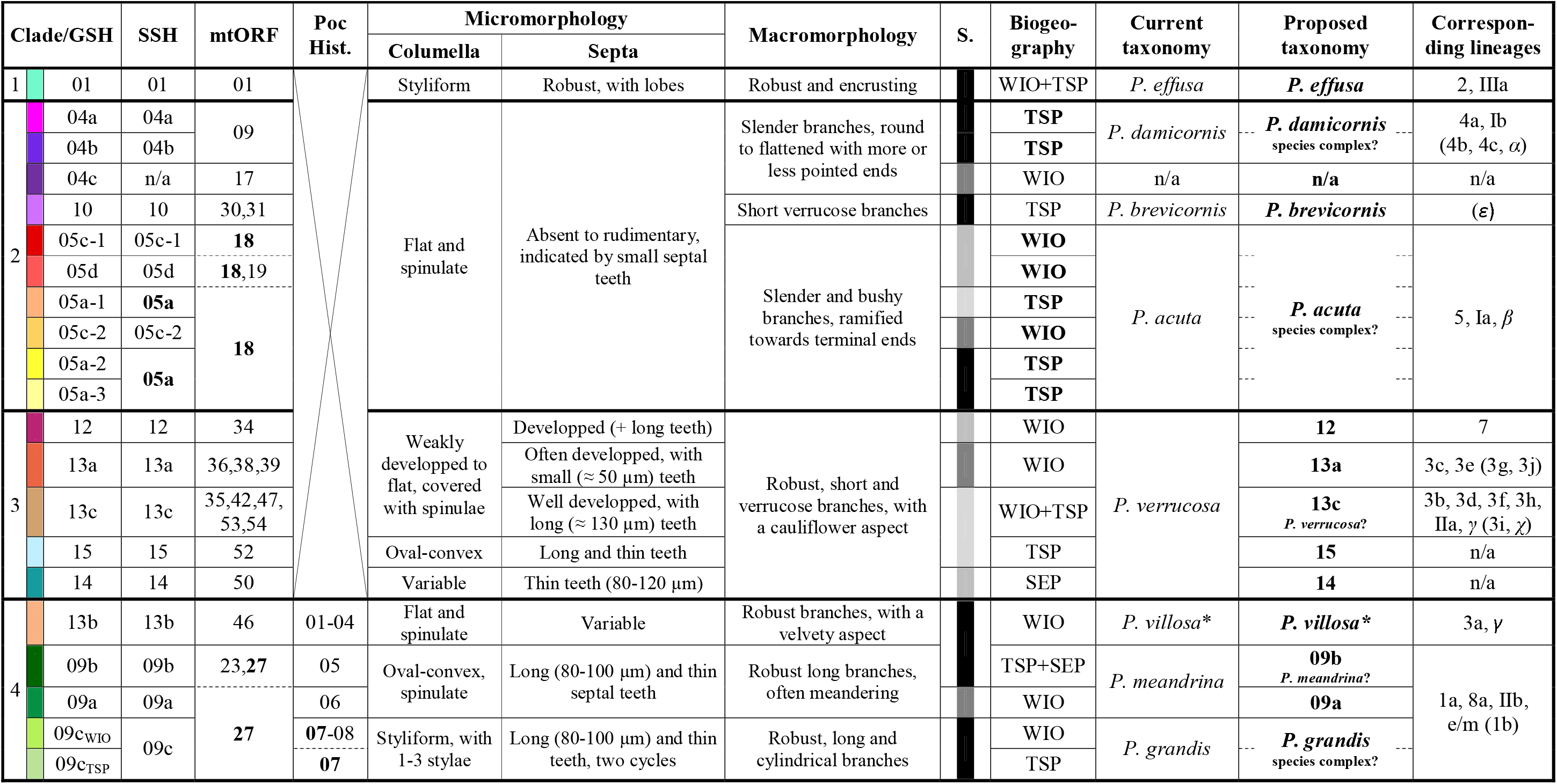
Summary of the different approaches exploring *Pocillopora* species limits: genetics [genomic and corresponding corrected secondary species hypotheses (GSHs and SSHs *sensu* Gélin et al. 2017b, respectively), mtORF (mitochondrial open reading frame) and PocHist. (PocHistone) haplotypes; values in bold are retrieved in several GSHs], micro- and macromorphological, symbiosis (S.; each colour denotes distinct dominant Symbiodiniaceae; see Fig. S14 in Appendix 4) and geographical (WIO: western Indian Ocean; TSP: tropical southwestern Pacific; SEP: south-east Polynesia; values in bold highlight sympatric GSHs within a species complex) evidences. Corresponding lineages from previous studies are also indicated (arabic numerals correspond to types from Pinzón et al. 2013, roman numerals to clades from Marti-Puig et al. 2014 and greek letters to types from Schmidt-Roach et al. 2014; lineages in parentheses were extrapolated from SSHs).*: *P. villosa nomen nudum* was proposed by Gélin et al. (2017b) but does not correspond to a currently valid species.

SSH06, SSH07, SSH08 and SSH16, previously defined from few individuals, were not retrieved here as the corresponding individuals were grouped with those from SSH09a, SSH09b-1 or SSH13c. Similarly, SSH09b-2 was grouped with SSH10 and was far apart from the rest of SSH09 *sensu lato*, suggesting that individuals from SSH09b-2 correspond to GSH10 (as observed with the mtORF). GSH09b, therefore, corresponds to only SSH09b-1. SSH12 and SSH15, previously grouped with SSH13a and SSH13c, respectively, using microsatellites (Gélin et al. 2017b), were retrieved, confirming the distinction between them. Conversely, the over-partitioning previously found with microsatellites inside SSH04a (Oury et al. 2020a), SSH05d (Clusters 1 and 4 in Gélin et al. 2018b), SSH09a, SSH09c (Gélin et al. 2018a) and SSH13c (Oury et al. 2021) was not retrieved, while the one found into SSH05c (Clusters 2 and 3 in Gélin et al. 2018b; *a posteriori* named SSH05c-1 and SSH05c-2 in Oury et al. 2020b) was. Finally, SSH05a was split into three new groups (GSH05a-1, GSH05a-2 and GSH05a-3), and SSH09c split into two new groups (GSH09c_WIO_ and GSH09c_TSP_) restricted to the WIO or the TSP, respectively (Table 1 & Fig. 1).

The three assignment methods, although estimating different admixture rates and suggesting different optimal *K* according to their respective criterion, gave similar results, retrieving almost all 21 GSHs (Fig. 1 & S3-S4 in Appendix 2). In particular, sNMF and Structure highlighted introgression signals among several GSHs, compatible with allopatrism, that were further investigated with NewHybrids. This was notably the case within GSH05 *sensu lato*, but also with GSH12 as hybrids between GSH13a and GSH13c, GSH15 between GSH13c and GSH14, or GSH09c_WIO_ between GSH09a and GSH09c_TSP_ (Fig. 1 & S3-S4). No individual was assigned to a hybrid class (i.e., F1, F2 or backcrosses), except the GSH09c_WIO_ ones that were assigned as F2 hybrids from GSH09a and GSH09c_TSP_ (data not shown). The two networks clustering also retrieved the 21 GSHs (Fig. S5 in Appendix 2). Thus, published genomes were assigned to the same GSHs with all datasets and analyses (Fig. 1 & Appendix 2): two [*P. acuta* (Vidal-Dupiol et al. 2019) and *P. verrucosa* (Buitrago-López et al. 2020)] were assigned to GSH13a (currently considered as *P. verrucosa*) and the third [*P. damicornis* (Cunning et al. 2018)] to GSH09c_TSP_ (*P. grandis*).

Finally, all pairwise *F_ST_* were significantly positive (*P* < 0.001***) and the dendrogram topology obtained from the clustering of *F_ST_* values was comparable to phylogenies (Fig. S6 in Appendix 2). Intra-clade *F_ST_* ranged from 0.092*** to 0.689*** [mean (± s.e.) = 0.332 ± 0.011], while inter-clade ones ranged from 0.420*** to 0.795*** [mean (± s.e.) = 0.551 ± 0.004; Table S7 in Appendix 2].

For the mtORF, 59 additional colonies were sequenced, but no new haplotype was found. Each haplotype (except three) was restricted to a single GSH (Table 1 & Fig. 1), confirming previous results from Gélin et al. (2017b). In particular, ORF27 was found in GSH09 *sensu lato*, thus corresponding to *P. grandis* and/or *P. meandrina*. To distinguish both species, we sequenced 10 colonies of each of the five GSHs from Clade 4 for the PocHistone. Among the 43 successfully sequenced colonies, no heterozygote was found and eight novel 588 bp-haplotypes were identified (Hist01-08; GenBank accession numbers ON155826-ON155833; Table S9 in Appendix 2), to which we added the two available in GenBank (MG587096 and MG587097, corresponding to *P. grandis* and *P. meandrina*, respectively; Johnston et al. 2018; Table S9). All, but one haplotype (Hist07), were restricted to a single GSH (Table 1 & Fig. 1). Hist07 and Hist08 had the *P. grandis* diagnostic SNP, suggesting that GSH09c corresponds to *P. grandis* (Table S9). The reconstructed PocHistone phylogeny consistently regrouped these two haplotypes with the *P. grandis* one from Johnston et al. (2018), but all other haplotypes were grouped inconsistently with the defined GSHs (Fig. S7 in Appendix 2).

#### Species delimitation analyses

Among the scenarios tested to delimit the four main clades (Clades 1-4; Table S5 in Appendix 2), the best-supported model was the one separating those four clades (model 4: MLE = −11,868.07, BF = –). The three other models were ranked with a decreasing number of clades (i.e., from three clades to a single one; 1,294.87 < BF < 4,706.03; Table S5). Within each clade, the model with the lowest MLE was the one separating colonies according to the different GSHs previously identified based on phylogenomic and clustering analyses. The best-supported model for Clade 1 was therefore the 1-species model (GSH01; model 1.1: MLE = −1,112.45, BF = –), followed by the 1-species-per-ocean model (model 1.2: MLE = −1,381.70, BF = 538.49). For Clade 2, analyses supported the 10-species model (model 2.17: MLE = −20,955.08, BF = –), followed by the models lumping GSH05a-1 and GSH05d (model 2.16: MLE = −21098.78, BF = 287.39) or GSH05a-2 and GSH05c-2 (model 2.14: MLE = −21,408.68, BF = 907.18). Finally, for Clades 3 and 4, the best supported models were the 5-species ones (model 3.8: MLE = −10,529.96, BF = –; model 4.11: MLE = −12,717.20, BF = –). However, Clade 4 5-species model was closely followed by the model lumping GSH09c_WIO_ and GSH09c_TSP_ (model 4.10: MLE = −12,763.15, BF = 91.89; Table S5). In summary, BFD* supported the 21 GSHs identified with the phylogenomic analyses.

A total of four species trees were estimated (i.e., one for the best-supported model in the initial batch of scenarios, and then one for each best-supported model for scenarios within Clades 2-4 separately; Fig. 2a). For the initial batch of scenarios, three (out of three) consensus tree topologies were identified in the 95% HPD set, and all grouped Clades 1 and 4 together, whereas Clade 1 was the most distant group according to all previous analyses. The three topologies differed in whether Clades 2 or 3 shared a direct common ancestor with Clades 1 and 4 (59.1% and 22.0% of the trees, respectively), or together (18.9%; Fig. S8 in Appendix 2). For Clade 2, two (out of nine) consensus tree topologies were identified in the 95% HPD set. Both topologies were very similar and consistent with previous analyses, except that GSH05a-3 was alternatively grouped with or without GSH05a-2 and GSH05c-2 (52.1% and 44.3% of the trees, respectively; Fig. 2 & S8).

**Fig. 2.**
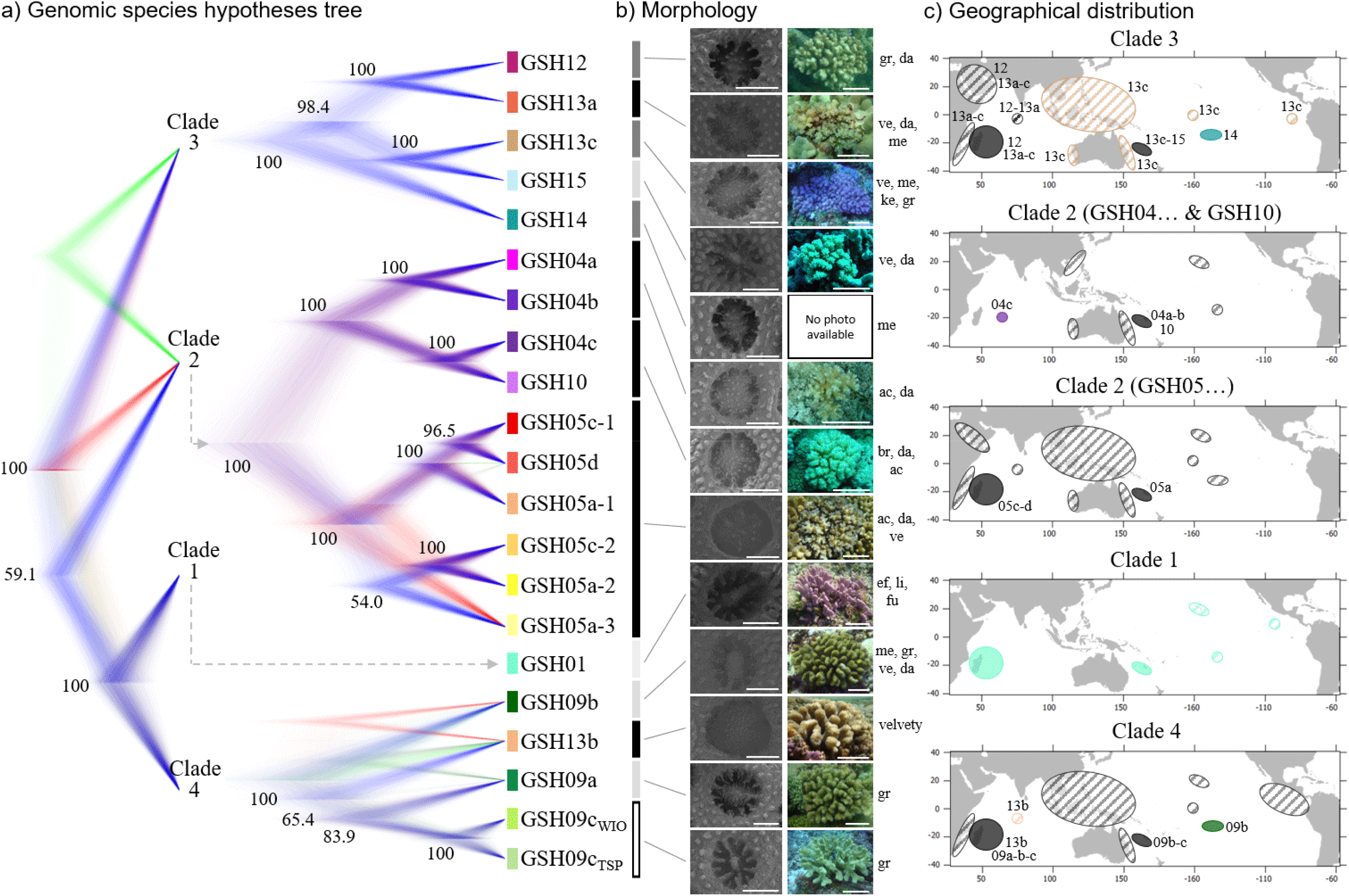
Species tree estimation for the 21 delimited *Pocillopora* genomic species hypotheses (GSHs). (a) complete set of consensus trees visualised with DensiTree for each best-supported model (i.e., for the initial batch of scenarios, and then for scenarios within Clades 2-4 separately). Higher density areas indicate greater topology agreement and different colours represent different topologies (trees with the highest clade credibility in blue). Node supports (Bayesian posterior probabilities) > 50% are indicated. (b) micro- (scale ≈ 500 μm) and macromorphological (scale ≈ 10 cm) overview of the GSHs (characteristic features only; see Appendix 5 for more illustrations). Bar colours symbolise separate micromorphological groups on the factorial analysis of mixed data (FAMD; see also Fig. S10 in Appendix 3) and morphotypes encountered in this study (sorted by occurrence) are indicated alongside photographs (ac: acuta, br: brevicornis, da: damicornis, ef: effusa, fu: fungiformis, gr: grandis, ke: kelleheli, li: ligulata, me: meandrina and ve: verrucosa). (c) geographical distribution of each GSH. Filled circles represent data from this study while hashed ones were taken from the literature [based on mtORF identifications; colours refer to the GSH and black denotes multiple sympatric GSHs (indicated alongside) or ambiguous identifications (no GSH indicated)].

Only one (out of four) consensus tree topology was identified in the 95% HPD set for Clade 3 (representing 98.4% of the trees). This topology was consistent with previous analyses, i.e. grouping GSH12 and GSH13a on one side, GSH13c and GSH15 on the other side, and GSH14 being the most distant species (Fig. 2). Finally, for Clade 4, a total of 15 consensus tree topologies were found, of which five were in the 95% HPD set. All topologies identified GSH09c_WIO_ and GSH09c_TSP_ as sister species, but then differed in whether GSH09a shared a direct common ancestor with them (83.9% of the trees) and whether GSH09b and GSH13b were sister species (17.8%) or progressive outgroups (65.4%; Fig. 2 & S8).

The time-calibrated phylogeny indicated a first divergence within the *Pocillopora* genus 20.4 Ma, separating on one side Clades 1 and 4, and on the other side Clades 2 and 3. Clade pairs then diverged 17.4 Ma and 16.0 Ma, respectively (Fig. 3). Each clade then went through a first diversification period in the late Miocene (6.5-7.5 Ma), followed by a second period in the Pliocene and the Quaternary (i.e., from 4.5 Ma). Thus, almost all *Pocillopora* GSHs appeared relatively recently (Fig. 3).

**Fig. 3.**
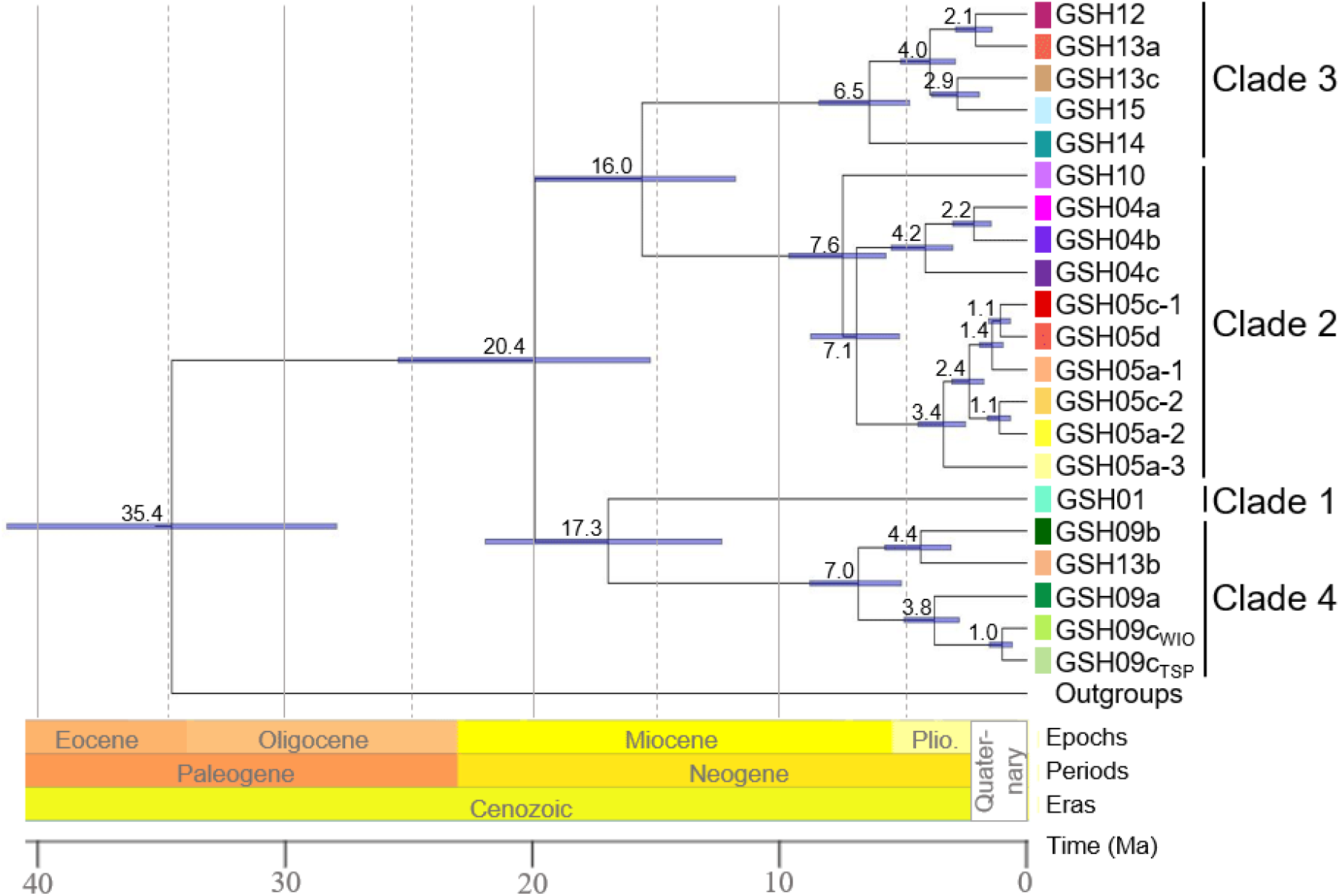
Time-calibrated phylogeny of *Pocillopora* genomic species hypotheses (GSHs). Values above nodes indicate median node ages and blue bars represent the 95% highest posterior density (HPD) interval. Plio.: Pliocene.

### Macro- and Micromorphological Analyses

Morphotypes based on macromorphology were not exclusive of a single GSH. Indeed, each GSH usually grouped colonies with a dominant morphotype, but also included several other morphotypes (e.g., GSH09b mostly grouped *P. meandrina-like* colonies, but also *P. damicornis-* like, *P. grandis-like* or *P. verrucosa-like*). Reciprocally, colonies from different GSHs can share the same morphotype (e.g., *P. damicornis-like* colonies were found in 14 GSHs; Fig. 2). Clades 1 and 4 were mostly characterised by robust morphs with large branches, while Clades 2 and 3 grouped more stunted colonies (Fig. 2 & Appendix 5).

Concerning micromorphology, intraspecific variations were smaller (Fig. 2 & S9 in Appendix 3). In particular, all species from Clades 1 and 4 (except GSH13b) and GSH15 were characterised by a styliform (GSH01, GSH09c_WIO_ and GSH09c_TSP_) or oval-convex (GSH09a, GSH09b and GSH15) columella, while all other species had a flat, more or less spinulate columella. Accordingly, significant differences among GSHs were found for the columella diameter variables (v6 and v7; non-parametric permutational anovas; v6: *F*_(5,46)_ = 92.95, *P* < 10^-3^***; v7: *F*_(5,46)_ = 98.20, *P* < 10^-3^***), distinguishing three groups: GSH01 + GSH15, GSH09a + GSH09b and GSH09c_WIO_ + GSH09c_TSP_ (pairwise permutational t tests; *P* < 0.05*; Fig. S9). Significant differences among GSHs were also found for all other five numeric morphological variables (v1-v5; non-parametric permutational anovas; 4.53 ≤ *F*_(19,150)_ ≤ 18.96; *P* < 10^-3^***), but no particular pattern was identified, except that GSHs from Clade 2, GSH13a and GSH13b had poorly developed septa (Fig. 2 & S9). The PERMANOVA and FAMD (Fig. S9 & S10 in Appendix 3) also highlighted these differences. Five micromorphological groups were thus distinguished on the first three principal components of the FAMD (explaining 68.4% of the variability): GSH01, Clade 2 + GSH13a + GSH13b, GSH12 + GSH13c + GSH14, GSH09a + GSH09b + GSH15 and GSH09c_WIO_ + GSH09c_TSP_ (Fig. 2 & S10). Detailed macromorphological and micromorphological illustrations of the GSHs are provided in Appendix 5.

### Characterisation of Associated Symbiodiniaceae

ITS2 amplicon sequencing yielded a total of 1.6 × 10^7^ reads (4.0 × 10^9^ bp) with between 1.7 × 10^4^ to 1.2 × 10^5^ reads per individual [mean ± s.e. = (6.1 ± 0.1) × 10^4^ reads]. After merging paired reads and removing chimeras, 9.0 × 10^6^ sequences were retained [with between 0 and 7.2 × 10^4^ sequences per individual; mean ± s.e. = (3.5 ± 0.0) × 10^4^ sequences], corresponding to 1,014 amplicon sequence variants that were clustered in 590 operational taxonomic units (OTUs; represented by 1 to 6.1 × 10^5^ sequences). Finally, 534 OTUs (90.5%) were taxonomically assigned, with a majority (511 OTUs, representing 97.6% of the sequences) belonging to *Cladocopium* (formerly *Symbiodinium* clade C), and mostly to clades C1 (173 OTUs and 35.6% of the sequences), C40 (267 OTUs; 56.6% of the sequences) and C42 (45 OTUs; 3.6% of the sequences). The other OTUs were assigned to *Symbiodinium* (clade A1; 6 OTUs; 0.2% of the sequences), *Durisdinium* (clade D1; 2 OTUs; < 0.1% of the sequences), *Gerakladium* (clade G3; 12 OTUs; 0.1% of the sequences) and Symbiodiniaceae clade I (clades I1 and I3; 3 OTUs; < 0.1% of the sequences). The reconstructed phylogeny based on these OTUs retrieved the five genera, with largely unresolved polytomies within *Cladocopium*, as previously observed (LaJeunesse 2005; Brener-Raffalli et al. 2018; Fig. S11 in Appendix 4). Nevertheless, such polytomies should not affect subsequent analyses, as performed at the OTU level.

From the remaining 252 individuals and 552 OTUs that passed the filtration steps, OTU richness within colonies varied from 0.14 to 2.67 for Shannon diversity index, and from 2 to 39 for Chao1 index. Both indexes were significantly different among GSHs (non-parametric permutational anova; Shannon: *F*_(20,231)_ = 3.89, *P* < 10^-3^***; Chao1: *F*_(20,231)_ = 2.96, *P* < 10^-3^***; Fig. S12 in Appendix 4), but no significant difference was found in Chao1 post-hoc tests (pairwise permutational t tests; *P* > 0.05^NS^), and no obvious pattern was found for Shannon (Fig. S12). Differences were clearer when looking at the proportion of each taxon within samples (Fig. S13 in Appendix 4). For example, individuals from GSH04c, GSH09a and GSH13a displayed mainly C1ky [38.8 ± 2.4% on average (± s.e.)], while it was almost absent in other GSHs. Similarly, GSH05c-2, GSH13c and GSH15 contained mainly C1ag (56.5 ± 4.6%) and GSH05c-1, GSH05d, GSH12 and GSH14 mainly C1d (50.0 ± 4.4%). Except few individuals, other GSHs contained almost exclusively C40c (Fig. S13). Accordingly, the NMDS based on Bray and Curtis (1957) dissimilarity index followed this partitioning with three groups on the first two principal components (explaining 41% of the variability): (1) individuals mostly composed of C1ky, (2) those mostly composed of C1ag or C1d (separated with the third principal component) and (3) individuals mostly composed of C40c (Table 1 & Fig. S14 in Appendix 4).

Concerning localities, significant differences were found for Chao1 (again, without any obvious pattern; non-parametric permutational anova; *F*_(13,238)_ = 5.94, *P* < 10^-3^***), but neither for Shannon (non-parametric permutational anova; *F*_(13,238)_ = 1.69, *P* = 0.07^NS^; Fig. S12), nor by looking the individual proportions (Fig. S13) or the NMDS (Fig. S14).

### Summary of all Evidences

Genomic analyses allowed the definition of 21 GSHs, while only four species hypotheses were distinguished if based only on Symbiodiniaceae communities, and up to 10 based only on qualitative micromorphology. Thus, combining evidences from all approaches (i.e., genomics, genetics, macro- and micromorphology, and symbiosis ecology) and considering the existence of two species only if supported by all criteria, only one single unambiguous species (corresponding to the entire genus) could be delimited (i.e., no separation appears fully supported; Table 1). Removing the Symbiodiniaceae criterion, five species, corresponding to Clades 1-3, GSH13b and GSH09 *sensu lato*, could be delimited. Then, sequentially removing the macro- and micromorphology criteria (i.e., considering all genetic evidences alone) could lead to nine and 12 species (Table 1). However, three out of the four GSHs within GSH09 *sensu lato* are split by other criteria (morphology and sometimes Symbiodiniaceae), supporting some genetic and genomic evidences. Conversely, considering two species once a single criterion separates them, genomics alone allows to distinguish all partitions. Consequently, we discuss below the usefulness of each criterion and propose a parsimonious consensus of 13 species strongly supported by most approaches (Table 1). Among them, three (*P. acuta, P. damicornis* and *P. grandis*) could represent species complexes according genetic evidences, and six were not attributed to a currently valid species, potentially representing new species (Table 1).

## Discussion

Although accurately delimiting species remains of particular importance and requires integrating multiple criteria, all investigated criteria do not provide the same resolution nor congruent insights. Not all criteria should therefore obviously be considered equally in order to define a consensus of the species limits as parsimonious as possible. As an illustration, both pigs and humans have four limbs, udders, etc., but that does not mean they belong to the same species. Conversely, eye colour does not distinguish different species in humans. Thus, in this study, focusing on the scleractinian genus *Pocillopora* across a wide range of sampled localities (18 islands or regions from three marine provinces), genetic, morphological, geographical and symbiosis data were collected and compared to define robust species limits and assess the usefulness of each criterion. The different genetic approaches allowed to delimit 21 genomic species hypotheses (GSHs) where only seven are currently recognised based on current taxonomy. Moreover, 13 species appear strongly supported by all approaches, supporting the presence of six potentially new species that need to be formally described. Some of the other GSHs were supported by biogeographic or symbiosis evidences, but additional investigations are needed to state on their species status. In any case, a taxonomical revision of the *Pocillopora* genus, taking into account evidences brought by these results and previous ones, becomes urgent. This will allow to give formal names to the new species and thus throw off the multitude of current nomenclatures based on genetic lineages which can be difficult to follow, even for specialists.

### On the (Ir)Relevance of Symbiosis Ecology to Define Species

As many scleractinian genera, *Pocillopora* species host diverse communities of symbionts (Cunning et al. 2017; Brener-Raffalli et al. 2018; Li et al. 2021; Rabbani et al. 2021). In this genus, Symbiodiniaceae are expected to be maternally transmitted (vertical transmission), as they are already present in oocytes before spawning (Sier and Olive 1994; Hirose et al. 2001; Harii et al. 2002). Such symbiont inheritance could result in species-specific associations and co-evolutions (Pinzón and LaJeunesse 2011; Schmidt-Roach et al. 2012; Johnston et al. 2022), that could also be responsible for habitat specialisations (driven by symbionts thermotolerance and photosynthetic needs; Jokiel and York 1982; Baker et al. 2013; Brener-Raffalli et al. 2018; Ros et al. 2021). Characterising associated Symbiodiniaceae communities can therefore bring additional elements to the delimitation of *Pocillopora* species, as in other scleractinian genera (Bongaerts et al. 2010; Keshavmurthy et al. 2013; Warner et al. 2015; Arrigoni et al. 2016; Forsman et al. 2020), but this does not guarantee a self-sufficient criterion.

Indeed, symbiosis ecology alone does not appear informative enough to delimit species, as evidenced by our results. We found a high prevalence of *Cladocopium* C1 (*C. goreaui*) and C40, both host-generalists, consistently with other studies on *Pocillopora* (e.g., Magalon et al. 2007; Pinzón and LaJeunesse 2011; Schmidt-Roach et al. 2012; Brener-Raffalli et al. 2018; Armstrong et al. 2021; Johnston et al. 2022). C1 variants allowed to distinguish five groups of colonies with distinct Symbiodiniaceae communities, but colonies within those groups were very distinct morphologically and genetically. Conversely, colonies from a single GSH generally shared the same communities.

These results should nevertheless be considered cautiously as (1) host-symbiont associations may vary over time and depth (Cunning et al. 2013), and (2) quantitative interpretation of metabarcoding results can be misleading (Lamb et al. 2019). First, Pinzón and LaJeunesse (2011) found that *Pocillopora* type 1 (ORF27; probably GSH09b or GSH09c_TSP_) was the only type associated to the thermotolerant *Durusdinium glynnii* (D1; Wham et al. 2017) in the tropical eastern Pacific. But it was later found in *Pocillopora* types 3a and 3b (ORF46 and ORF47; GSH13b and GSH13c, respectively), with different prevalence among sites (Cunning et al. 2013), suggesting variable host-symbiont associations. In particular, *Durusdinium* would represent an opportunist genus, replacing specialist symbionts in health-compromised (e.g., bleached) corals (Stat and Gates 2010), potentially explaining these results. In our study, *Durusdinium* was rare (representing ca. 0.5% of the sequences in a single individual), suggesting no recent bleaching event prior to sampling, and thus mature host-symbiont associations. However, horizontal (i.e., from the water column) acquisition of Symbiodiniaceae remains possible, potentially corrupting species-specific associations. Second, PCR inherent biases (reviewed in Lamb et al. 2019) can result in differential sequence amplifications, either quantitatively or qualitatively. This can result in artificial differences in Symbiodiniaceae compositions among individuals and GSHs. Conversely, rare or specific Symbiodiniaceae taxa that could be diagnostic of a GSH might not be amplified, sequenced or detected.

Species limits evidences from symbiosis ecology inferred with metabarcoding data should therefore be taken cautiously, and rather used in support of other criteria in an integrative context. Besides, this criterion has not been systematically explored in previous taxonomic revisions of scleractinian genera (e.g., Benzoni et al. 2010; Arrigoni et al. 2020, 2021; Wepfer et al. 2020), demonstrating that it is not the most relevant criterion.

### Should we Trust Morphology?

While most of the delimited GSHs grouped colonies with one major morphotype (which could be shared between GSHs), they also harboured high morphotype diversities. This demonstrates, once again (e.g., Pinzón et al. 2013; Marti-Puig et al. 2014; Gélin et al. 2017b), the obsolescence of *corallum* macromorphology to define *Pocillopora* species limits, as in other scleractinian genera (e.g., Warner et al. 2015; Shimpi et al. 2019; Bongaerts et al. 2021; Terraneo et al. 2021). Indeed, *Pocillopora* corals can display great morphological plasticity mostly driven by light and currents (Todd 2008). As an illustration, in the Gulf of California, five morphospecies have been reported (Glynn and Ault 2000), all belonging to mtORF type 1a (= ORF27; Pinzón et al. 2013). Switches from one morphospecies to another have also been demonstrated following shifts in environmental conditions (Paz-García et al. 2015a, 2015b).

Contrary to macromorphology, micromorphology brought additional insights to the refining of *Pocillopora* species limits, as in other scleractinian genera (e.g., Benzoni et al. 2007; Forsman et al. 2010; Budd and Stolarski 2011; Stefani et al. 2011; Budd et al. 2012; Arrigoni et al. 2020). Intraspecific variations were smaller, and several differences, either qualitative or quantitative, allowed to distinguish almost all GSHs. The GSHs within Clade 2 were not separated, but Schmidt-Roach et al. (2014) raised several differences that we could not recover. It is also possible that the morphological characters investigated here were not the most relevant to distinguish these GSHs (e.g., as for the number of limbs to distinguish humans and pigs).

Morphology-based criteria are thus questionable and subject to interpretation (particularly for the presence/absence of subtle characters) which, coupled with morphological plasticity, makes them unsuitable for identifying *Pocillopora* species. The misidentification of two out of the three currently available *Pocillopora* genomes perfectly illustrates this point. While the *P. verrucosa* genome (Buitrago-López et al. 2020) has been assigned to a GSH consistent with this identification (GSH13a), the two others were not [the *P. acuta* genome (Vidal-Dupiol et al. 2019) was assigned to GSH13a (currently considered as *P. verrucosa*) too and the *P. damicornis* genome (Cunning et al. 2018) to GSH09c_TSP_ (*P. grandis*)]. Surprisingly, the colony sequenced for the *P. acuta* genome has been identified molecularly using the mtORF, but the haplotype was not provided (Vidal-Dupiol et al. 2019), so we could not verify the identification.

### Exploring Species Limits: Lessons from Genomics

*Pocillopora* species limits have been extensively studied using genetic markers over the past decades (e.g., Schmidt-Roach et al. 2012; Pinzón et al. 2013; Marti-Puig et al. 2014; Gélin et al. 2017b), revealing a great diversity within some morphospecies (e.g., *P. damicornis;* Schmidt-Roach et al. 2012). Most of these previous studies used mtDNA and microsatellites to explore species limits. Only Johnston et al. (2017, 2022) inferred genetic relationships among few tens of *Pocillopora* colonies from the Pacific using genomic data. Consequently, our study represents the most extensive investigation to date of the taxonomy of the *Pocillopora* genus using genomics.

Our genomic analyses based on SNPs collected from the sequence capture of UCEs and exon loci provided very congruent results among methods and allowed the robust definition of four main clades comprising 21 GSHs. However, despite thousands of SNPs and loci analysed, we were not able to fully resolve GSH relationships, and multiple species tree topologies were inferred (Fig. 2 & S8 in Appendix 2). Recent species divergences and the presence of several closely related sister species, as well as introgression, could explain unresolved topologies. Indeed, most GSHs appeared less than 5 Ma, with a substantial number in the Quaternary (i.e., 0-2.6 Ma; Fig. 3). This suggests a recent radiation, probably linked to major geological and climatic events during the Pliocene or the Pleistocene [e.g., changes in currents (Philander and Fedorov 2003), glacial-interglacial cycles (Adams et al. 1999; Lambeck et al. 2002) and formation of the Isthmus of Panama (O’Dea et al. 2016)], as already suggested in this genus (Johnston et al. 2017). Recent divergences also suggest that some sister GSHs might still be in speciation and are in the grey zone (*sensu* De Queiroz 2007) where distinctive characters are set up and gene flow are still possible. Not all investigated criteria can therefore distinguish them and the question of their validity as two distinct species arises. However, since they harbour distinct allelic states for the SNPs used, and since some SNPs are coding, differential characters between these GSHs are expected. The question is whether these characters allow to distinguish species (e.g., eye colour in humans is encoded by over 150 genes, resulting in many SNPs, and yet it is still a single species). Therefore, parsimoniously, these GSHs should be considered as a single species that potentially represents a species complex (e.g., *P. damicornis* with two GSHs or *P. acuta* with six GSHs), waiting for further (e.g., ecological or reproductive) evidences to separate them.

Interestingly, almost all 21 GSHs corresponded to previously defined genetic species hypotheses or clusters (based on the mtORF marker and microsatellites). Several GSHs had their own mtORF or PocHistone haplotypes, confirming that both can be used as diagnostic markers for some (but not all) *Pocillopora* species. Conversely, the over-partitioning previously found in several SSHs using microsatellites (e.g., Gélin et al. 2018a; Oury et al. 2021) was not retrieved. This could be an effect either of the limited numbers of loci in microsatellite inferences, or of genus-level phylogenetic inferences masking such genetic patterns. Genetic criteria therefore appear robust to define species limits but present a risk of overestimating their number. BFD*, as other molecular species delimitation methods, has already been suggested to overestimate the number of species (Grummer et al. 2014; Hundsdoerfer et al. 2019; Derkarabetian et al. 2022). This supports the need for integrative approaches, where molecular criteria should be the first criteria to robustly and objectively explore species limits and define genetic species hypotheses that are then confirmed with other criteria (as previously suggested by Pante et al. 2015). In particular, genomics, although not systematically necessary to molecularly identify species, appears fundamental to set robust species limits in such taxa whose phylogenetic reconstructions are complex. So for biodiversity monitoring (e.g., the global coral reef monitoring network), for each region, an exhaustive inventory of *Pocillopora* species using genomics/genetics is precognised to first identify the species present in the field.

### *From Multiple Criteria to Integrative Taxonomy: Towards a Revision of the* Pocillopora *Genus*

Putting together evidences from all approaches (i.e., genetics, morphology, geography and symbiosis ecology), 13 species appeared strongly supported, where only seven are currently recognised based on current taxonomy. Six species thus need formal taxonomic descriptions (Table 1). Clades 1 and 2 support current taxonomy, the first consisting of a single species (*P. effusa*, corresponding to GSH01), and the second being consistent with Schmidt-Roach et al. (2014) taxonomic revision [i.e., three species: *P. damicornis* (GSH04 *sensu lato), P. brevicornis* (GSH10) and *P. acuta* (GSH05 *sensu lato*)]. Further investigations are nevertheless needed to state whether *P. damicornis* and *P. acuta* represent both species complexes. Indeed, *P. damicornis* was separated into two GSHs and SSHs (04a and 04b) not supported by other criteria, but which could be ecologically distinct as previously suggested (Oury et al. 2020a). Similarly, *P. acuta* was partitioned into several GSHs and SSHs/clusters, either sympatric or allopatric, and some associated to distinct Symbiodiniaceae. Multiple genetic entities were previously delimited in this species (Gélin et al. 2017a, 2018b; Torres et al. 2020), questioning its monophyly. Clades 3 and 4 are less congruent with current taxonomy. First, within Clade 3, all five GSHs are strongly supported by all other criteria, but we were not able to rely them with a currently accepted species (one of them, probably GSH13c, should correspond to *P. verrucosa*, but the others seem to be new species). Then, within Clade 4, four species seemed strongly supported. GSH09a, GSH09b and GSH13b each correspond to distinct species (GSH09b most probably corresponding to *P. meandrina* Dana, 1846, while *P. villosa nomen nudem* was previously suggested for GSH13b; Gélin et al. 2017b). Only GSH09c_WIO_ and GSH09c_TSP_ could not be distinguished with certainty and are parsimoniously considered as different *P. grandis* lineages in allopatry for now, waiting further evidences.

In the light of these results, a new taxonomical revision of the *Pocillopora* genus, formally describing and naming these six new species (corresponding to GSH09a, GSH12, GSH13a, GSH13b, GSH14 and GSH15) becomes urgent. This will allow to throw off the multitude of current nomenclatures based on genetic lineages (Pinzón et al. 2013; Marti-Puig et al. 2014; Schmidt-Roach et al. 2014; Gélin et al. 2017b) which can be difficult to follow, even for specialists.

In conclusion, this study is the most extensive exploration to date of the taxonomy of the *Pocillopora* genus in terms of both genomic and geographic coverage. This genus represents a scleractinian taxon for which the definition of species limits has been challenging for decades. Several other criteria including morphology, biogeography or symbiosis ecology were also investigated to refine species limits and propose consensually and parsimoniously species hypotheses in the most integrative way possible. Some criteria appeared thus more informative than others, but all provided helpful insights for refining species limits. Here, we clearly delimited 21 genomic species hypotheses from 356 colonies sampled in three marine provinces (western Indian Ocean, tropical southwestern Pacific and south-east Polynesia), of which 13 species were strongly supported by all approaches and six appear to be new species. Importantly, we demonstrate once again the obsolescence of *corallum* macromorphology to identify most of the species. Conversely, micromorphological diagnostic characters and mtORF and PocHistone diagnostic haplotypes were highlighted for several species. Our recommendation is therefore to systematically identify *Pocillopora* species using these diagnostic criteria, prior to all types of studies involving the colonies (e.g., biodiversity, ecology, reproduction, adaptation, connectivity, exo- and endosymbiosis…) in order to reduce misidentifications. Finally, our results give new insights into the puzzle of defining *Pocillopora* species limits, supporting the existence of several new species. Next steps are to formally revise the taxonomy of the *Pocillopora* genus.

## Supporting information

Appendix 1: Sampling

Appendix 2: Molecular Analyses

Appendix 3: Morphological Analyses

Appendix 4: Symbiodiniaceae Analyses

Appendix 5: Illustration of the Pocillopora Genomic Species Hypotheses

## Acknowledgements

Coral sampling in New Caledonia was carried out during COBELO (http://dx.doi.org/10.17600/13100100), BIBELOT (http://dx.doi.org/10.17600/14003700), and CHEST (http://dx.doi.org/10.17600/15004500) oceanographic campaigns on board of RV Alis (IRD), and in the northeastern and northwestern of Madagascar during MAD (http://dx.doi.org/10.17600/16004700) oceanographic campaign on board of RV Antea (IRD). Sampling in Reunion Island was supported by program CONPOCINPA (LabEx CORAIL fund); in the south of Madagascar in collaboration with the Institut Halieutique des Sciences Marines (Tulear); and in Rodrigues Island with the collaboration of the Rodrigues Regional Assembly and the South-East Marine Protected Area supported by project Biodiversity (POCT FEDER fund); in Europa, Juan de Nova, and Glorioso Islands by program BIORECIE (financial supports from INEE, INSU, IRD, AAMP, FRB, TAAF, and the foundation Veolia Environnement); in Tromelin Island by program ORCIE (INEE), and in Mayotte by program SIREME (FED). HM thanks all the buddies who helped in photographs during diving (J. Butscher, S. Andréfouët, L. Bigot, and M. Pinault). We acknowledge the Plateforme Gentyane (microsatellite genotyping) of the Institut National de Recherche pour l’Agriculture, l’alimentation et l’Environnement (INRAE, Clermont-Ferrand, France), GenoScreen (Sanger sequencing; Lille, France), the Plateforme iGenSeq (library preparations and NGS sequencing) of the Institut du Cerveau et de la Moelle épinière (ICM, Paris, France), and G. Toutirais from the Plateau technique de Microscopie Électronique (scanning electron microscopy) of the Muséum National d’Histoire Naturelle (MNHN, Paris, France) for technical supports. We would also like to acknowledge S. Schmidt-Roach for his valuable advice on preparing SEM samples. Bioinformatics analyses were performed on the Genotoul bioinformatics platform Toulouse Occitanie (Bioinfo Genotoul, https://doi.org/10.15454/1.5572369328961167E12). NO was financially supported by a PhD contract from the Doctoral School “Sciences, Technologies, Santé” of Reunion Island University.

## Data Availability Statements

All data underlying this article are available online or upon reasonable request to the corresponding author. Raw sequencing reads (BioProject PRJNA831687; *Pocillopora* sequence capture: accession numbers SRR19052129-SRR19052500; Symbiodiniaceae ITS2 metabarcoding: accession numbers SRR19152377-SRR19152635) and new haplotype sequences (GenBank accession numbers ON155826-ON155833) were deposited on the NCBI. Microsatellite genotypes, morphometric data, reference sequences and SNP datasets were deposited on Dryad: https://doi.org/XXXXXXXXX.

## Author Contribution Statements

NO and HM designed the study. HM collected samples. NO and HM did lab steps. CN performed the bioinformatics for the ITS2 metabarcoding. NO performed all other bioinformatics and analysed the results with helpful guidance from SM and DA. NO wrote the original draft and all authors reviewed and edited the manuscript.

## Appendices

**Appendix 1** Sampling.

**Appendix 2** Molecular Analyses.

**Appendix 3** Morphological Analyses.

**Appendix 4** Symbiodiniaceae Analyses.

**Appendix 5** Illustration of the *Pocillopora* Genomic Species Hypotheses.

