## Appendix 1: Sampling for "From Genomics to Integrative Taxonomy? The Case Study of *Pocillopora* Corals"

### ***Sampling and Previous Identifications***

The sampling was the same as in Gélin et al. (2017). Briefly, ca. 9,000 *Pocillopora* colonies were sampled [branch tip + photographs except for Tromelin Island (Scattered Islands) due to field difficulties and the Society Islands as colonies were not collected for this purpose], independently of their *corallum* macromorphology, from March 2001 to October 2016, within three marine provinces: the western Indian Ocean (WIO), the tropical southwestern Pacific (TSP) and the south-east Polynesia (SEP), extended over six ecoregions (Spalding et al. 2007) and 18 islands/regions (Table S1; Fig. S1). In each locality, different habitats (reef slope, fringing reef, flat reef, or lagoon) were sampled, at various depths (from sea surface to 30 m depth), to maximise colonies genetic diversity. Fragments were preserved in 90% ethanol at room temperature, and deposited at Reunion Island University (Saint-Denis, La Réunion).

All colonies were also previously genotyped with 13 microsatellites (Table S2) and for a subset, we also sequenced the mitochondrial ORF locus (mtORF; see Gélin et al. 2017b for more details; Table S2). Each colony was thus assigned beforehand a primary and a secondary species hypothesis (PSH and SSH, respectively; *sensu* Gélin et al. 2017b), and a cluster when appropriate, based on these genetic data (see, for example, Oury et al. 2021). From now, to simplify the reading, PSHs that were not subdivided into several SSHs are designated SSHs, keeping their corresponding number (e.g.,

PSH01 switches to SSH01). These SSHs remain easily recognisable as no lowercase letter follows the number.

In this study, a subset of 356 *Pocillopora* colonies, covering the totality of the localities and morphotypes sampled, as well as all SSHs and clusters, was considered to maximise the genetic diversity explored (Table S1; Fig. S1). Additionally, four *Seriatopora hystrix* and four *Stylophora pistillata* colonies were sampled in Glorioso Islands (WIO) and Grande Terre (New Caledonia; TSP) to serve as outgroups in the phylogenomic analyses [both species are Pocilloporidae, diverging from the *Pocillopora* genus in the middle-end Paleogene (42.7-28.4 Ma; Simpson et al. 2011)].

**Table S1** Sampling localities of *Pocillopora* colonies.
*N*: total number of sampled *Pocillopora* colonies, $N_{mtORF}$ and $N_{Hist}$: numbers of sequences obtained for the mitochondrial open reading frame (mtORF) and the PocHistone loci, respectively, and $N_{ITS2}$: number of colonies high-throughput sequenced for the characterisation of Symbiodiniaceae communities.

| Province | Ecoregion | Island/Region | Code | Latitude | Longitude | *N* | $N_{mtORF}$ | $N_{Hist}$ | $N_{ITS2}$ |
|---|---|---|---|---|---|---|---|---|---|
| Western Indian Ocean (WIO) | Western and Northern Madagascar | Mayotte | MAY | -12.83131 | 45.16044 | 24 | 9 | 4 | 17 |
| | | Glorioso Islands | GLO | -11.56377 | 47.29394 | 10 | 10 | 2 | - |
| | | Juan de Nova Island | JDN | -17.04855 | 42.72176 | 20 | 10 | 4 | 17 |
| | | Europa Island | EUR | -22.36783 | 40.37185 | 13 | 8 | 3 | 13 |
| | | Northeastern Madagascar | MADne | -16.18321 | 49.94950 | 26 | 5 | 2 | 17 |
| | | Northwestern Madagascar | MADnw | -13.46366 | 48.25272 | 28 | 8 | 2 | 17 |
| | | Southwestern Madagascar | MADsw | -23.47539 | 43.66148 | 21 | 18 | 2 | 16 |
| | Mascarene Islands | Reunion Island | REU | -21.16115 | 55.57841 | 30 | 15 | 6 | 22 |
| | | Rodrigues Island | ROD | -19.69775 | 63.44172 | 19 | 9 | 1 | 15 |
| | Cargados Carajos/Tromelin Island | | TRO | -15.88083 | 54.52714 | 3 | 3 | - | 3 |
| Tropical Southwestern Pacific (TSP) | New Caledonia | Chesterfield Islands | CHE | -20.41574 | 158.80233 | 46 | 32 | 11 | 32 |
| | | Western Grande Terre | NCAw | -21.47567 | 165.57125 | 54 | 38 | 6 | 41 |
| | | Eastern Grande Terre | NCAe | | | 42 | 12 | - | 30 |
| | | Loyalty Islands | LOY | -20.96939 | 167.20426 | 7 | 5 | - | 7 |
| | Tonga Islands | | TON | -21.13061 | -175.22125 | 3 | 1 | - | - |
| South-East Polynesia (SEP) | Society Islands | Bora-Bora | BOR | -16.50025 | -151.73874 | 2 | 2 | - | 1 |
| | | Moorea | MOO | -17.52767 | -149.83867 | 7 | 6 | - | 7 |
| | | Tahiti | TAH | -17.65834 | -149.47704 | 1 | 1 | - | 1 |
| | | | | | **Total** | **356** | **192** | **43** | **256** |

**Table S2** Primers and PCR conditions used for the amplification of the mitochondrial open reading frame (mtORF), the PocHistone, the ribosomal RNA internal transcribed spacer 2 (ITS2) and the 13 microsatellite loci. *%NA*: percentage of missing data and *Na*: number of alleles (based on 356 *Pocillopora* colonies).

| Panel | Locus | Primers (5'-3') | Repeat motif | Dye | Size (bp) | *%NA* | *Na* | Reference |
|---|---|---|---|---|---|---|---|---|
| N/A | mtORF | FATP6.1: TTTGGGSATTCGTTTAGCAG<br>RORF: SCCAATATGTTAAACASCATGTCA | N/A | N/A | N/A | N/A | N/A | [1] |
| N/A | PocHistone | F: ATTCAGTCTCACTCACTCACTCAC<br>R: TATCTTCGAACAGACCCACCAAAT | N/A | N/A | N/A | N/A | N/A | [2] |
| N/A | ITS2 | itsD: GTGAATTGCAGAACTCCGTG<br>its2rev2: CCTCCGCTTACTTATATGCTT | N/A | N/A | N/A | N/A | N/A | [3]<br>[4] |
| 1 | Pd3-004 | F: M13-ACCAGACAGAAACACGCACA<br>R: GCAATGTGTAACAGAGGTGGAA | $(ATG)_8$ | 6-FAM | 172-202 | 11.4% | 11 | [5] |
| | Poc40 | F: M13-GTTATTATATGGGTGTATGC<br>R: CTCAAAGTGCGATTAAAGCC | $(CAA)_X$ | 6-FAM | 302-344 | 34.4% | 13 | [6] |
| | Pd3-005 | F: M13-AGAGTGTGGACAGCGAGGAT<br>R: GTTCCTTCGCCTTCGATTTT | $(TGA)_9$ | NED | 206-264 | 7.2% | 21 | [5] |
| | PV2 | F: M13-GCCAGGACCCATTTATACTCC<br>R: TGCAGTGTTCTACTTGTCAGTGC | $(GA)_{20}$ | VIC | 127-189 | 12.6% | 16 | [7] |
| | PV7 | F: M13-GGAGATGGATGGAGACTGC<br>R: GGTATCTCTGTGCTCAGTTCTTTG | $(GT)_5$ $(CT)_2$ GT $(CT)_3$ | VIC | 238-266 | 6.7% | 15 | [7] |
| 2 | Pd2-001 | F: M13-CAGACTTGTCGGAATGAAAGC<br>R: TTTTGTTTATAAGTCGATACAATGCA | $(CA)_{11}$ | VIC | 206-230 | 6.7% | 15 | [5] |
| | Pd2-006 | F: M13-ATCTCCATGTGATCGGCATT<br>R: GTTCCCCCAGCTGAGAAGTT | $(CA)_8$ | NED | 202-237 | 6.6% | 13 | [5] |
| | Pd3-008 | F: M13-AGTTGAGGTTGTTGAAACATG<br>R: TCCATGCAGAACCCC | $(CTG)_7$ | 6-FAM | 173-203 | 4.2% | 11 | [5] |
| | Pd3-009 | F: M13-CCAATGCGTCCGTAGCTCTC<br>R: ATCACCTAAAAATTTCAGTCCCTTACC | $(CAA)_7$ $(GAG)_6$ | 6-FAM | 334-369 | 25.8% | 12 | [5] |
| 3 | Pd3-EF65 | F: M13-TGTGCAGGTGTTGTGACTGA<br>R: TGTCTTTTTCACTTTTGCTTCAA | $(GTT)_5$ $(TGC)_{11}$ | PET | 194-242 | 12.5% | 17 | [8] |
| | Pd4 | F: M13-ACGCACACAAACCAACAAAC<br>R: TAATTCCATCAACTCAAAGGGG | $(AAAC)_5$ | 6-FAM | 142-184 | 13.3% | 21 | [9] |
| | Pd11 | F: M13-TCGTTTGAAGGGAAATGCTC<br>R: GGCATGCTATGTATGCGAGA | $(CA)_7$ T $(AC)_{13}$ | VIC | 144-184 | 28.5% | 20 | [9] |
| | Pd13 | F: M13-TGTTCCTCTCTTTCTCTCTTCCA<br>R: CATTTATGTTCCTTTCACGGC | $(TCTT)_5$ | NED | 146-236 | 28.9% | 22 | [9] |
| | | | | | **Total** | **15.3%** | **15.9 ± 1.1** | |

**mtORF/PocHistone:**

PCR mix (total volume = 25 µL): 1X of MasterMix (Applied Biosystems) + 0.3 µM of F primer + 0.3 µM of R primer + 2 ng.µL$^{-1}$ of genomic DNA.

Thermocycling program: 94°C/5 min + 40 × (94°C/60 s + 53°C/60 s + 72°C/60 s) + 72°C/5 min.

**ITS2:**

PCR mix (total volume = 20 µL): 1X of MasterMix (Applied Biosystems) + 0.3 µM of F primer + 0.3 µM of R primer + 2 ng.µL$^{-1}$ of genomic DNA.

Thermocycling program: 94°C/5 min + 35 × (94°C/45 s + 54°C/30 s + 72°C/60 s) + 72°C/5 min.

**Microsatellites:**

PCR mix (total volume = 10 µL): 1X of MasterMix (Applied Biosystems) + 0.025 µM of F primer (M13 tailed) + 0.25 µM of R primer + 0.25 µM of fluorescent dyed M13 tail + 2 ng.µL$^{-1}$ of genomic DNA.

Thermocycling program: 94°C/5 min + 7 × (94°C/30 s + 62°C [-1°C at each cycle]/30 s + 72°C/30 s) + 30 × (94°C/30 s + 55°C/30 s + 72°C/30 s) + 8 × (94°C/30 s + 56°C/30 s + 72°C/30 s) + 72°C/5 min

**Fig. S1** Sampling localities of *Pocillopora* colonies (number of sampled colonies in parentheses). MAY: Mayotte, GLO: Glorioso Islands, JDN: Juan de Nova Island, EUR: Europa Island, MADne: northeastern Madagascar, MADnw: northwestern Madagascar, MADsw: southwestern Madagascar, REU: Reunion Island, ROD: Rodrigues Island, TRO: Tromelin Island, CHE: Chesterfield Islands, NCA: Grande Terre (New Caledonia), LOY: Loyalty Islands (New Caledonia), TON: Tonga Islands, BOR: Bora-Bora, MOO: Moorea and TAH: Tahiti.

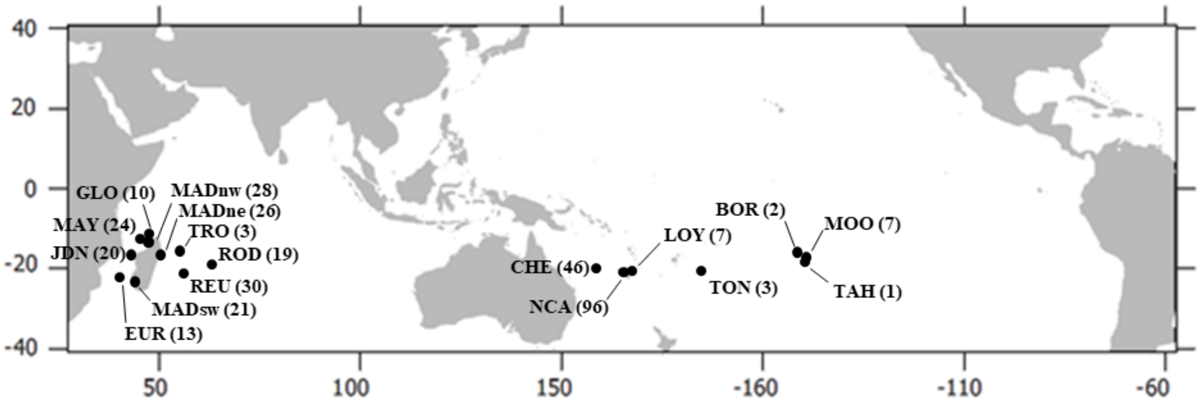
