## Appendix 2: Molecular Analyses for "From Genomics to Integrative Taxonomy? The Case Study of *Pocillopora* Corals"

#### *DNA Extraction, Library Preparation, and Sequencing*

Total genomic DNA was extracted using DNeasy® Blood & Tissue kit (QIAGEN GmbH, Hilden, Germany), following the manufacturer protocol. DNA quality was assessed using a NanoDrop™ 2000 spectrophotometer (Thermo Fisher Scientific, Wilmington, DE), while initial DNA concentration was measured with a Qubit® 2.0 fluorometer, using the Qubit® dsDNA BR Assay kit (Invitrogen, Carlsbad, CA). Library preparation with NEBNext® Ultra™ II FS DNA Library Prep kit for Illumina (New England Biolabs, Ipswich, MA) and sequencing were then performed at the platform iGenSeq (ICM, Paris, France). Briefly, ca. 100 ng of DNA was enzymatically fragmented to 200-450 bp, followed by end repair, A-tailing, and adaptor ligation according to the manufacturer's instructions. Then, adaptor-ligated fragments were size-selected (200-350 bp insert sizes), PCR-enriched, and indexed. PCR reactions were cleaned up and library quality was assessed on an Agilent 4200 TapeStation System (Agilent Technologies, Santa Clara, CA). Eight samples were independently prepared twice (sequencing replicates) to estimate the sequencing error rate, and the variant calling and filtering accuracy. Individual libraries were equimollary pooled per 32, and each resulting pool was enriched for a set of 1,248 ultraconserved elements (UCEs) and 1,385 exon loci originally designed from alignments of various anthozoan genomes and transcriptomes (Quattrini et al. 2018), using 17,302 synthetic RNA capture probes (myBaits®, Arbor Bioscences, Ann Arbor, MI). Target-enriched pools were then combined at equimolar ratios prior to PE150 sequencing with an Illumina NovaSeq 6000 (Illumina, San Diego, CA).

#### ***Reference Sequences Construction***

Following sequencing, the Illumina BaseSpace platform converted BCL data to fastq format and demultiplexed each sample according to individual-specific indexes (no mismatches allowed). All softwares and parameters used hereafter are summarised in Table S3.

Sequence reads from 50 individuals distributed among localities and morphotypes were used to *de novo* construct reference sequences for the targeted UCEs and exons. Reads were quality checked with FastQC v0.11.7 (http://www.bioinformatics.babraham.ac.uk/projects/fastqc/) and MultiQC v1.7 (Ewels et al. 2016), before and after adapter contamination and low-quality bases removal with cutadapt v2.1 (Martin 2011), available in the wrapper script Trim Galore! v0.6.0 (http://www.bioinformatics.babraham.ac.uk/projects/trim_galore/). Trimmed reads were *de novo* assembled using SPAdes v3.13.0 (Bankevich et al. 2012), and the resulting scaffolds were matched to the 17,302 RNA capture probes, using the *phyluce_assembly_match_contigs_to_probes* script from the PHYLUCE program (Faircloth 2016), to assign each scaffold to one of the 2,633 targeted loci. All scaffolds assigned to the same locus were then aligned using MAFFT v7.713 (Katoh and Standley 2013). A consensus was calculated from each sequence alignment with the cons program from the EMBOSS v6.5.7 suite (Rice et al. 2000) and constituted the reference sequence for the corresponding locus. When sequence alignments grouped highly divergent sequences (indicating potential paralogs or probes targeting several, more or less independent, regions), multiple alignments and consensus were calculated (this was the case for one UCE and 24 exons). This approach allowed to recover a reference sequence for 1,054 of the 1,248 targeted UCEs (84.5%) and 957 of the 1,385 targeted exons (69.1%).

Afterward, to retrieve additional loci, the trimmed reads from the 50 individuals were mapped to the already recovered locus reference sequences using a Burrows-Wheeler aligner (BWA v0.7.17; Li and Durbin 2009). Unmapped reads were filtered using samtools v1.9 (Li et al. 2009; http://www.htslib.org/), then mapped, along with the scaffolds, to the genome of *P. acuta* (Vidal-Dupiol et al. 2019). Genomic regions with a high read depth (> 30×) over 100 bp for any individual were identified using samtools and custom R v4.0.4 (R Core Team 2021) scripts, and the corresponding scaffolds were aligned as described above to calculate consensus reference sequences. This allows to retrieve 132 additional unidentified loci (named unk-001 to unk-132).

Finally, to identify overlaps or close reference sequences (indicating a physical linkage disequilibrium among loci), the sequences were mapped to the *P. acuta* genome, as previously for the scaffolds, and merged if necessary (gaps between sequences were filled using the *P. acuta* genome). Trimmed reads from the 50 individuals were mapped a last time to the reference sequences to perform a final visual check with IGV v2.8.0 (Robinson et al. 2011; Thorvaldsdóttir et al. 2013).

The resulting reference (available at https://XXXXXX) consists of 2,068 sequences with a mean length (± s.e.) of 863.2 ± 7.3 bp.

#### *Reads Mapping, SNP Calling, and Filtering*

Sequence reads were quality checked and trimmed as previously, then mapped to the reference sequences using BWA v0.7.17 (Li and Durbin 2009). Reads sorting and duplicates marking were performed with Picard v2.20.7 (https://broadinstitute.github.io/picard/), followed by local realignment with the genome analysis toolkit (GATK) v3.8.1 (McKenna et al. 2010), as in Van der Auwera et al. (2013; Table S3). The reference sequences coverage was estimated for each individual using samtools v1.9 (Li et al. 2009; http://www.htslib.org/), and loci presenting a mean read depth over all individuals < 10× or > 300× (potential paralogs) were discarded. The remaining 1,646 loci were manually investigated and those presenting a pattern compatible with duplicated elements (i.e., paralogs) were also removed to avoid introducing an artificial excess of heterozygous sites generated by the alignment of these divergent regions to the same reference sequence. Finally, 1,620 loci passed these preliminary filtering steps and were retained for subsequent analyses (Table S4).

BCFtools v1.9 (http://samtools.github.io/bcftools/) was used for single-nucleotide polymorphism (SNP) calling, treating all samples simultaneously. A base quality (BQ), a mapping quality (MQ) and a quality score (QUAL) of at least 20, 30 and 20, respectively, were required for calling a variant, while a minimum read depth (DP) of 12× and non-significant strand biases (SP ≤ 13) were required to call a genotype (Tables S3 & S4). From that, a total of 252,043 variants were called out, with a mean divergence (± s.e.) between sequencing replicates of the same individual of 0.16 ± 0.02% (Table S4).

Variants were then filtered using BCFtools and the R library '*vcfR*' (Knaus and Grünwald 2017; Table S4). Tri- and tetra-allelic sites, as well as sites presenting more than 20% of missing data or a minor allele frequency (MAF) inferior to 0.05 over all individuals were discarded. Three individuals, over the 364 sequenced (outgroups included and sequencing replicates excluded), presenting high proportions of missing data (> 60%) were also discarded at this step, leading to 17,465 SNPs, with a mean divergence (± s.e.) between sequencing replicates of 0.40 ± 0.05% (Table S4). Then, only one representative per replicate was kept (the one with the least missing data), leading to a final dataset of 361 individuals and 17,465 SNPs (Table S4). To reduce the effect of linkage disequilibrium, a second dataset was generated by choosing randomly one SNP per locus (361 individuals × 1,559 SNPs; Table S4).

***Phylogenomic Analyses***

All following analyses were performed on both datasets (i.e., with one or all SNPs per locus). Available *Pocillopora* genomes [i.e., *P. acuta* (Vidal-Dupiol et al. 2019), *P. damicornis* (Cunning et al. 2018) and *P. verrucosa* (Buitrago-López et al. 2020)] were also included. For these latter, the SNPs were retrieved by first mapping the reference sequences on each published genome, as previously. Genome regions where the reference sequences were mapped were then extracted using samtools and custom R v4.0.4 (R Core Team 2021) scripts and mapped in return to the reference sequences. SNP genotypes were then called for each genome separately using BCFtools (see Table S3 for the softwares and parameters used).

Filtered SNPs were first converted from VCF to PHYLIP and NEXUS formats using the python script *vcf2phylip* (https://github.com/edgardomortiz/vcf2phylip). Then, maximum-likelihood (ML) phylogenetic inferences were performed with RAxML-NG v0.9.0 (Kozlov et al. 2019). The phylogeny was constructed using the GTR+G model by searching for the ML tree, with 10 random and 10 parsimony-based starting trees and $10^3$ bootstrap replicates. Phylogenetic relationships were also investigated by means of Bayesian inference with BEAST v2.6.3 (Bouckaert et al. 2019), using the GTR+G model and an uncorrelated (lognormal) clock model. Chains were run for $5 \times 10^8$ iterations, sampling every $5 \times 10^4$ iterations, and checked for stationarity and parameter effective sample sizes with Tracer v1.7.1 (Rambaut et al. 2018). Final trees were built using the BEAST module TreeAnnotator v2.6.3 (Bouckaert et al. 2019), discarding the first 20% as burn-in, as indicated by Tracer, and visualised with FigTree v1.4.4 (https://github.com/rambaut/figtree).

To support the phylogenetic analyses and further explore the genetic partitioning of the dataset, several clustering approaches were used. First, assignment tests were performed with STRUCTURE v2.3.4 (Pritchard et al. 2000), sNMF (Frichot et al. 2014) and discriminant analyses of principal components (dAPC; Jombart et al. 2010). STRUCTURE was run with the admixture model, assuming correlated allele frequencies. Five iterations of $5 \times 10^5$ MCMC generations after an initial burn-in of $5 \times 10^4$ generations were run for each $K$, varying from $K = 2$ to $K = 30$. sNMF and dAPC were performed with the R libraries *'LEA'* (Frichot and François 2015) and '*adegenet'* (Jombart 2008), respectively. Five repetitions per $K$, with $K$ varying from 2 to 30, were run for sNMF, with a maximum of 500 iterations before reaching stationarity. Results were STRUCTURE–like plotted for all three assignment tests (i.e., STRUCTURE, sNMF and dAPC) with CLUMPAK (Kopelman et al. 2015), to allow their comparison. Signals of admixture between two clusters were further investigated with NEWHYBRIDS v1.1 (Anderson and Thompson 2002), parallelised with the R library *'parallelnewhybrid'* (Wringe et al. 2017). For each pair of clusters found admixed, NEWHYBRIDS was first run only with pure lineage individuals (i.e., assigned to one cluster or the other with a probability ≥ 0.9) to verify that the clusters were retrieved. Then, admixed individuals were included. One hundred runs, each with 100 randomly chosen SNPs, were performed with $10^6$ iterations after a burn-in period of $10^5$, and resulting assignment probabilities were averaged. Second, Nei (1972) individual genetic distances were computed with the R library *'StAMPP'* (Pembleton et al. 2013), and then used to build a minimum spanning tree (MST) and an unrooted equal-angle split network using EDENETWORKS v2.18 (Kivelä et al. 2015) and SplitsTree v4.15.1 (Huson and Bryant 2006), respectively.

Finally, SSH and cluster assignments issued from microsatellite data (Gélin et al. 2017a, 2017b, 2018a, 2018b; Oury et al. 2020, 2021, 2022) were compared to the groups identified with all above analyses, named hereafter genomic species hypotheses (GSHs). $F_{ST}$ (Weir and Cockerham 1984) were computed with *'StAMPP'* (Pembleton et al. 2013), for each pair of GSHs, and the resulting matrix was clustered using the *heatmap.2* function from the R library *'gplots'* (Warnes et al. 2020).

Besides, as some GSHs did not include any individual whose mtORF had previously been sequenced, and in order to retrieve the correspondence with previous studies, we completed the set of mtORF-sequenced colonies, following Gélin et al. (2017b), and further sequenced a subset of colonies for the PocHistone, a recently discovered marker partly mapped to partial histone 3 genes from other cnidarians, and allowing to identify *P. grandis* (the senior synonym of *P. eydouxi*) colonies (Johnston et al. 2018). The same laboratory protocol as for the mtORF was used, and sequencing in both directions was performed on an ABI 3730XL DNA Analyzer at GenoScreen (Lille, France). Sequences were checked and edited using GENEIOUS v8.1.9 (Kearse et al. 2012). mtORF and PocHistone haplotypes were identified by aligning them with those from Gélin et al. (2017b), and from Johnston et al. (2018), respectively (all sequences are available in GenBank). Alignments were performed with MAFFT v7.713 (Katoh and Standley 2013), and sequences were trimmed to the appropriate lengths. Phylogenetic relationships among PocHistone haplotypes were reconstructed in

MEGA v7.0.26 (Kumar et al. 2016), with the most appropriate substitution model according to jModelTest v2.1.10 (i.e., the JC+I model; Darriba et al. 2012).

***Species Delimitation Analyses and Divergence Time Estimation***

To confirm GSHs, Bayes factor delimitation with genomic data (BFD*; Leaché et al. 2014) was used to test several possible species delimitation models, using the SNAPP package (Bryant et al. 2012) implemented in BEAST v2.6.3 (Bouckaert et al. 2019). To deal with computationally intensive demands from SNAPP, we first tested a batch of species delimitation scenarios to confirm the four main clades from the phylogeny (arbitrarily defined as monophyletic groups of individuals separated by nucleotide substitution per site distances of at least 0.4 on the ML tree). Four scenarios were distinguished: (1) one single clade (Clades 1 + 2 + 3 + 4); (2) two clades (Clade 1 | Clades 2 + 3 + 4); (3) three clades (Clade 1 | Clades 2 + 3 | Clade 4) and (4) four clades (Clade 1 | Clade 2 | Clade 3 | Clade 4). Three individuals per clade were sampled, and BCFtools was used to generate the corresponding VCF file, including only unlinked biallelic SNPs from the dataset with one SNP per locus, which was then converted into binary NEXUS format using *vcf2phylip*. A path sampling method with 48 steps, each one consisting of $10^5$ MCMC generations, storing every $10^3$ generations with a pre-burn-in of $10^3$ steps, was run for each scenario, and the first 80% stored data was discarded. Model convergence was assessed with Tracer v1.7.1 (Rambaut et al. 2018) and models were compared and ranked using their marginal likelihood estimation (MLE) and by calculating the Bayes factor (BF; Kass and Raftery 1995). Once this first batch of scenarios tested, we retained the best-supported model, i.e. four clades (see Results). Then, several possible species delimitation models were tested within each clade separately, from one single species to the number of GSHs found for each clade (Table S5). SNAPP was run as above, and the best-supported model was retained for each clade.

For each of the best-supported model (except for Clade 1, constituted of a single GSH), a coalescent-based tree was calculated with SNAPP, keeping only two individuals per GSH to reduce the complexity in species tree estimation and increase the parameter convergence probability. Chains were run for $10^7$ iterations, sampling every $10^3$ iterations, and checked for convergence with Tracer v1.7.1 (Rambaut et al. 2018). Species trees that were contained in the 95% highest posterior density (HPD) set were then identified with SNAPP-TreeSetAnalyser v2.6.3 (Bryant et al. 2012), discarding the first 10% as burn-in, as suggested by Tracer. DensiTree v2.2.7 (Bouckaert 2010) was used to visualise the posterior distributions of topologies as cladograms, hence allowing for a clear depiction of uncertainty in the topology.

Finally, GSH divergence times were estimated with the BEAST package SNAPPER v1.0.1 (Stoltz et al. 2021). The latter is similar to SNAPP, except that it uses a diffusion model to calculate the likelihood of the species tree instead of mathematically integrating over all possible gene trees. Therefore, SNAPPER is far less computationally demanding than SNAPP and allowed us to include all delimited *Pocillopora* GSHs (each represented by three individuals) and outgroups (i.e., *S. hystrix* and *S. pistillata*; merged into one group) in a single analysis. The divergence between *Pocillopora* and outgroups was constrained to the middle-end Paleogene (42.7-28.4 Ma; Simpson et al. 2011) with a lognormal distribution (offset = 0, $\mu = 36$, $\sigma = 0.1$), and the most probable species tree identified with TreeSetAnalyser was used as starting tree. We used the ruby script *snapp_prep* (https://github.com/mmatschiner/snapp_prep) to prepare the xml input file, setting a chain of $10^7$ iterations with a sampling every $10^3$ iterations. Convergence and mixing were checked using Tracer v1.7.1 (Rambaut et al. 2018), and the maximum clade credibility tree with median node heights was generated with TreeAnnotator v2.6.3 (Bouckaert et al. 2019), discarding the first 10% as burn-in.

Van der Auwera G.A., Carneiro M.O., Hartl C., Poplin R., del Angel G., Levy-Moonshine A., Jordan T., Shakir K., Roazen D., Thibault J., Banks E., Garimella K.V., Altshuler D., Gabriel S., DePristo M.A. 2013. From FastQ data to high-confidence variant calls: the Genome Analysis Toolkit best practices pipeline. Curr. Protoc. Bioinforma. 43.

Vidal-Dupiol J., Chaparro C., Pratlong M., Pontaroti P., Grunau C., Mitta G. 2019. Sequencing, *de novo* assembly and annotation of the genome of the scleractinian coral, *Pocillopora acuta*. BioRxiv.

Warnes G.R., Bolker B., Bonebakker L., Gentleman R., Huber W., Liaw A., Lumley T., Maechler M., Magnusson A., Moeller S., Schwartz M., Venables B. 2020. gplots: various R programming tools for plotting data. R package version 3.1.1.

Weir B.S., Cockerham C.C. 1984. Estimating F-statistics for the analysis of population structure. Evolution. 38:1358–1370.

Wringe B.F., Stanley R.R.E., Jeffery N.W., Anderson E.C., Bradbury I.R. 2017. parallelnewhybrid: an R package for the parallelization of hybrid detection using newhybrids. Mol. Ecol. Resour. 17:91–95.

**Table S3** Softwares, tools, and parameters used for processing reads.

| Function | Tool | Software/package | Parameters | Reference |
|---|---|---|---|---|
| **Reference sequences construction** | | | | |
| Quality control | FastQC v0.11.7<br>MultiQC v1.7 | | N/A<br>N/A | [1]<br>[2] |
| Adapter trimming and low-quality bases removal | Trim Galore! v0.6.0 | | --paired R1.fq R2.fq<br>--trim-n<br>--illumina<br>--nextseq 20 | [3] |
| *De novo* reads assembly | SPAdes v3.13.0 | | -1 R1.fq -2 R2.fq<br>--careful<br>--cov-cutoff 10<br>-k 21,33,55,77 | [4] |
| Match scaffolds to probes | phyluce_assembly_match_contigs_to_probes | PHYLUCE | --contigs /path/to/scaffolds.fa<br>--probes hexa-v2-sclerac-subset-final-probes.fa<br>--regex='^((uce\|trans)-\d+)(?:_p\d+.*)' | [5] |
| Scaffolds alignment | MAFFT v7.713 | | --adjustdirection | [6] |
| Calculate consensus | cons | EMBOSS v6.5.7 | -sequence scaffolds_alignment.msa | [7] |
| Reads, scaffolds and sequences mapping | BWA v0.7.17 | | mem reference.fa R1.fq R2.fq<br>mem reference.fa scaffolds/sequences.fa | [8] |
| Unmapped reads filtering | view | samtools v1.9 | -f 4 | [9,10] |
| Calculate read depth | depth | samtools v1.9 | -m 50 | [9,10] |
| **Reads mapping, SNP calling, and filtering** | | | | |
| Quality control | FastQC v0.11.7<br>MultiQC v1.7 | | N/A<br>N/A | [1]<br>[2] |
| Adapter trimming and low-quality bases removal | Trim Galore! v0.6.0 | | --paired R1.fq R2.fq<br>--trim-n<br>--illumina<br>--nextseq 20 | [3] |
| Reads mapping | BWA v0.7.17 | | mem reference.fa R1.fq R2.fq | [8] |
| Reads sorting | SortSam | Picard v2.20.7 | SORT_ORDER=coordinate | [11] |
| Duplicates marking | MarkDuplicates | Picard v2.20.7 | ASO=coordinate | [11] |
| Local realignment | RealignerTargetCreator<br>IndelRealigner | GATK v3.8.1 | -R reference.fa<br>-R reference.fa | [12] |
| Calculate read depth | depth | samtools v1.9 | N/A | [9,10] |

| Function | Tool | Software/package | Parameters | Reference |
|---|---|---|---|---|
| **Reads mapping, SNP calling, and filtering** (continued) | | | | |
| SNP calling and filtering | mpileup | BCFtools v1.9 | -A -I -Q 20 -q 30 -a AD,DP,SP,INFO/AD<br>-f reference.fa | [13] |
| | call | | -mv | |
| | filter | | -i ‘QUAL>=20’<br>-S ‘.’ -e ‘FORMAT/DP<12’<br>-S ‘.’ -e ‘FORMAT/SP>13’<br>-m2 -M2 | |
| **Published genomes SNP genotyping** | | | | |
| References mapping | BWA v0.7.17 | | mem genome.fa reference.fa | [8] |
| Mapped regions extraction | view | samtools v1.9 | N/A | [9,10] |
| Genome mapping | BWA v0.7.17 | | mem reference.fa genome.fa | [8] |
| SNP genotyping | mpileup | BCFtools v1.9 | -A -B -I -a AD,DP,SP,INFO/AD -R SNP_positions.txt<br>-f reference.fa | [13] |
| | call | | -m | |

[10] http://www.htslib.org/

[11] https://broadinstitute.github.io/picard/

[12] McKenna A, Hanna M, Banks E, Sivachenko A, Cibulskis K, Kernytsky A, Garimella K, Altshuler D, Gabriel S, Daly M (2010) The Genome Analysis Toolkit: a MapReduce framework for analyzing next-generation DNA sequencing data. Genome Res 20:1297–1303. doi: 10.1101/gr.107524.110

[13] http://samtools.github.io/bcftools/

**Table S4** Single-nucleotide polymorphism (SNP) filtering steps. $N_{ind}$, $N_{loci}$, $N_{SNP}$ and $N_{GT}$: numbers of individuals, loci, SNPs and genotypes, respectively, *%NA*, $\%NA_{ind}$ and $\%NA_{SNP}$: percentages of missing data for the overall dataset, per individual and per SNP, respectively, *Δrep*: mean (± s.e.) divergence between sequencing replicates of the same individual (*N* = 8), MQ: mapping quality, BQ: base quality, $DP_{locus}$: mean locus depth of coverage overall individuals, QUAL: quality score, DP: genotype depth of coverage, SP: strand bias and MAF: minor allele frequency.

| **Filters** | $N_{ind}$ | $N_{loci}$ | $N_{SNP}$ | $N_{GT}$ | ***%NA*** | ***Δrep*** |
|---|---|---|---|---|---|---|
| MQ ≥ 30 & BQ ≥ 20 | 372 | 2 068 | 430 120 | 138 641 244 | 13.4% | 1.14 ± 0.07% |
| & 10 ≤ $DP_{locus}$ ≤ 300<br>& no paralogs | 372 | 1 620 | 370 498 | 122 436 721 | 11.2% | 0.94 ± 0.07% |
| & QUAL ≥ 20 | 372 | 1 620 | 347 685 | 116 530 354 | 9.9% | 0.97 ± 0.07% |
| & DP ≥ 12 | 372 | 1 620 | 254 575 | 74 756 826 | 21.1% | 0.19 ± 0.02% |
| & SP < 13 | 372 | 1 620 | 252 043 | 73 941 075 | 21.1% | 0.16 ± 0.02% |
| & no tri/tetra-allelic sites | 372 | 1 620 | 209 658 | 62 249 416 | 20.2% | 0.12 ± 0.01% |
| & $\%NA_{SNP}$ ≤ 20<br>& MAF ≥ 0.05 | 372 | 1 559 | 17 465 | 6 079 709 | 6.4% | 0.40 ± 0.05% |
| & $\%NA_{ind}$ ≤ 60 | 369 | 1 559 | 17 465 | 6 074 034 | 5.7% | 0.40 ± 0.05% |
| & no replicate | 361 | 1 559 | 17 465 | 5 939 477 | 5.8% | - |
| & 1 SNP/locus | 361 | 1 559 | 1 559 | 528 855 | 6.0% | - |

**Table S5** Bayes factor delimitation with genomic data (BFD*) results for each model for the first batch of scenarios, and then within each of the four clades (Clades 1-4) separately (scenarios are detailed below). The numbers of individuals (maximum three per clade/species) and SNPs (the remaining biallelic ones from the dataset with one SNP per locus) used for each batch of scenarios are indicated above. $N_{Species}$: numbers of species (or clades), MLE: marginal likelihood estimate, BF: Bayes factor, relative to the model with the lowest MLE (highlighted in grey) within each batch of scenarios.

| Model | $N_{Species}$ | MLE | BF | Rank |
|---|---|---|---|---|
| **Initial batch (12 individuals × 1 355 SNPs)** | | | | |
| (1) One clade (Clades 1 + 2 + 3 + 4) | 1 | -14221.08 | 4706.03 | 4 |
| (2) Two clades (Clade 1 \| Clades 2 + 3 + 4) | 2 | -13507.26 | 3278.39 | 3 |
| (3) Three clades (Clade 1 \| Clades 2 + 3 \| Clade 4) | 3 | -12515.50 | 1294.87 | 2 |
| (4) Four clades (Clade 1 \| Clade 2 \| Clade 3 \| Clade 4) | 4 | -11868.07 | - | 1 |
| **Clade 1 (5 individuals × 232 SNPs)** | | | | |
| (1.1) One species (Current taxonomy) | 1 | -1112.45 | - | 1 |
| (1.2) One species per marine province (WIO \| TSP) | 2 | -1381.70 | 538.49 | 2 |
| **Clade 2 (29 individuals × 876 SNPs)** | | | | |
| (2.1) One species (Previous taxonomy) | 1 | -25895.47 | 9880.77 | 17 |
| (2.2) One species per ocean (Indian \| Pacific) | 2 | -25385.23 | 8860.30 | 16 |
| (2.3) Three species (Current taxonomy) | 3 | -22502.79 | 3095.41 | 14 |
| (2.4) Second level of subclading on SNPs phylogenies | 3 | -23254.97 | 4599.78 | 15 |
| (2.5) Current taxonomy + first level of subclading on SNPs phylogenies | 4 | -22148.95 | 2387.73 | 11 |
| (2.6) Split per mtORF haplotype (ORF09 \| ORF17 \| ORF18-19 \| ORF30-31) | 4 | -22198.15 | 2486.13 | 12 |
| (2.7) Current taxonomy but splitting *P. acuta* per ocean | 4 | -22281.59 | 2653.02 | 13 |
| (2.8) Third level of subclading on SNPs phylogenies | 5 | -21726.92 | 1543.67 | 7 |
| (2.9) Current taxonomy but splitting *P. acuta* per ocean + first level of subclading on SNPs phylogenies | 5 | -22033.76 | 2157.35 | 10 |
| (2.10) Current taxonomy but splitting *P. acuta* according to fourth level of subclading on SNPs phylogenies | 6 | -21665.78 | 1421.39 | 6 |
| (2.11) Proposed SNPs phylogenies but lumping *P. damicornis* subclades + lumping *P. acuta* main clade by ocean | 6 | -21770.15 | 1630.14 | 9 |
| (2.12) Proposed SNPs phylogenies but lumping *P. brevicornis* subclades + lumping *P. acuta* main clade by ocean | 6 | -21728.71 | 1547.25 | 8 |
| (2.13) Gélin et al. (2017) + Oury et al. (2020) secondary species hypotheses (SSHs) | 7 | -21649.99 | 1389.81 | 5 |
| (2.14) Proposed SNPs phylogenies but lumping some *P. acuta* subclades | 7 | -21408.68 | 907.18 | 3 |
| (2.15) Proposed SNPs phylogenies but lumping *P. acuta* main clade by ocean | 7 | -21463.53 | 1016.90 | 4 |
| (2.16) Proposed SNPs phylogenies but lumping some *P. acuta* subclades | 8 | -21098.78 | 287.39 | 2 |
| (2.17) Proposed SNPs phylogenies | 10 | -20955.08 | - | 1 |

| Model | $N_{Species}$ | MLE | BF | Rank |
|---|---|---|---|---|
| **Clade 3 (15 individuals × 763 SNPs)** | | | | |
| (3.1) One species (Current taxonomy) | 1 | -11784.88 | 2509.84 | 8 |
| (3.2) One species per ocean (Indian \| Pacific) | 2 | -11331.36 | 1602.80 | 7 |
| (3.3) First level of subclading on SNPs phylogenies | 2 | -11255.65 | 1451.38 | 6 |
| (3.4) One species per marine province (WIO \| TSP \| SEP) | 3 | -10863.52 | 667.12 | 5 |
| (3.5) ~ Gélin et al. (2017) primary species hypotheses (PSHs) | 4 | -10765.61 | 471.30 | 4 |
| (3.6) Proposed SNPs phylogenies but lumping TSP subclades | 4 | -10729.69 | 399.46 | 3 |
| (3.7) Proposed SNPs phylogenies but lumping WIO subclades | 4 | -10664.78 | 269.65 | 2 |
| (3.8) Proposed SNPs phylogenies | 5 | -10529.96 | - | 1 |
| **Clade 4 (15 individuals × 958 SNPs)** | | | | |
| (4.1) One species (Current taxonomy) | 1 | -15006.26 | 4578.12 | 11 |
| (4.2) Two species (~ previous taxonomy) | 2 | -13622.59 | 1810.78 | 8 |
| (4.3) First level of subclading on SNPs phylogenies | 2 | -13540.03 | 1645.67 | 7 |
| (4.4) Split per mtORF haplotype (ORF46 \| ORF23-27) | 2 | -13937.59 | 2440.79 | 9 |
| (4.5) One species per ocean (Indian \| Pacific) | 2 | -14468.37 | 3502.34 | 10 |
| (4.6) Proposed SNPs phylogenies but lumping *P. grandis* and GSH09a | 3 | -13217.99 | 1001.58 | 5 |
| (4.7) First level of subclading, but splitting *P. grandis* and GSH09a | 3 | -13086.57 | 738.75 | 3 |
| (4.8) Previous taxonomy, but splitting GSH13b and *P. meandrina* | 3 | -13169.44 | 904.48 | 4 |
| (4.9) Previous taxonomy, but splitting *P. meandrina* per ocean | 3 | -13334.55 | 1234.71 | 6 |
| (4.10) Gélin et al. (2017) secondary species hypotheses (SSHs) | 4 | -12763.15 | 91.89 | 2 |
| (4.11) Proposed SNPs phylogenies | 5 | -12717.20 | - | 1 |

### Detail of the models:

#### 1) Initial batch

| Model | Clade 1 | Clade 2 | Clade 3 | Clade 4 |
|---|---|---|---|---|
| 1 | One species | | | |
| 2 | Clade 1 | Clade 2 + 3 + 4 | | |
| 3 | Clade 1 | Clade 2 + 3 | | Clade 4 |
| 4 | Clade 1 | Clade 2 | Clade 3 | Clade 4 |

#### 2) Clade 1

| Model | $01_{WIO}$ | $01_{TSP}$ |
|---|---|---|
| 1.1 | *P. effusa* | |
| 1.2 | $01_{WIO}$ | $01_{TSP}$ |

#### 3) Clade 2

| Model | 04a | 04b | 04c | 10 | 05c-1 | 05d | 05a-1 | 05c-2 | 05a-2 | 05a-3 |
|---|---|---|---|---|---|---|---|---|---|---|
| 2.1 | *P. damicornis sensu lato* | | | | | | | | | |
| 2.2 | TSP | | WIO | TSP | WIO | | TSP | WIO | TSP | |
| 2.3 | *P. damicornis* | | *P. brevicornis* | | *P.acuta* | | | | | |
| 2.4 | 04a-b-c + 10 | | | | 05a-1 + 05a-2 + 05c-1 + 05c-2 + 05d | | | | | 05a-3 |
| 2.5 | *P. damicornis* | | *P. brevicornis* | | 05a-1 + 05a-2 + 05c-1 + 05c-2 + 05d | | | | | 05a-3 |
| 2.6 | ORF09 | | ORF17 | 30-31 | ORF18-19 | | | | | |
| 2.7 | *P. damicornis* | | *P. brevicornis* | | WIO | | TSP | WIO | TSP | |
| 2.8 | 04a-b | | 04c + 10 | | 05a-1 + 05c-1 + 05d | | | 05a-2 + 05c-2 | | 05a-3 |
| 2.9 | *P. damicornis* | | *P. brevicornis* | | WIO | | TSP | WIO | TSP | 05a-3 |
| 2.10 | *P. damicornis* | | *P. brevicornis* | | 05c-1 | 05a-1 + 05d | | 05a-2 + 05c-2 | | 05a-3 |
| 2.11 | 04a | 04b | *P. brevicornis* | | 05c-1 | 05a-1 + 05d | | 05a-2 + 05c-2 | | 05a-3 |
| 2.12 | *P. damicornis* | | 04c | 10 | WIO | | TSP | WIO | TSP | 05a-3 |
| 2.13 | SSH04a | SSH04b | *P. brevicornis* | | SSH05c | SSH05d | SSH05a | SSH05c | SSH05a | SSH05a |
| 2.14 | 04a | 04b | *P. brevicornis* | | 05c-1 | 05a-1 + 05d | | 05a-2 + 05c-2 | | 05a-3 |
| 2.15 | 04a | 04b | 04c | 10 | WIO | | TSP | WIO | TSP | 05a-3 |
| 2.16 | 04a | 04b | 04c | 10 | 05c-1 | 05a-1 + 05d | | 05a-2 + 05c-2 | | 05a-3 |
| 2.17 | 04a | 04b | 04c | 10 | 05c-1 | 05d | 05a-1 | 05c-2 | 05a-2 | 05a-3 |

#### 4) Clade 3

| Model | 12 | 13a | 13c | 15 | 14 |
|---|---|---|---|---|---|
| 3.1 | *P. verrucosa* | | | | |
| 3.2 | WIO | | TSP + SEP | | |
| 3.3 | 12 + 13a-c + 15 | | | | 14 |
| 3.4 | WIO | | TSP | | SEP |
| 3.5 | PSH12 | PSH13 | | PSH15 | PSH14 |
| 3.6 | 12 | 13a | TSP | | 14 |
| 3.7 | WIO | | 13c | 15 | 14 |
| 3.8 | 12 | 13a | 13c | 15 | 14 |

#### 5) Clade 4

| Model | 13b | 09b | 09a | $09c_{WIO}$ | $09c_{TSP}$ |
|---|---|---|---|---|---|
| 4.1 | Clade 4 | | | | |
| 4.2 | ~ *P. meandrina* | | | *P. grandis* | |
| 4.3 | 13b + 09b | | 09a + 09c | | |
| 4.4 | ORF46 | ORF23 + ORF27 | | | |
| 4.5 | WIO | TSP | WIO | | TSP |
| 4.6 | 13b | 09b | 09a + 09c | | |
| 4.7 | 13b + 09b | | 09a | 09c | |
| 4.8 | 13b | *P. meandrina* | | *P. grandis* | |
| 4.9 | WIO | TSP | WIO | *P. grandis* | |
| 4.10 | SSH13b | SSH09b | SSH09a | SSH09c | |
| 4.11 | 13b | 09b | 09a | $09c_{WIO}$ | $09c_{TSP}$ |

**Table S6** Genomic species hypotheses (GSHs) geographical distribution. The number of colonies from each GSH and each locality is indicated, allowing to visualise sympatric and allopatric GSHs. WIO: western Indian Ocean (MAY: Mayotte, GLO: Glorioso Islands, JDN: Juan de Nova Island, EUR: Europa Island, MADne: northeastern Madagascar, MADnw: northwestern Madagascar, MADsw: southwestern Madagascar, REU: Reunion Island, ROD: Rodrigues Island and TRO: Tromelin Island); TSP: tropical southwestern Pacific [CHE: Chesterfield Islands, NCA: Grande Terre (New Caledonia), LOY: Loyalty Islands (New Caledonia) and TON: Tonga Islands]; SEP: south-east Polynesia (BOR: Bora-Bora, MOO: Moorea and TAH: Tahiti).

| Clade/GSH | | | WIO | | | | | | | | | | TSP | | | | | SEP | | |
|---|---|---|---|---|---|---|---|---|---|---|---|---|---|---|---|---|---|---|---|---|
| | | | MAY | GLO | JDN | EUR | MADne | MADnw | MADsw | REU | ROD | TRO | CHE | NCAw | NCAe | LOY | TON | BOR | MOO | TAH |
| 1 | | 01 | | | | | | | 1 | 1 | | | 5 | | | | | | | |
| 2 | | 04a | | | | | | | | | | | 6 | 7 | 3 | | | | | |
| | | 04b | | | | | | | | | | | | 3 | 1 | 2 | | | | |
| | | 04c | | | | | | | | | 2 | | | | | | | | | |
| | | 10 | | | | | | | | | | | | 6 | 4 | | | | | |
| | | 05c-1 | | | | | | | | 7 | 4 | | | | | | | | | |
| | | 05d | 8 | | 4 | | 8 | 5 | 4 | | | | | | | | | | | |
| | | 05a-1 | | | | | | | | | | | 4 | | | | | | | |
| | | 05c-2 | 1 | | 2 | | | 2 | 2 | 4 | 2 | | | | | | | | | |
| | | 05a-2 | | | | | | | | | | | 3 | 8 | | | | | | |
| | | 05a-3 | | | | | | | | | | | 1 | 6 | 4 | | | | | |
| 3 | | 12 | | | | | 3 | | | 1 | | 3 | | | | | | | | |
| | | 13a | 3 | 2 | 3 | 4 | 4 | 5 | 4 | 3 | 2 | | | | | | | | | |
| | | 13c | 1 | | | | | 1 | | | | | 8 | 12 | 11 | 1 | 3 | | | |
| | | 15 | | | | | | | | | | | | 3 | 1 | 1 | | | | |
| | | 14 | | | | | | | | | | | | | | | | 2 | 2 | 1 |
| 4 | | 13b | 2 | 2 | 2 | 2 | 3 | 3 | 2 | 2 | 2 | | | | | | | | | |
| | | 09b | | | | | | | | 2 | | | 6 | 4 | 7 | 3 | | | 4 | |
| | | 09a | 6 | 6 | 7 | 6 | 8 | 11 | 7 | 7 | 6 | | | | | | | | | |
| | | 09c$_{WIO}$ | 3 | | 2 | 1 | | | 1 | 2 | 2 | | | | | | | | | |
| | | 09c$_{TSP}$ | | | | | | | | | | | 12 | 6 | 9 | | | | | |

**Table S7** $F_{ST}$ (Weir and Cockerham 1984) between genomic species hypotheses (GSHs) for the datasets with one (lower triangle) or all (upper triangle) SNPs per locus. All $F_{ST}$ were significative ($P < 0.001$***). Group sizes are indicated in parentheses. *Ser*: *Seriatopora hystrix* and *Sty*: *Stylophora pistillata*.

| Group | *Ser* | *Sty* | 01 | 04a | 04b | 04c | 10 | 05c-1 | 05d | 05a-1 | 05c-2 | 05a-2 | 05a-3 | 12 | 13a | 13c | 15 | 14 | 13b | 09b | 09a | 09c WIO | 09c TSP |
|---|---|---|---|---|---|---|---|---|---|---|---|---|---|---|---|---|---|---|---|---|---|---|---|
| ***Ser* (4)** | -- | 0.788 | 0.805 | 0.781 | 0.849 | 0.853 | 0.861 | 0.686 | 0.613 | 0.665 | 0.688 | 0.699 | 0.757 | 0.710 | 0.664 | 0.614 | 0.785 | 0.738 | 0.669 | 0.644 | 0.615 | 0.652 | 0.629 |
| ***Sty* (4)** | 0.753 | -- | 0.777 | 0.768 | 0.835 | 0.834 | 0.852 | 0.665 | 0.593 | 0.642 | 0.664 | 0.680 | 0.735 | 0.683 | 0.643 | 0.594 | 0.767 | 0.706 | 0.649 | 0.624 | 0.602 | 0.632 | 0.616 |
| **01 (7)** | 0.846 | 0.802 | -- | 0.679 | 0.730 | 0.689 | 0.752 | 0.577 | 0.503 | 0.555 | 0.579 | 0.587 | 0.634 | 0.568 | 0.532 | 0.474 | 0.641 | 0.568 | 0.542 | 0.489 | 0.481 | 0.516 | 0.489 |
| **04a (16)** | 0.814 | 0.819 | 0.707 | -- | 0.279 | 0.340 | 0.580 | 0.455 | 0.388 | 0.423 | 0.455 | 0.448 | 0.429 | 0.575 | 0.560 | 0.499 | 0.621 | 0.588 | 0.617 | 0.594 | 0.585 | 0.623 | 0.600 |
| **04b (6)** | 0.860 | 0.866 | 0.757 | 0.314 | -- | 0.340 | 0.643 | 0.451 | 0.380 | 0.416 | 0.450 | 0.451 | 0.457 | 0.584 | 0.563 | 0.497 | 0.653 | 0.603 | 0.611 | 0.586 | 0.580 | 0.613 | 0.592 |
| **04c (2)** | 0.862 | 0.857 | 0.723 | 0.376 | 0.336 | -- | 0.366 | 0.316 | 0.258 | 0.270 | 0.310 | 0.313 | 0.311 | 0.488 | 0.501 | 0.431 | 0.576 | 0.508 | 0.553 | 0.526 | 0.534 | 0.533 | 0.535 |
| **10 (10)** | 0.896 | 0.901 | 0.795 | 0.635 | 0.689 | 0.432 | -- | 0.451 | 0.371 | 0.433 | 0.453 | 0.454 | 0.511 | 0.633 | 0.588 | 0.529 | 0.700 | 0.658 | 0.643 | 0.613 | 0.595 | 0.650 | 0.617 |
| **05c-1 (11)** | 0.715 | 0.690 | 0.598 | 0.501 | 0.477 | 0.339 | 0.493 | -- | 0.119 | 0.158 | 0.203 | 0.246 | 0.270 | 0.476 | 0.503 | 0.448 | 0.516 | 0.475 | 0.549 | 0.526 | 0.538 | 0.534 | 0.533 |
| **05d (30)** | 0.657 | 0.636 | 0.536 | 0.436 | 0.412 | 0.292 | 0.409 | 0.123 | -- | 0.112 | 0.173 | 0.211 | 0.220 | 0.433 | 0.468 | 0.420 | 0.456 | 0.426 | 0.515 | 0.499 | 0.523 | 0.502 | 0.508 |
| **05a-1 (11)** | 0.687 | 0.666 | 0.573 | 0.452 | 0.421 | 0.284 | 0.462 | 0.151 | 0.116 | -- | 0.219 | 0.222 | 0.240 | 0.450 | 0.485 | 0.427 | 0.487 | 0.449 | 0.534 | 0.513 | 0.528 | 0.516 | 0.522 |
| **05c-2 (12)** | 0.710 | 0.688 | 0.609 | 0.503 | 0.484 | 0.335 | 0.485 | 0.213 | 0.191 | 0.238 | -- | 0.123 | 0.211 | 0.479 | 0.500 | 0.447 | 0.519 | 0.475 | 0.550 | 0.529 | 0.538 | 0.536 | 0.535 |
| **05a-2 (12)** | 0.719 | 0.702 | 0.612 | 0.488 | 0.473 | 0.324 | 0.476 | 0.259 | 0.235 | 0.248 | 0.133 | -- | 0.206 | 0.483 | 0.502 | 0.447 | 0.523 | 0.483 | 0.553 | 0.530 | 0.538 | 0.539 | 0.537 |
| **05a-3 (4)** | 0.771 | 0.759 | 0.649 | 0.479 | 0.483 | 0.323 | 0.545 | 0.284 | 0.257 | 0.251 | 0.261 | 0.236 | -- | 0.465 | 0.486 | 0.420 | 0.532 | 0.471 | 0.544 | 0.520 | 0.527 | 0.526 | 0.528 |
| **12 (7)** | 0.768 | 0.748 | 0.619 | 0.614 | 0.619 | 0.525 | 0.680 | 0.524 | 0.485 | 0.492 | 0.531 | 0.531 | 0.495 | -- | 0.190 | 0.144 | 0.324 | 0.328 | 0.478 | 0.473 | 0.492 | 0.486 | 0.486 |
| **13a (30)** | 0.730 | 0.723 | 0.573 | 0.595 | 0.594 | 0.529 | 0.627 | 0.546 | 0.515 | 0.523 | 0.553 | 0.551 | 0.514 | 0.224 | -- | 0.209 | 0.333 | 0.346 | 0.490 | 0.493 | 0.510 | 0.511 | 0.505 |
| **13c (37)** | 0.680 | 0.657 | 0.521 | 0.544 | 0.536 | 0.469 | 0.571 | 0.493 | 0.473 | 0.471 | 0.506 | 0.502 | 0.448 | 0.182 | 0.226 | -- | 0.172 | 0.248 | 0.458 | 0.455 | 0.485 | 0.467 | 0.467 |
| **15 (5)** | 0.840 | 0.819 | 0.687 | 0.647 | 0.675 | 0.598 | 0.739 | 0.538 | 0.493 | 0.508 | 0.555 | 0.552 | 0.538 | 0.360 | 0.345 | 0.183 | -- | 0.371 | 0.528 | 0.512 | 0.516 | 0.526 | 0.517 |
| **14 (5)** | 0.781 | 0.753 | 0.626 | 0.630 | 0.642 | 0.555 | 0.710 | 0.504 | 0.467 | 0.475 | 0.524 | 0.521 | 0.498 | 0.380 | 0.378 | 0.271 | 0.397 | -- | 0.473 | 0.444 | 0.467 | 0.458 | 0.459 |
| **13b (20)** | 0.718 | 0.700 | 0.585 | 0.654 | 0.650 | 0.598 | 0.686 | 0.588 | 0.560 | 0.567 | 0.595 | 0.593 | 0.575 | 0.542 | 0.546 | 0.521 | 0.583 | 0.524 | -- | 0.267 | 0.317 | 0.369 | 0.378 |
| **09b (26)** | 0.699 | 0.690 | 0.545 | 0.632 | 0.627 | 0.573 | 0.659 | 0.568 | 0.546 | 0.551 | 0.576 | 0.575 | 0.555 | 0.535 | 0.549 | 0.520 | 0.564 | 0.501 | 0.296 | -- | 0.252 | 0.317 | 0.315 |
| **09a (64)** | 0.651 | 0.650 | 0.509 | 0.615 | 0.612 | 0.567 | 0.626 | 0.572 | 0.562 | 0.557 | 0.574 | 0.572 | 0.552 | 0.534 | 0.553 | 0.535 | 0.556 | 0.507 | 0.360 | 0.287 | -- | 0.223 | 0.276 |
| **09cWIO (11)** | 0.694 | 0.692 | 0.564 | 0.661 | 0.650 | 0.571 | 0.690 | 0.573 | 0.548 | 0.552 | 0.581 | 0.579 | 0.558 | 0.550 | 0.581 | 0.538 | 0.583 | 0.522 | 0.440 | 0.391 | 0.260 | -- | 0.092 |
| **09cTSP (27)** | 0.672 | 0.678 | 0.541 | 0.639 | 0.632 | 0.572 | 0.652 | 0.576 | 0.558 | 0.562 | 0.584 | 0.582 | 0.564 | 0.549 | 0.572 | 0.539 | 0.571 | 0.520 | 0.438 | 0.386 | 0.326 | 0.107 | -- |

**Table S8** Number of *Pocillopora* colonies per mitochondrial open reading frame (mtORF) haplotype and locality. *N*: total number of sampled colonies.

| Province | Ecoregion | Locality | Code | *N* | ORF.. | 01 | 09 | 17 | 18 | 19 | 23 | 27 | 30 | 31 | 34 | 35 | 36 | 38 | 39 | 42 | 43 | 46 | 47 | 49 | 50 | 52 | 53 | 54 | TOTAL |
|---|---|---|---|---|---|---|---|---|---|---|---|---|---|---|---|---|---|---|---|---|---|---|---|---|---|---|---|---|---|
| Western Indian Ocean (WIO) | Western and Northern Madagascar | Mayotte | MAY | 24 | | | | | 2 | | | 3 | | | | | 1 | | | | | 2 | 1 | | | | | | **9** |
| | | Glorioso Islands | GLO | 10 | | | | | | | | 6 | | | | | 2 | | | | | 2 | | | | | | | **10** |
| | | Juan de Nova Island | JDN | 20 | | | | | 2 | | | 4 | | | | | | | 2 | | | 2 | | | | | | | **10** |
| | | Europa | EUR | 13 | | | | | | | | 5 | | | | | | 1 | | | | 2 | | | | | | | **8** |
| | | Madagascar | MADne | 26 | | | | | | | | | | | 3 | | | | 2 | | | | | | | | | | **5** |
| | | | MADnw | 28 | | | | | 1 | | | 2 | | | | | 1 | | 3 | | | | | | | | 1 | | **8** |
| | | | MADsw | 21 | | 1 | | | 3 | 2 | | 6 | | | | | | | 4 | | | 2 | | | | | | | **18** |
| | Mascarene Islands | Reunion | REU | 30 | | 1 | | | 6 | | | 4 | | | 1 | | | 1 | 1 | | 1 | | | | | | | | **15** |
| | | Rodrigues | ROD | 19 | | | | 2 | 3 | | | 1 | | | | | | | 2 | | | 1 | | | | | | | **9** |
| | Cargados Carajos/Tromelin | | TRO | 3 | | | | | | | | | | | 3 | | | | | | | | | | | | | | **3** |
| Tropical Southwestern Pacific (TSP) | New Caledonia | Chesterfield Islands | CHE | 46 | | 5 | 4 | | 6 | | 3 | 7 | | | | | | | | | | | 4 | | | | 3 | | **32** |
| | | Grande Terre | NCAw | 54 | | | 7 | | 9 | | 1 | 6 | 3 | 3 | | | | | | | | | 2 | | | 2 | 3 | 2 | **38** |
| | | | NCAe | 42 | | | | | 1 | | 2 | 2 | | 1 | | 1 | | | | | | | 2 | | | 1 | | 2 | **12** |
| | | Loyalty Islands | LOY | 7 | | | 2 | | | | 1 | 1 | | | | | | | | 1 | | | | | | | | | **5** |
| | Tonga Islands | Tonga Islands | TON | 3 | | | | | | | | | | | | | | | | | | | | | | | 1 | | **1** |
| South-East Polynesia (SEP) | Society Islands | Bora-Bora | BOR | 2 | | | | | | | | | | | | | | | | | | | | | 2 | | | | **2** |
| | | Moorea | MOO | 7 | | | | | | | 1 | 2 | | | | | | | | | | | | 1 | 2 | | | | **6** |
| | | Tahiti | TAH | 1 | | | | | | | | | | | | | | | | | | | | | 1 | | | | **1** |
| | | | **TOTAL** | **356** | | **7** | **13** | **2** | **33** | **2** | **8** | **49** | **3** | **4** | **7** | **1** | **4** | **2** | **14** | **1** | **1** | **11** | **9** | **1** | **5** | **3** | **8** | **4** | **192** |

**Table S9** PocHistone haplotype diversity. (a) number of *Pocillopora* colonies per PocHistone haplotype and locality (*N*: total number of sampled colonies) and (b) haplotype divergent bases [numbered from the 588 bp-alignment; position 222 (in grey) contain the *P. grandis* diagnostic thymine; *N*: haplotype occurrence in this study].

(a)

| Province | Ecoregion | Locality | Code | *N* | Hist.. | 01 | 02 | 03 | 04 | 05 | 06 | 07 | 08 | TOTAL |
|---|---|---|---|---|---|---|---|---|---|---|---|---|---|---|
| Western Indian Ocean (WIO) | Western and Northern Madagascar | Mayotte | MAY | 24 | | | | 1 | | | 1 | 2 | | **4** |
| | | Glorioso Islands | GLO | 10 | | | 1 | | | | 1 | | | **2** |
| | | Juan de Nova Island | JDN | 20 | | 1 | | | | | 1 | 2 | | **4** |
| | | Europa | EUR | 13 | | 1 | | | | | 1 | 1 | | **3** |
| | | Madagascar | MADne | 26 | | 1 | | | | | 1 | | | **2** |
| | | | MADnw | 28 | | | 1 | | | | 1 | | | **2** |
| | | | MADsw | 21 | | | | | 1 | | 1 | | | **2** |
| | Mascarene Islands | Reunion | REU | 30 | | 1 | | 1 | | | 2 | 1 | 1 | **6** |
| | | Rodrigues | ROD | 19 | | | | | | | 1 | | | **1** |
| | Cargados Carajos/Tromelin | | TRO | 3 | | | | | | | | | | **-** |
| Tropical Southwestern Pacific (TSP) | New Caledonia | Chesterfield Islands | CHE | 46 | | | | | | 5 | | 6 | | **11** |
| | | Grande Terre | NCAw | 54 | | | | | | 2 | | 4 | | **6** |
| | | | NCAe | 42 | | | | | | | | | | **-** |
| | | Loyalty Islands | LOY | 7 | | | | | | | | | | **-** |
| | Tonga Islands | | TON | 3 | | | | | | | | | | **-** |
| South-East Polynesia (SEP) | Society Islands | Bora-Bora | BOR | 2 | | | | | | | | | | **-** |
| | | Moorea | MOO | 7 | | | | | | | | | | **-** |
| | | Tahiti | TAH | 1 | | | | | | | | | | **-** |
| | | | **TOTAL** | **356** | | **4** | **2** | **2** | **1** | **7** | **10** | **16** | **1** | **43** |

(b)

| **Haplotype** | ***N*** | **Position (in bp)** 9 | **41** | **100** | **114** | **156** | **202** | **222** | **235** | **237** | **376** | **490** | **525** | **580** |
|---|---|---|---|---|---|---|---|---|---|---|---|---|---|---|
| Hist01 (GSH13b) | 4 | C | C | C | A | C | A | G | A | A | T | G | A | T |
| Hist02 (GSH13b) | 2 | C | C | C | G | C | A | G | A | A | T | G | A | T |
| Hist03 (GSH13b) | 2 | G | C | C | A | C | C | G | T | A | T | A | A | T |
| Hist04 (GSH13b) | 1 | G | C | A | A | T | C | G | A | A | T | G | A | T |
| Hist05 (GSH09b) | 7 | N | C | C | A | C | C | G | A | A | T | G | A | T |
| Hist06 (GSH09a) | 10 | C | G | C | A | C | A | G | A | A | T | G | A | T |
| Hist07 (GSH09c$_{WIO + TSP}$) | 16 | G | C | C | A | C | C | **T** | A | C | T | G | A | T |
| Hist08 (GSH09c$_{WIO}$) | 1 | G | C | C | A | C | C | **T** | A | C | C | G | A | T |
| MG587096 (*P. grandis*) | - | G | C | C | A | C | C | **T** | A | C | T | G | C | C |
| MG587097 (*P. meandrina*) | - | C | C | C | A | C | A | G | A | A | T | G | A | C |

**Fig. S2** *Pocillopora* maximum likelihood (ML) phylogenetic tree reconstructed with all SNPs (361 individuals × 17,465 SNPs). Branches are coloured according to marine provinces [black: western Indian Ocean (WIO); light grey: tropical southwestern Pacific (TSP); dark gray: south-east Polynesia (SEP)], and branch support, based on ML bootstrap analyses (first number) and Bayesian posterior probabilities (second number), is indicated for branches supporting the genomic species hypotheses (GSHs; delimited by dashed lines; full lines delimit the clades indicated alongside). Published genomes (in grey) are indicated by lowercase letters [a: *P. verrucosa* (Buitrago-López et al. 2020); b: *P. acuta* (Vidal-Dupiol et al. 2019); c: *P. damicornis* (Cunning et al. 2018)]. Clades and GSHs (number of colonies in parentheses), as well as corresponding colour code, are also indicated.

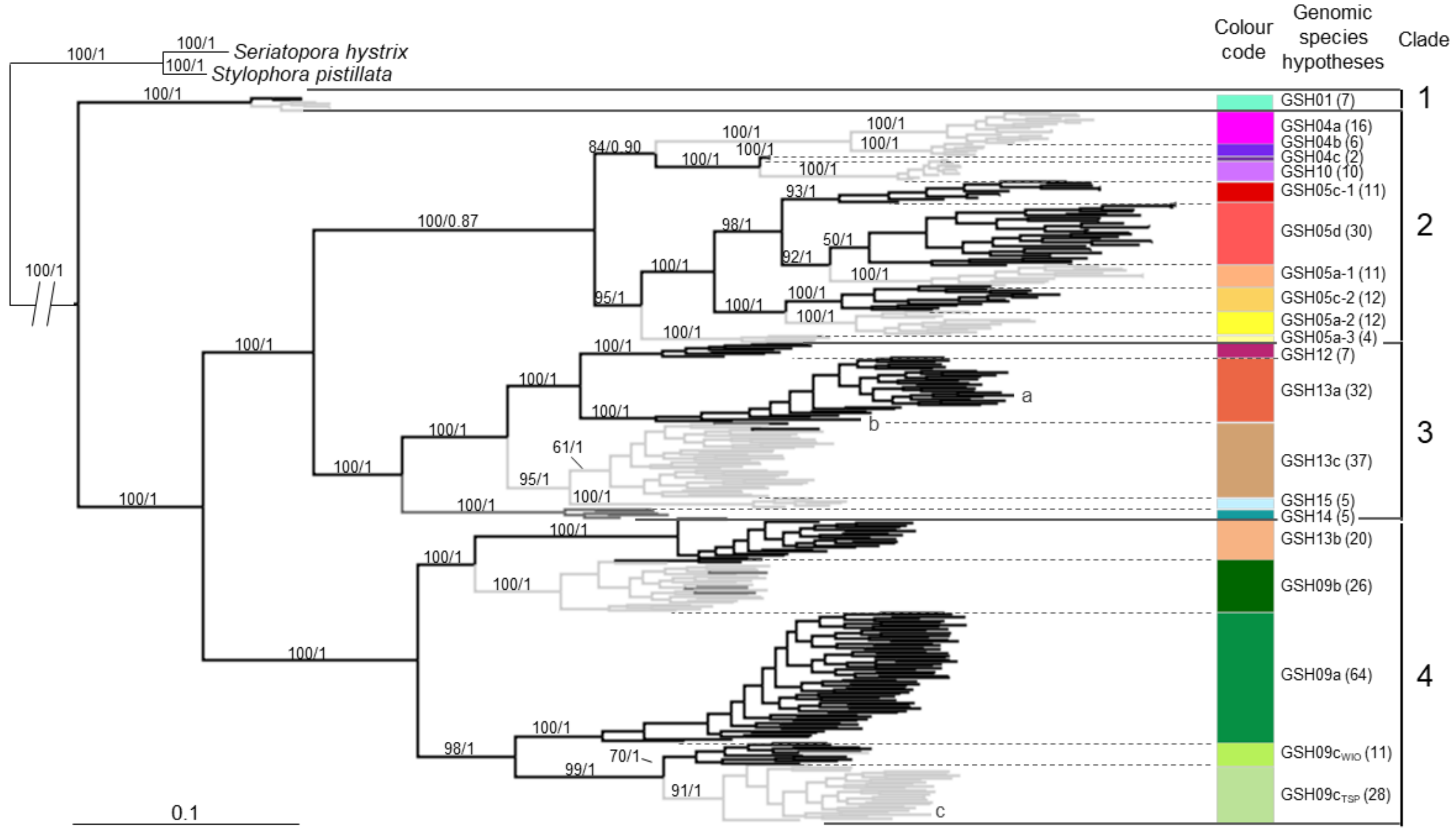

**Fig. S3** Assignment tests with one SNP per locus (361 individuals × 1,559 SNPs). (a) cross-entropy (sNMF), (b) mean likelihood over the five iterations of the same *K* (STRUCTURE), (c) Bayesian information criterion (BIC; dAPC) and (d) plots of the three assignment methods (i.e., sNMF, STRUCTURE and dAPC) from *K* = 2 to *K* = 30, with individuals grouped by genomic species hypothesis (GSH; indicated above). Assignments for the three published genomes [a: *P. verrucosa* (Buitrago-López et al. 2020); b: *P. acuta* (Vidal-Dupiol et al. 2019); c: *P. damicornis* (Cunning et al. 2018)] are detailed on the right panel.

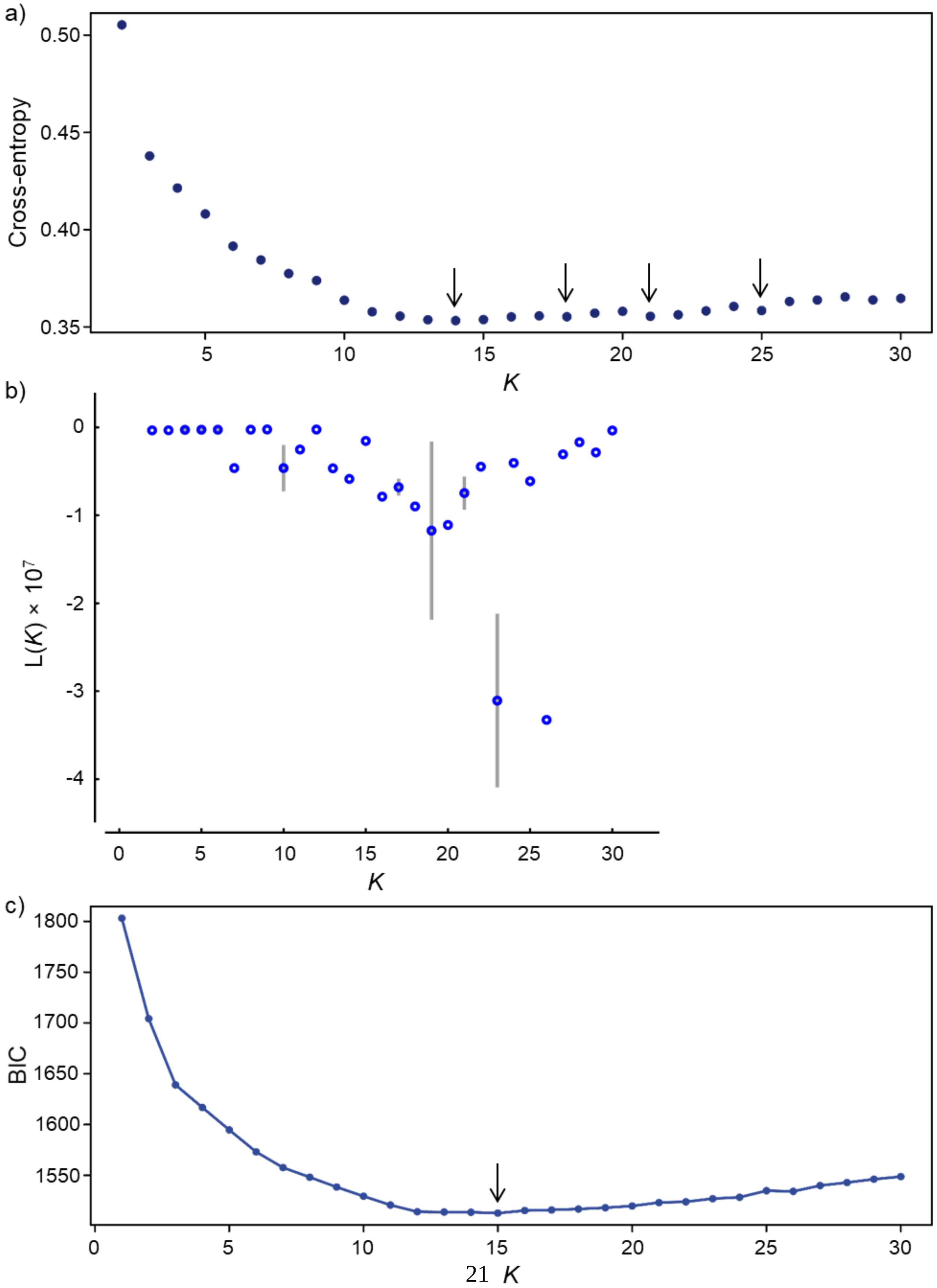

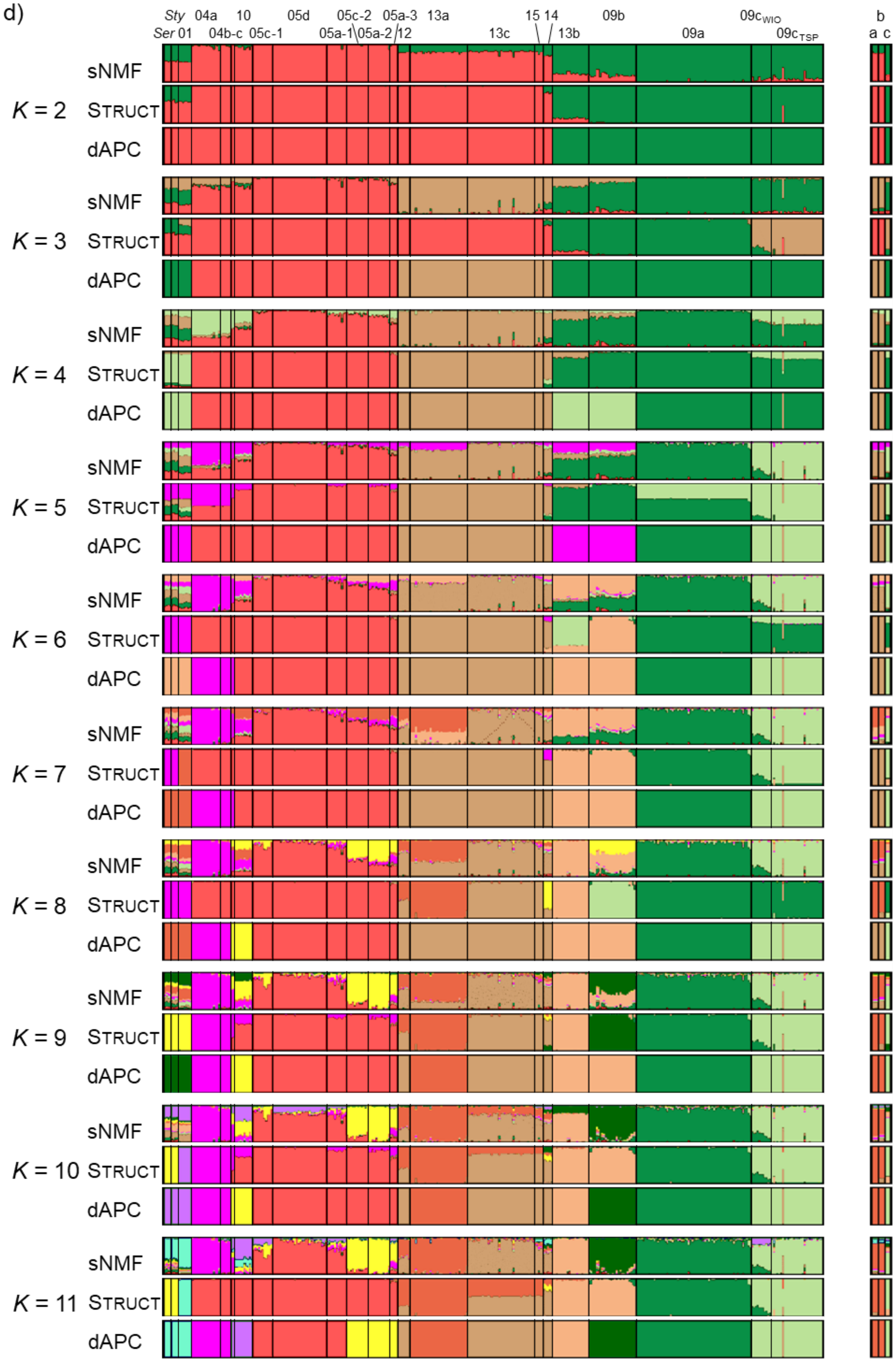
d)
Sty
Ser
01
04a
04b-c
10
05c-1
05d
05c-2
05a-1
05a-2
05a-3
12
13a
13c
15
14
13b
09b
09a
$09c_{WIO}$
$09c_{TSP}$
b
a
c
sNMF
STRUCT
dAPC
K = 2
K = 3
K = 4
K = 5
K = 6
K = 7
K = 8
K = 9
K = 10
K = 11

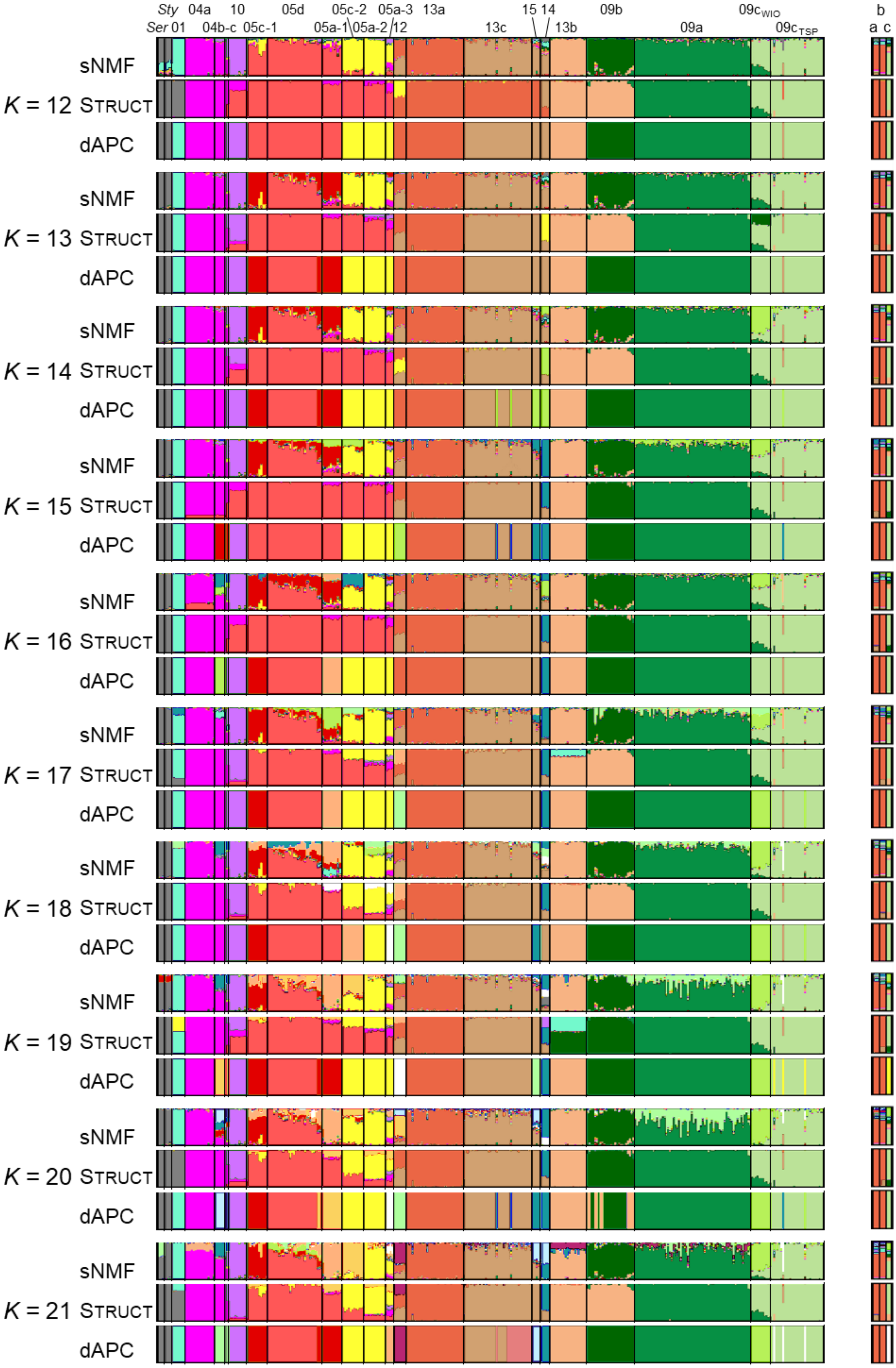
Sty
Ser
01
04a
04b-c
10
05c-1
05d
05c-2
05a-1
05a-2
05a-3
12
13a
13c
15
14
13b
09b
09a
09cWIO
09cTSP
b
a
c
sNMF
K = 12 STRUCT
dAPC
K = 13
K = 14
K = 15
K = 16
K = 17
K = 18
K = 19
K = 20
K = 21

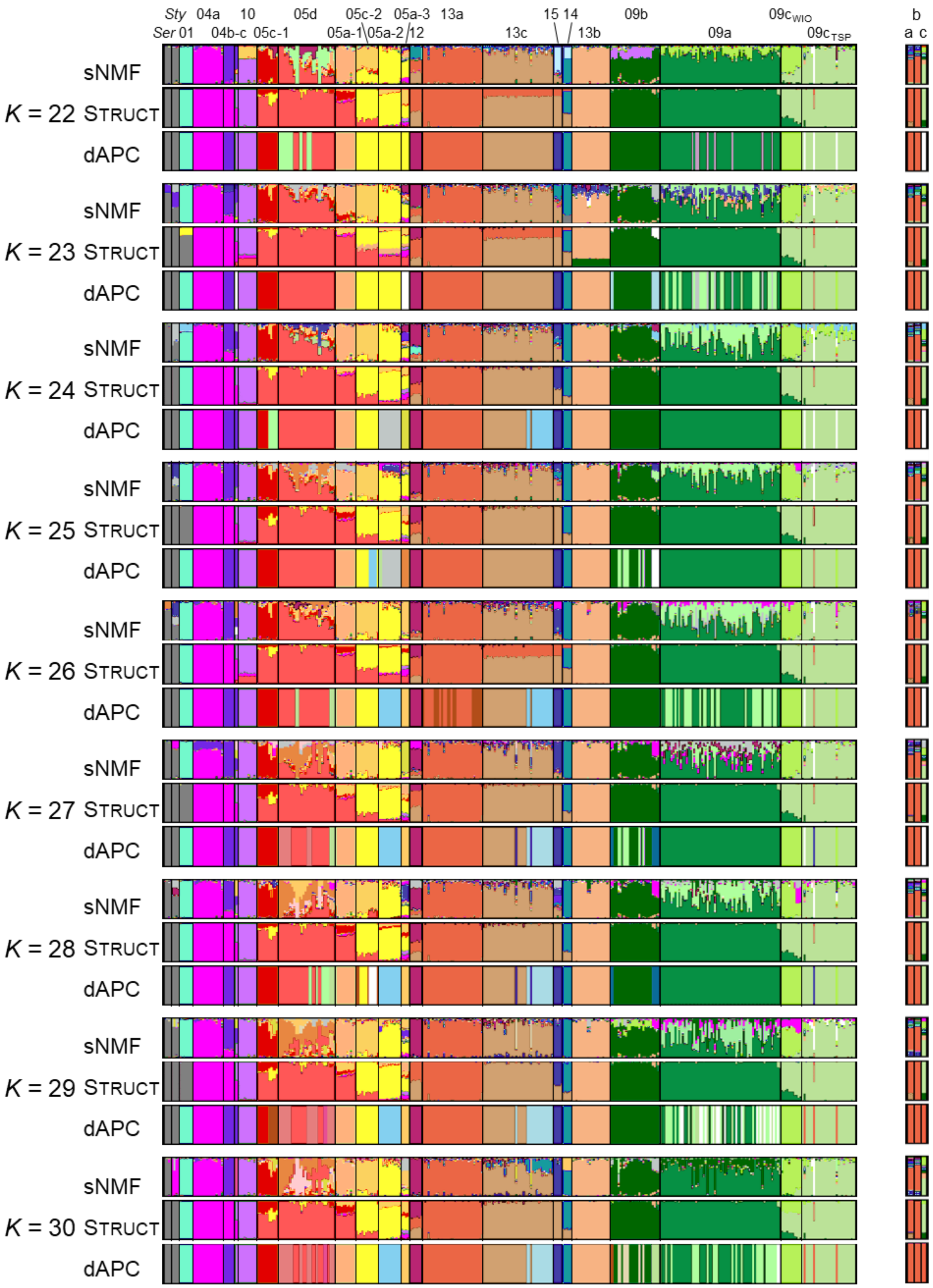

**Fig. S4** Assignment tests with all SNPs (361 individuals × 17,465 SNPs). (a) cross-entropy (sNMF), (b) Bayesian information criterion (BIC; dAPC) and (c) plots of the two assignment methods (i.e., sNMF and dAPC) from $K$ = 2 to $K$ = 30, with individuals grouped by genomic species hypothesis (GSH; indicated above). Assignments for the three published genomes [a: *P. verrucosa* (Buitrago-López et al. 2020); b: *P. acuta* (Vidal-Dupiol et al. 2019); c: *P. damicornis* (Cunning et al. 2018)] are detailed on the right panel.

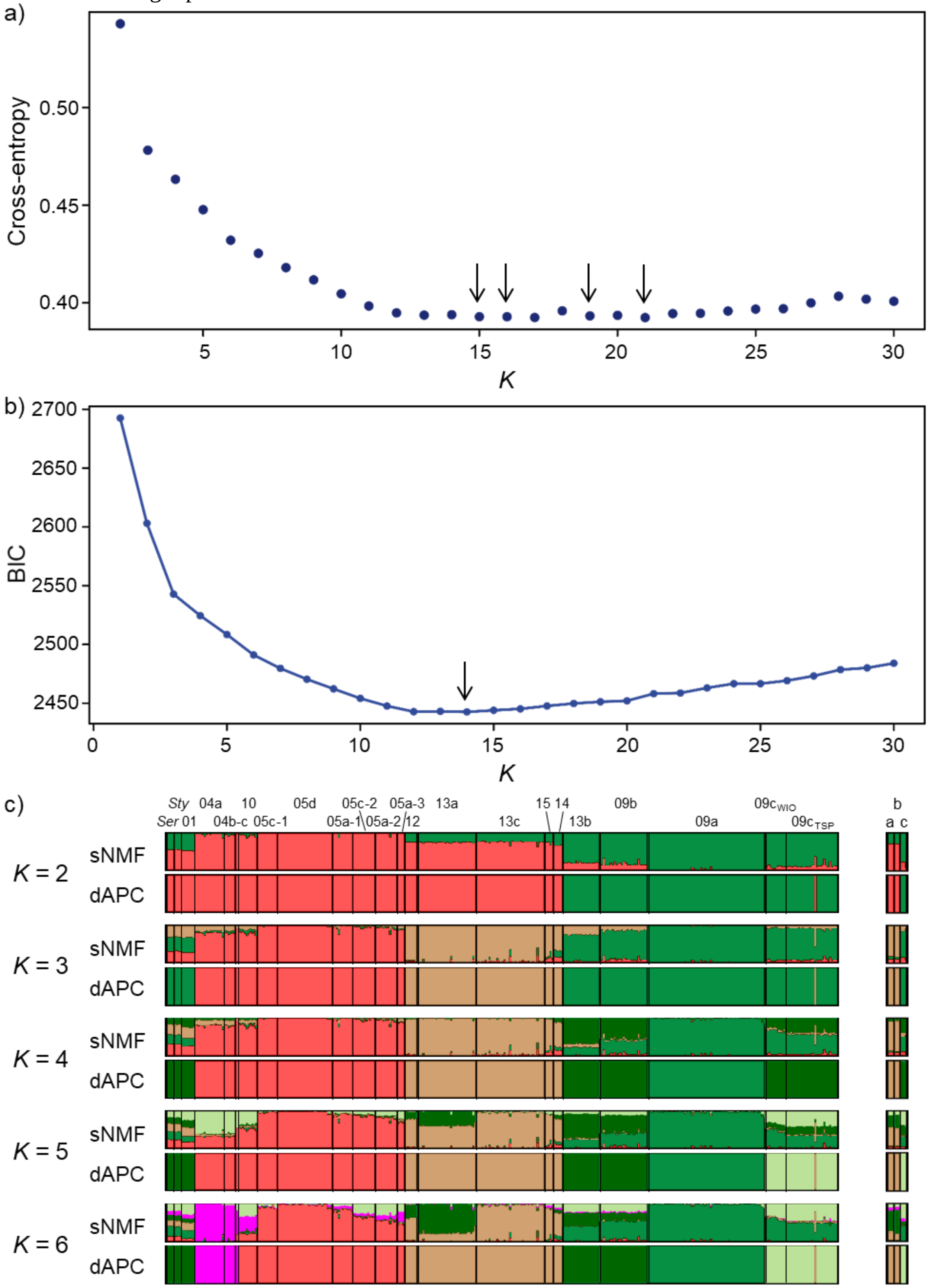

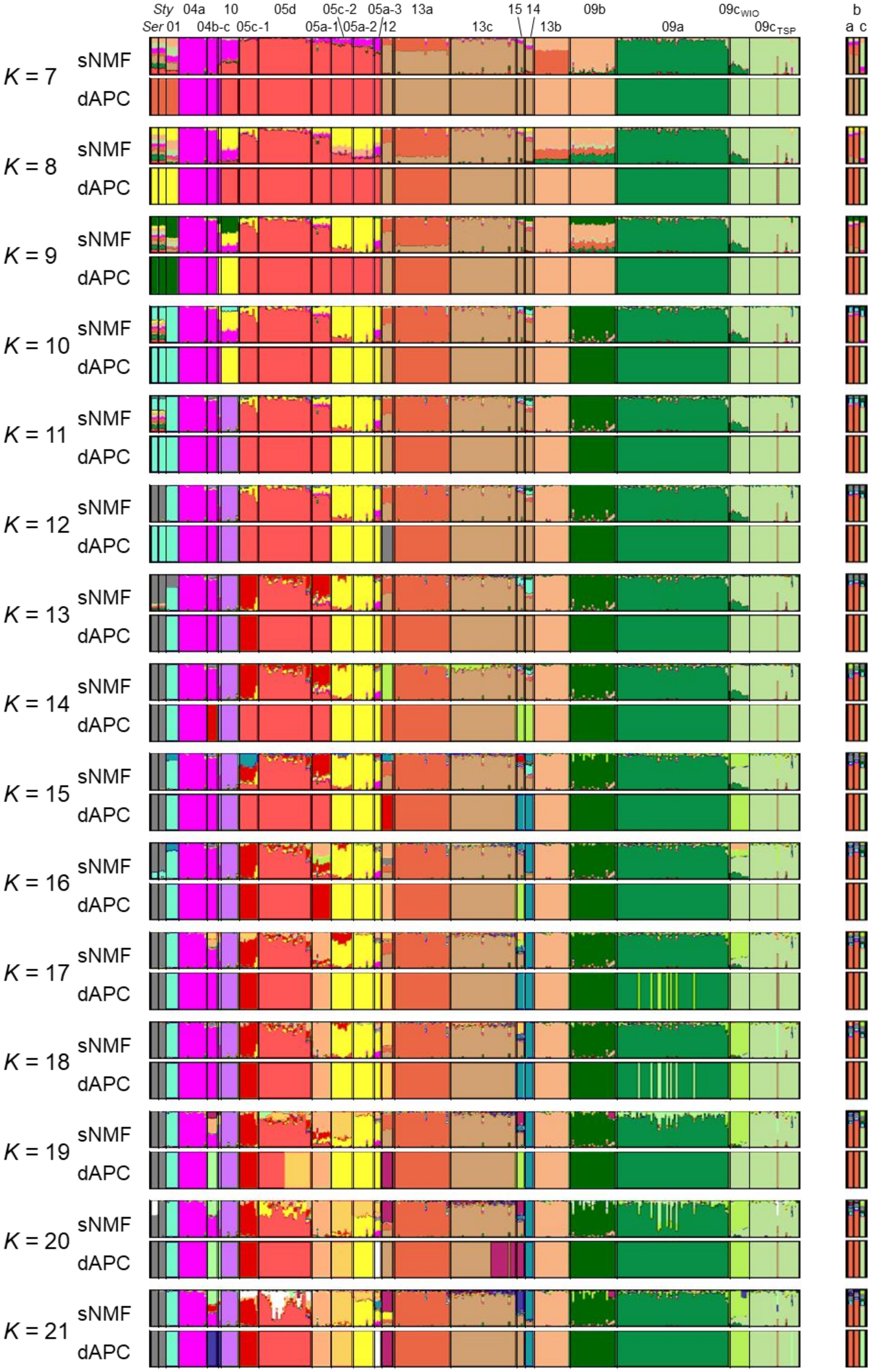
Sty 04a 10 05d 05c-2 05a-3 13a 15 14 09b 09c$_{WIO}$ b
Ser 01 04b-c 05c-1 05a-1 05a-2 12 13c 13b 09a 09c$_{TSP}$ a c
K = 7
K = 8
K = 9
K = 10
K = 11
K = 12
K = 13
K = 14
K = 15
K = 16
K = 17
K = 18
K = 19
K = 20
K = 21
sNMF
dAPC

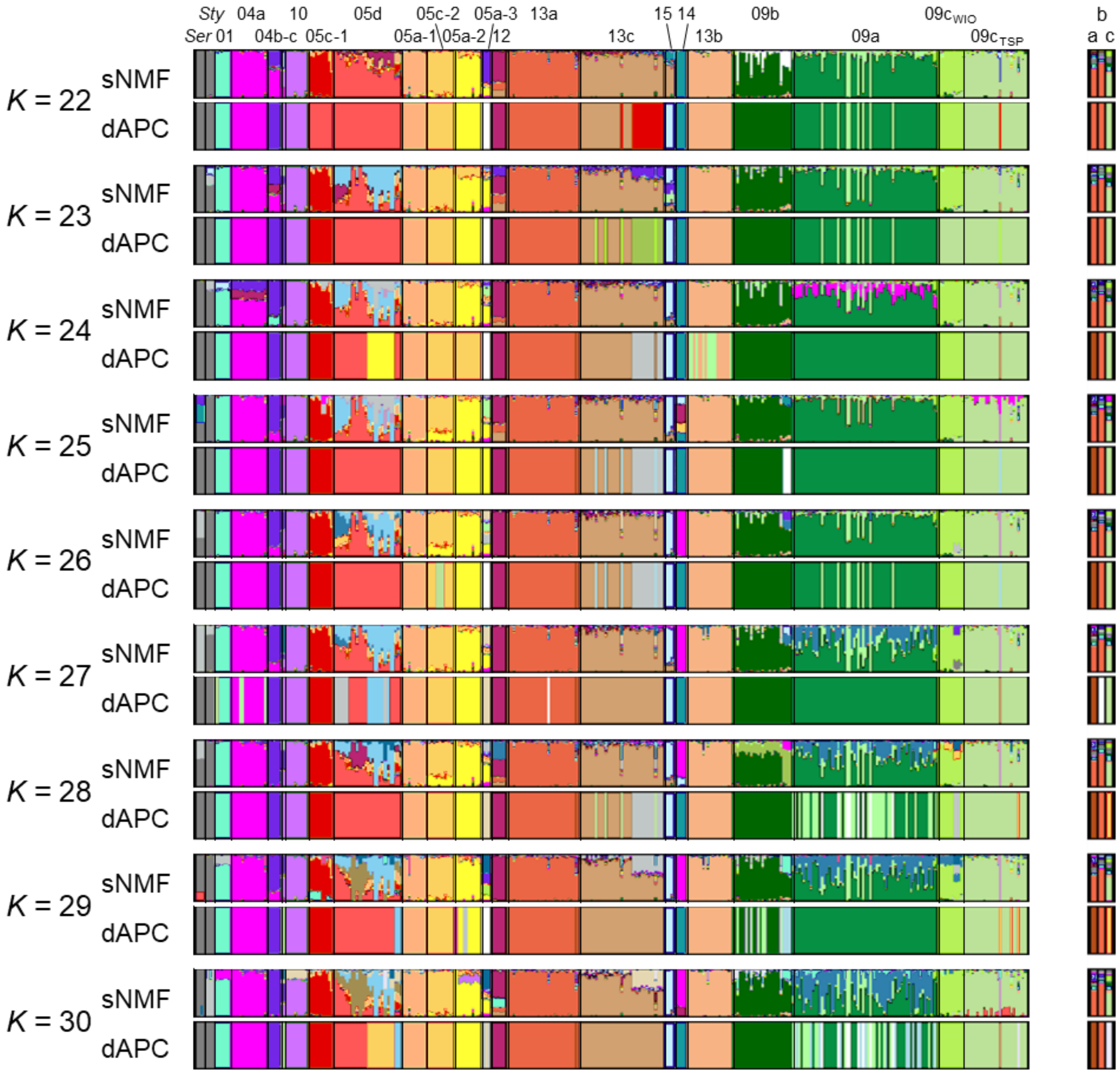

**Fig. S5** Networks based on Nei (1972) individual genetic distances for the datasets with (a) one SNP per locus (361 individuals × 1,559 SNPs) and (b) all SNPs (361 individuals × 17,465 SNPs), without outgroups: unrooted equal-angle split network (left panels) and minimum spanning tree (MST; right panels). Nodes and branches were coloured according *Pocillopora* genomic species hypotheses (GSHs). Published genomes are indicated by lowercase letters [a: *P. verrucosa* (Buitrago-López et al. 2020); b: *P. acuta* (Vidal-Dupiol et al. 2019); c: *P. damicornis* (Cunning et al. 2018)].

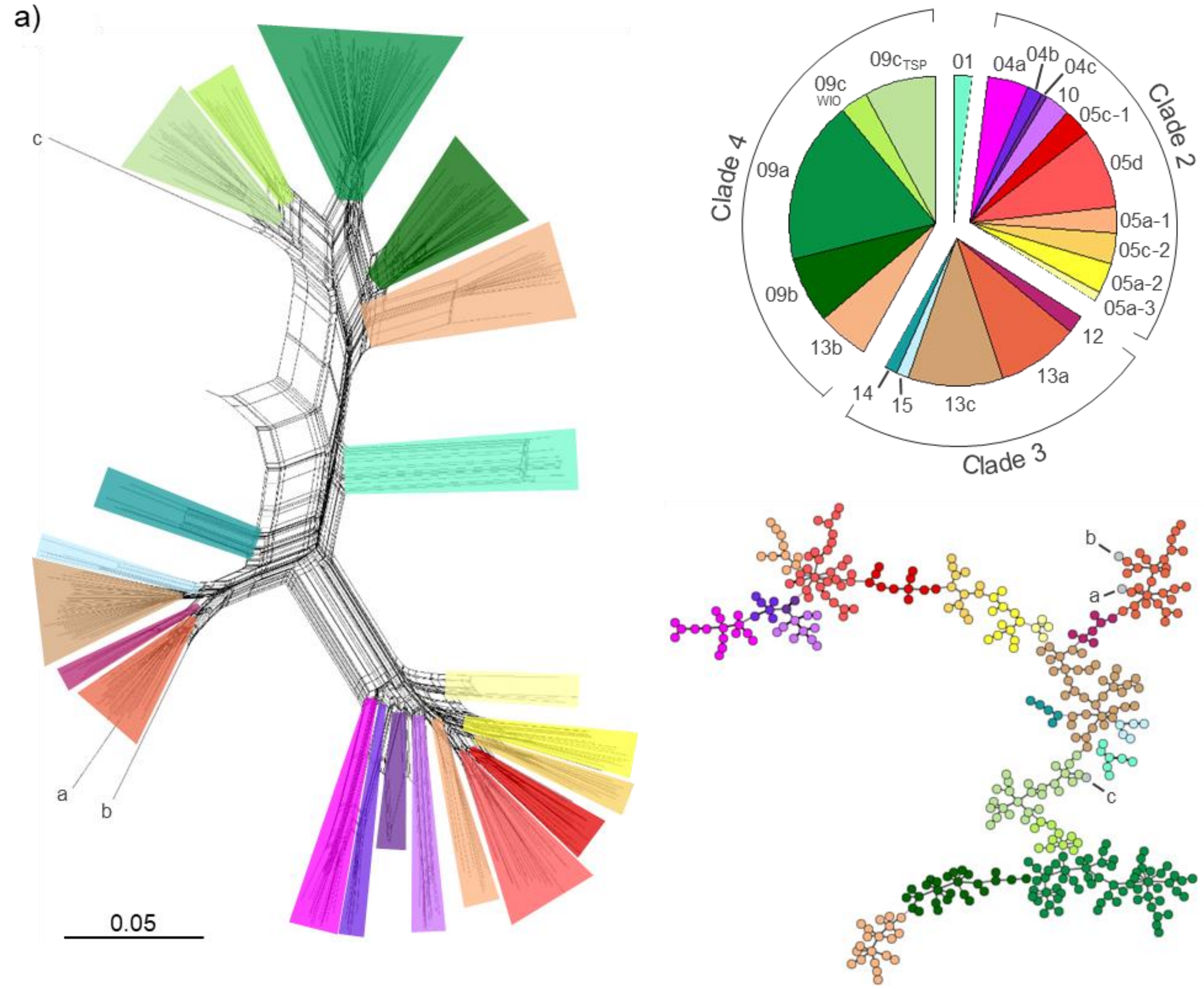

b)

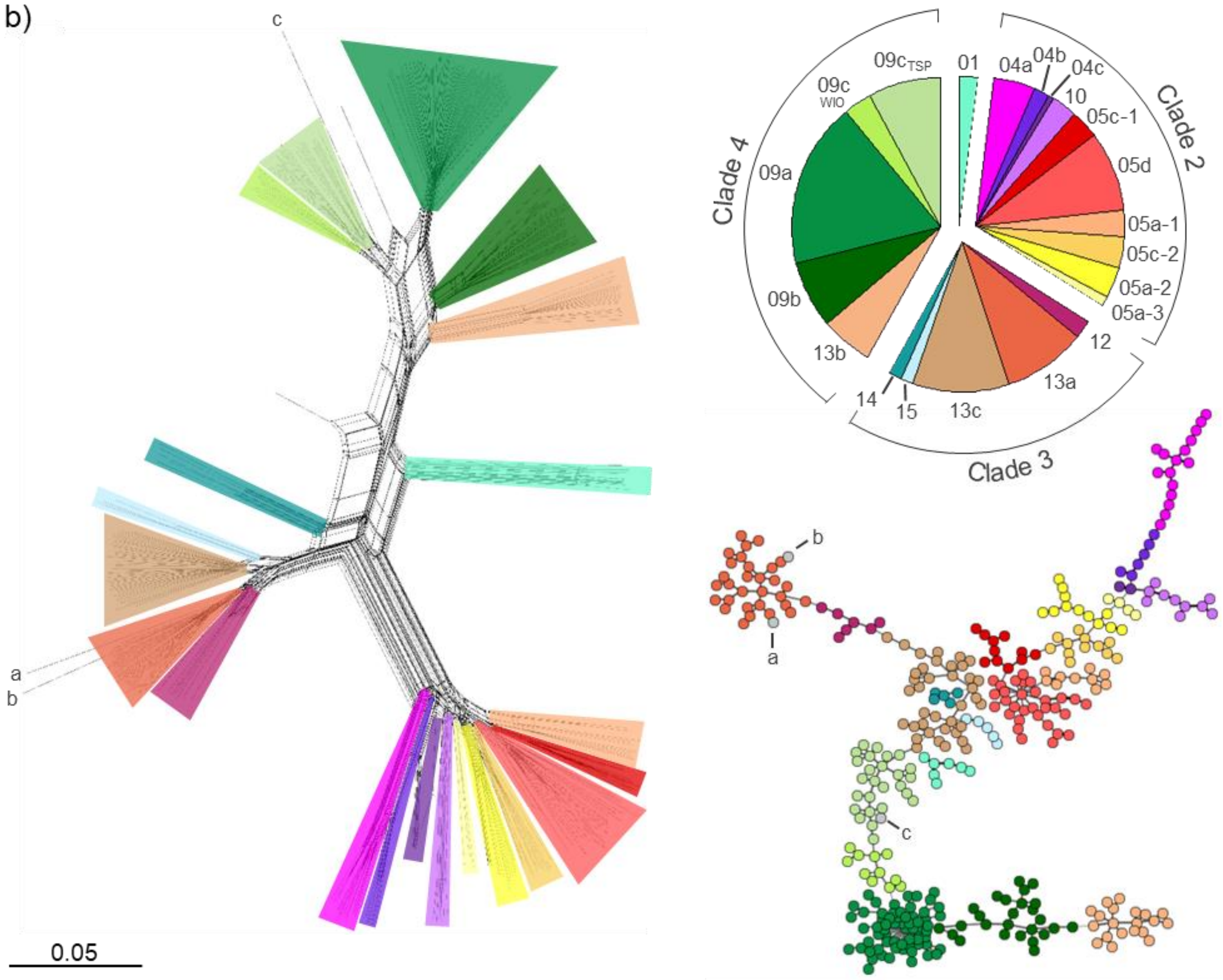

**Fig. S6** $F_{ST}$ (Weir and Cockerham 1984) between *Pocillopora* genomic species hypotheses (GSHs) for the datasets with (a) one SNP per locus (361 individuals × 1,559 SNPs) and (b) all SNPs (361 individuals × 17,465 SNPs). The heat map represents the $F_{ST}$ matrix (Table S7), with GSHs sorted according to the collapsed maximum-likelihood tree (ML-Tree; see also Fig. 1 & S2) in rows, and the $F_{ST}$ clustering in columns, for comparison. Main clades are separated by lines. *Ser*: *Seriatopora hystrix* and *Sty*: *Stylophora pistillata*.

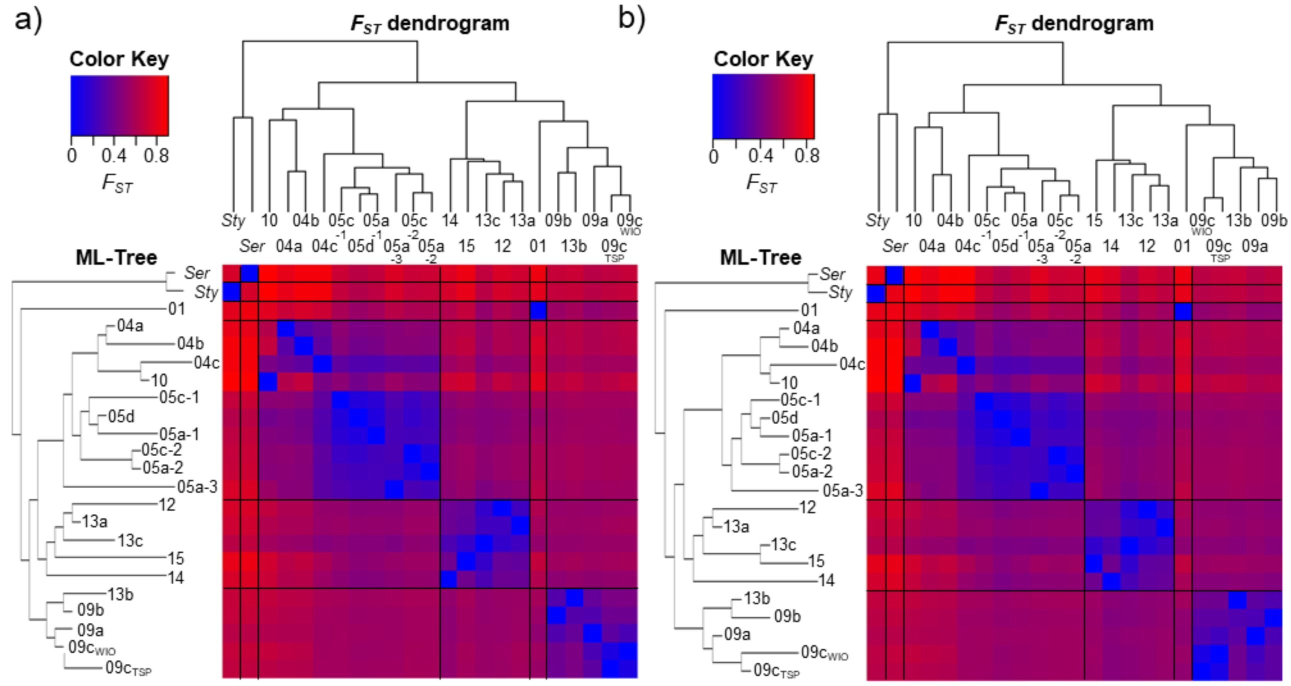

**Fig. S7** Maximum likelihood (ML) phylogenetic tree reconstructed with the PocHistone haplotypes, keeping one representative per haplotype (occurrence in parentheses). Values at nodes indicate bootstrap support (over 100 iterations).

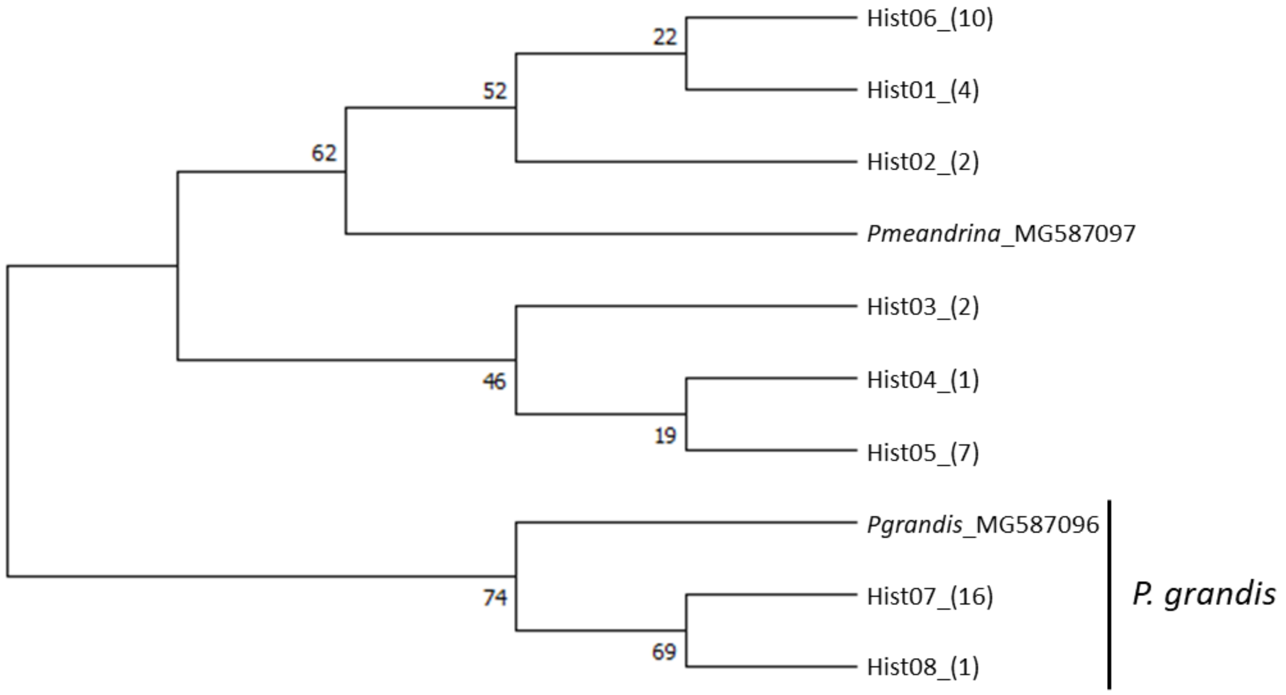

**Fig. S8** Alternative species tree topologies identified in the 95% highest posterior density (HPD) set for each best-supported species model: (a) Initial batch, (b) Clade 2 and (c) Clade 4 (numbers of individuals and SNPs used for each batch of scenarios indicated above). Node support (Bayesian posterior probability) and proportion of represented trees (out of $10^4$) are shown.

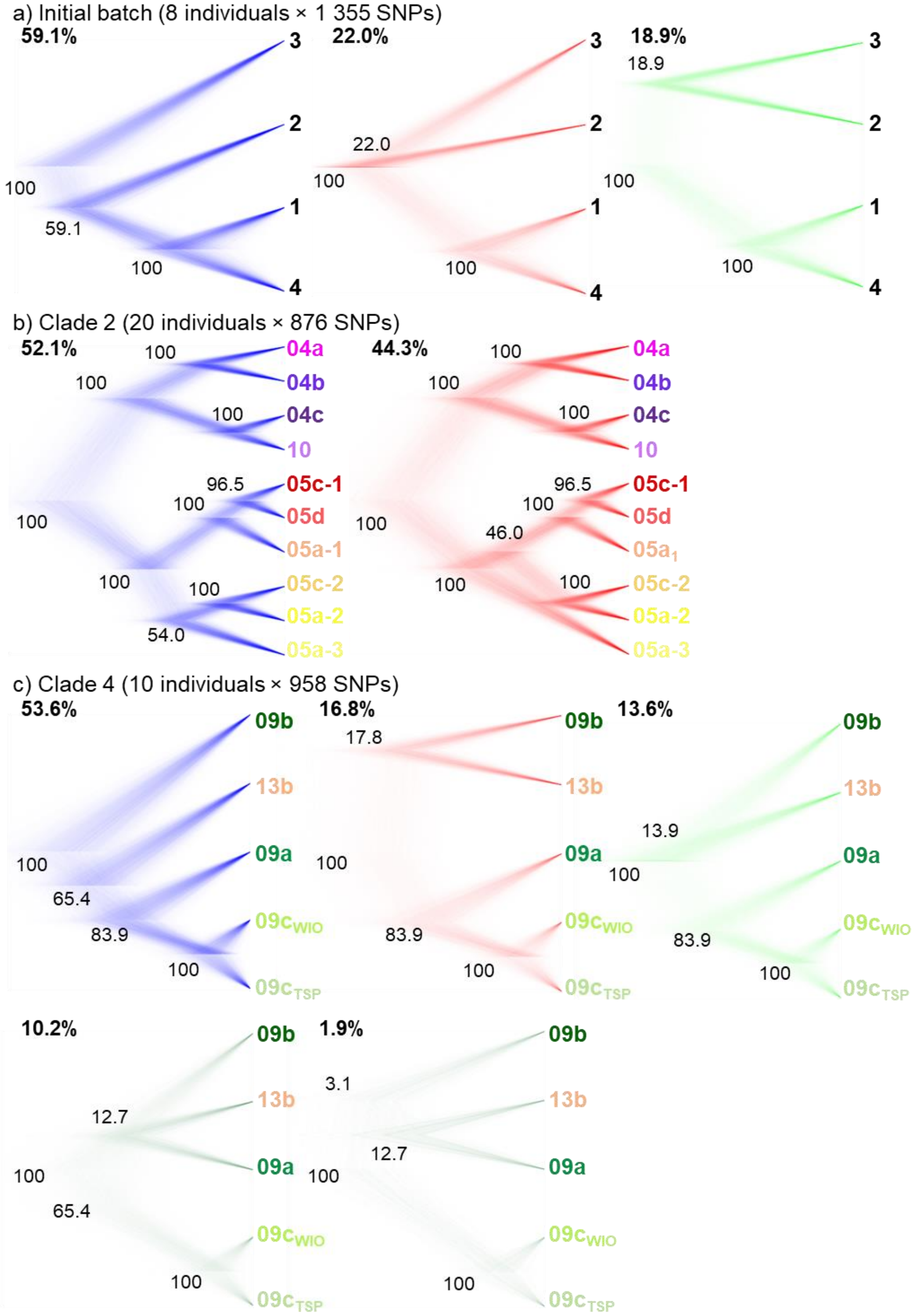
