## Appendix 3: Morphological Analyses for "From Genomics to Integrative Taxonomy? The Case Study of *Pocillopora* Corals"

Systematic Biology – Supporting Information

### **Appendix 3** Morphological Analyses.

#### *Macro- and Micromorphological Analyses*

In order to compare previously described morpho-species with genomic species hypotheses (GSHs), each colony was attributed a morphotype (or several when morphology was unclear), defined only by its *corallum* macromorphology [branch shape and thickness, size and uniformity of verrucae, and overall growth form as described in Veron (2000) and Schmidt-Roach et al. (2014)]. Morphotype identification was verified by sending a subset of photographs to three coral specialists (F. Benzoni, G. Faure, and D. Obura).

A subset of 10 colonies per GSH were also randomly selected for morphological observations and morphometric analyses. Fragments were bleached in ca. 1.8% sodium hypochlorite for 48h, then rinsed with distilled water and 90% ethanol to speed up air drying. Macromorphological images of the bleached skeletons were first captured at known magnificence, with a reference scale, using a Leica MC170 HD camera mounted on a Leica MZ16 stereomicroscope (Leica Camera AG, Wetzlar, Germany). Then, micromorphological observations were performed using scanning electron microscopy (SEM). Fragments were mounted on stubs using conductive double-sided adhesive carbon tapes, ca. 7 nm platinum sputter coated with a Leica ACE600 (Leica Camera AG, Wetzlar, Germany), and examined with a Hitachi SU3500 SEM (Hitachi High-Tech Analytical Science, Abingdon, UK) at the Plateforme technique de Microscopie Électronique (PtME) of the Muséum National d’Histoire Naturelle (MNHN, Paris, France). A collection of skeleton images was thus obtained for each selected specimen, and measurements were done with ImageJ2 (Rueden et al. 2017; https://imagej.nih.gov/ij/).

Seven characters were measured for each specimen (see Supplementary Methods for an illustration of each character; skeletal terms follow the glossary in Budd et al. 2012): (v1) maximum calice diameter, (v2) maximum calice diameter perpendicular to v1, (v3) distance between the center of the corallite and the center of the closest adjacent corallite, (v4) distance between denticles of the coenosteum, (v5) height of septa or septal teeth, (v6) maximum columella diameter and (v7) maximum columella diameter perpendicular to v6. All seven metrics were measured on 10 different corallites per specimen, except v4 (15 measures per specimen) and v5 (10 measures per corallite × 10 corallites per specimen), then averaged per specimen. A non-parametric permutational multivariate anova (PERMANOVA) was then performed using the R v4.0.4 (R Core Team 2021) library *'RVAideMemoire'* (Hervé 2021), with the GSHs as factor. Each metric was also analysed separately using a non-parametric permutational anova.

Two additional categorical variables were also considered: (v8) shape of septa and (v9) shape of columella (see Supplementary Methods for the different categories distinguished). A factorial analysis of mixed data (FAMD) was then performed for all nine variables using the R library *'FactoMineR'* (Lê et al. 2008). A reference specimen representative of each species enclosed in the latest *Pocillopora* taxonomic revision (Schmidt-Roach et al. 2014) was included by measuring the variables on the images incorporated.

**Supplementary Methods** Macro- and micromorphological analyses.

• **Glossary of morphological terms used in this study** (adapted from Budd et al. 2012):

Calice (-s): Cup-like structure at the distal end of solitary and outermost surface of solitary and, respectively, colonial coralla. Corresponding to the part of the skeleton occupied by a polyp.

Coenosteum (-a): Skeleton between corallites. May be covered with denticles.

Colony (-ies): A corallum consisting of two or more corallites whose polyps are integrated to different degree.

Columella (-e): Vertical axial structure within a corallite.

Corallite (-s): Skeleton of an individual within a colony.

Corallum (-a): Entire skeleton of a coral.

Septal tooth (teeth): Projections along the septal margins, extending from the septa and not formed by septal substitution.

Septum (-a): Radially-arranged vertical partition within a calice.

• **Illustration of the morphological characters considered in this study:**

| **Variable** | | **Type** | **Number of measures** |
|---|---|---|---|
| v1 | Maximum calice diameter | continuous | 10 corallites/specimen |
| v2 | Maximum calice diameter perpendicular to v1 | continuous | 10 corallites/specimen |
| v3 | Distance between the center of the corallite and the center of the closest adjacent corallite | continuous | 10 corallites/specimen |
| v4 | Distance between denticles of the coenosteum | continuous | 15/specimen |
| v5 | Height of septa or septal teeth | continuous | 10/corallite × 10 corallites/specimen |
| v6 | Maximum columella diameter | continuous | 10 corallites/specimen |
| v7 | Maximum columella diameter perpendicular to v6 | continuous | 10 corallites/specimen |
| v8 | Shape of septa | categorical | N/A |
| v9 | Shape of columella | categorical | N/A |

(v1-v7) Numeric variables

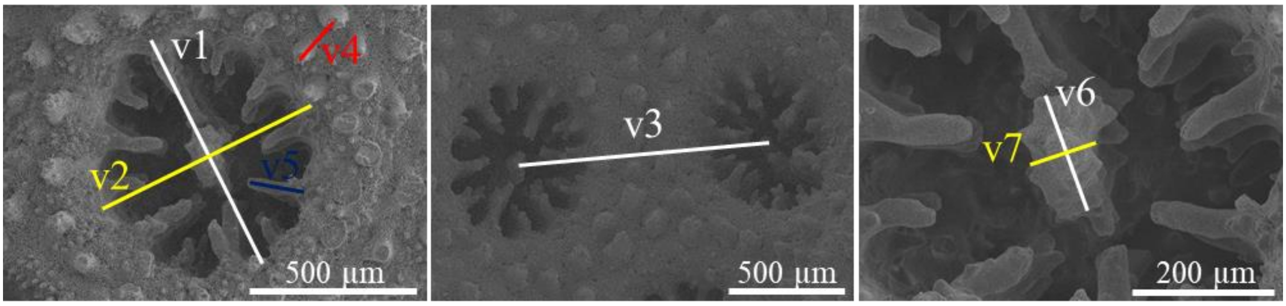

(v8) Shape of septa

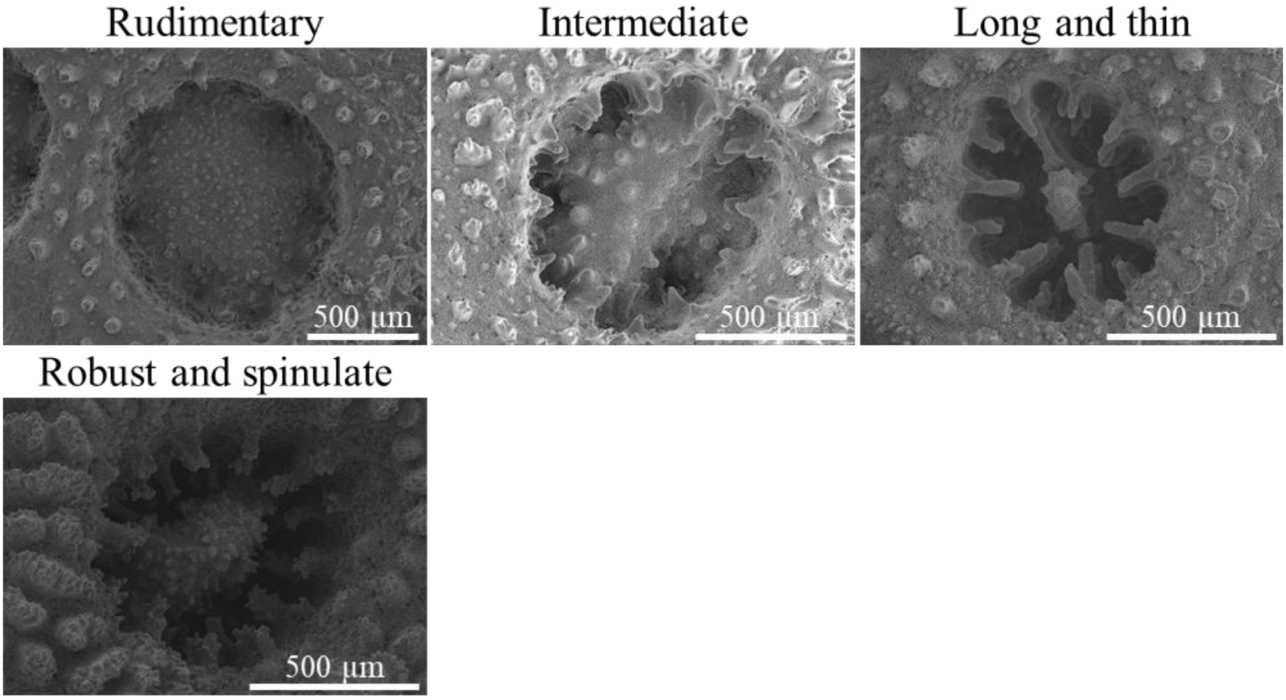

(v9) Shape of columella

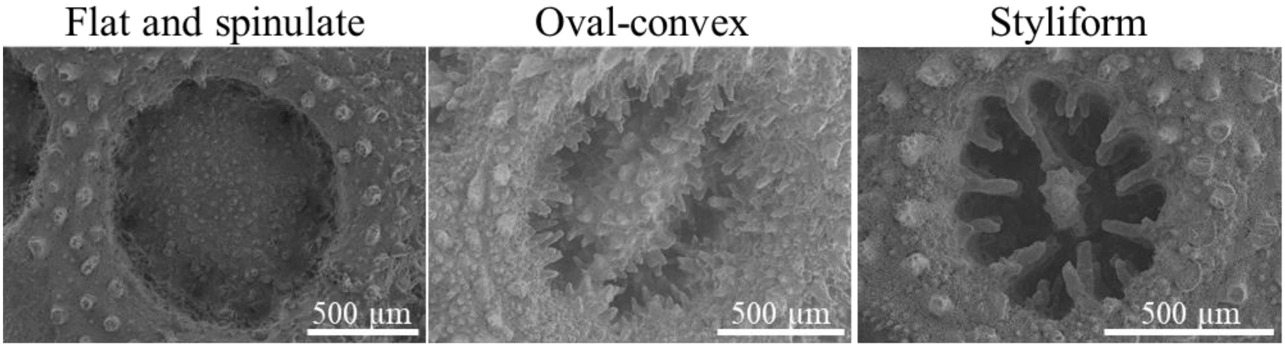

**Fig. S9** Morphometry of *Pocillopora* colonies. (a) Box plots of the numeric morphological variables (v1-v7; see Supp. Methods) for each genomic species hypothesis (GSH). Diamonds represent means, and boxes colours indicate GSHs (labelled below; number of colonies in parentheses). Letters above each box denote significance groups according to pairwise permutational t tests [i.e., GSHs sharing the same letter are not significantly different ($P$ < 0.05)]. (b) Means (± s.e.) of the numeric morphological variables (v1-v7; see Supp. Methods), for each GSH separately (labelled below; number of colonies in parentheses). Letters above means of each GSH denote significance groups according to pairwise permutational multivariate anova [i.e., GSHs sharing the same letter are not significantly different ($P$ < 0.05)].

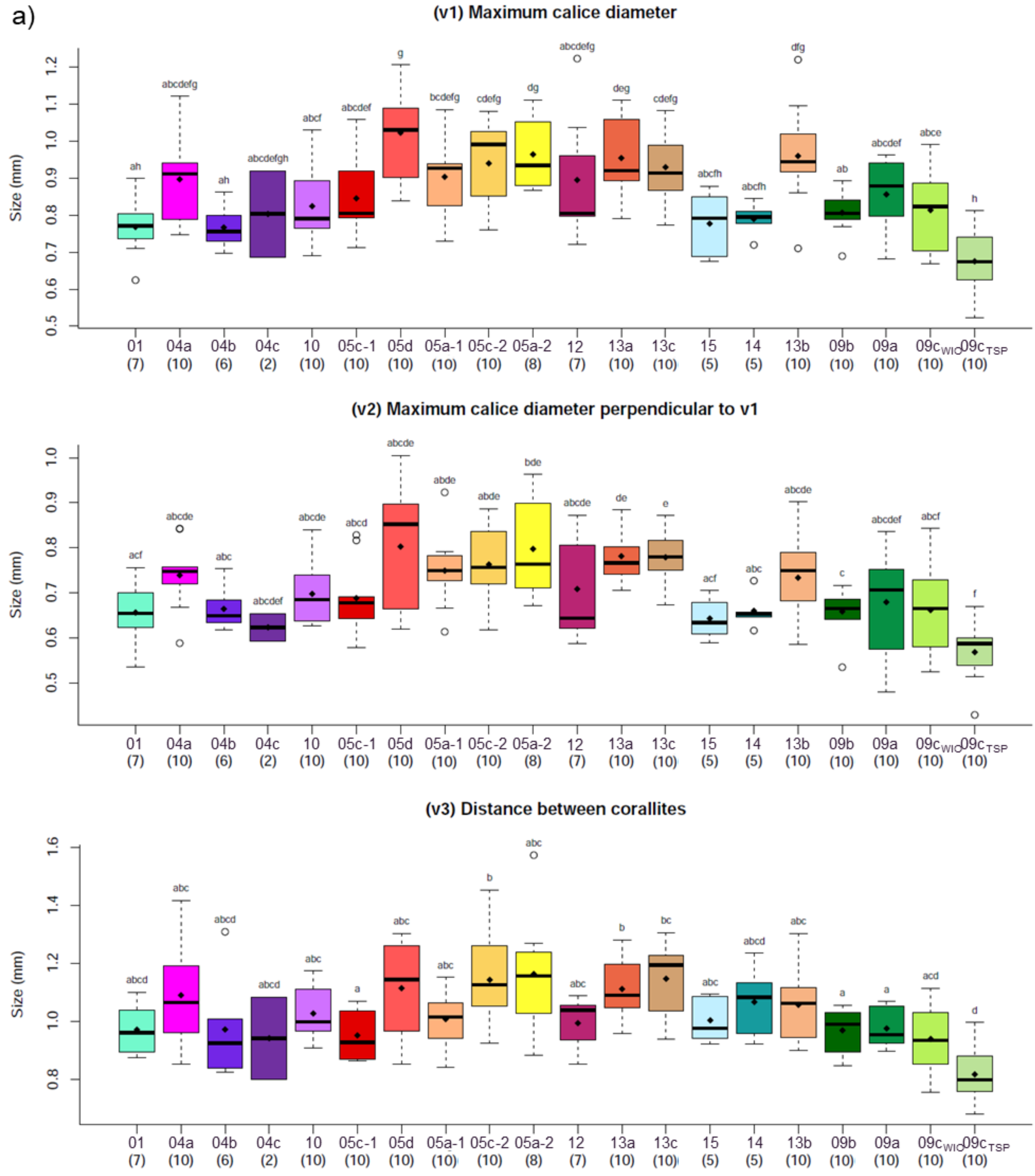

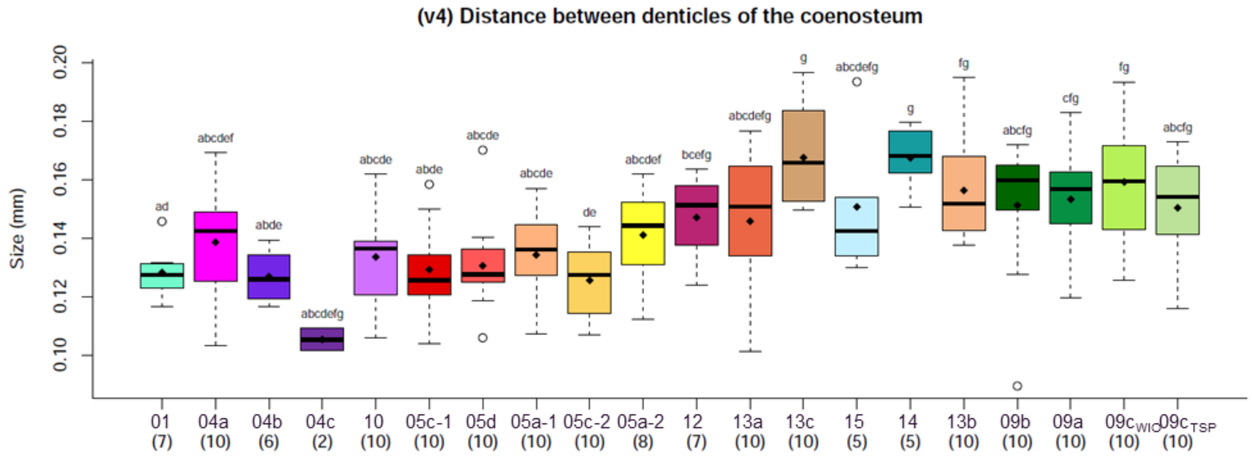
(v4) Distance between denticles of the coenosteum
Size (mm)
01 (7) 04a (10) 04b (6) 04c (2) 10 (10) 05c-1 (10) 05d (10) 05a-1 (10) 05c-2 (10) 05a-2 (8) 12 (7) 13a (10) 13c (10) 15 (5) 14 (5) 13b (10) 09b (10) 09a (10) 09cWIO (10) 09cTSP (10)

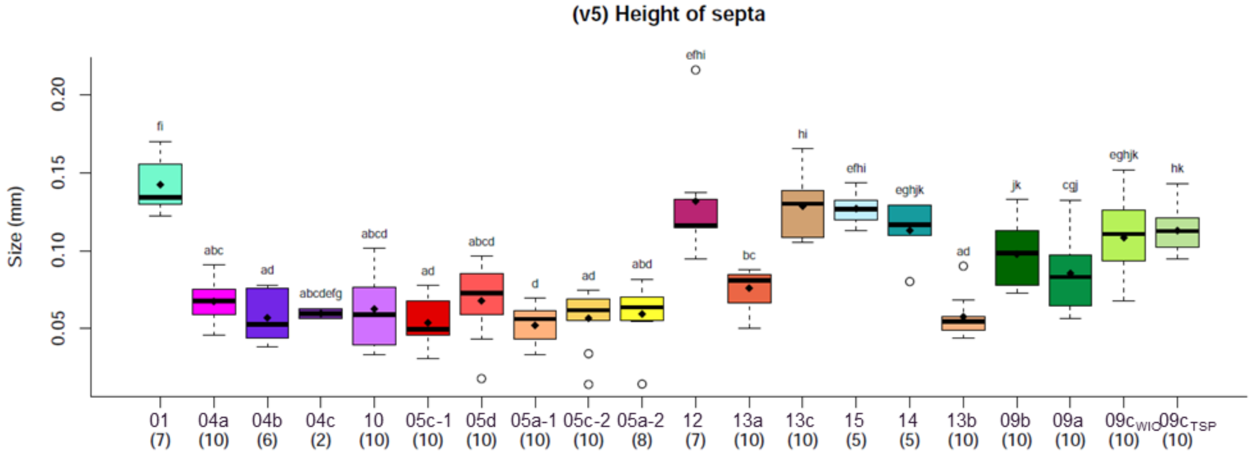
(v5) Height of septa
Size (mm)
01 (7) 04a (10) 04b (6) 04c (2) 10 (10) 05c-1 (10) 05d (10) 05a-1 (10) 05c-2 (10) 05a-2 (8) 12 (7) 13a (10) 13c (10) 15 (5) 14 (5) 13b (10) 09b (10) 09a (10) 09cWIO (10) 09cTSP (10)

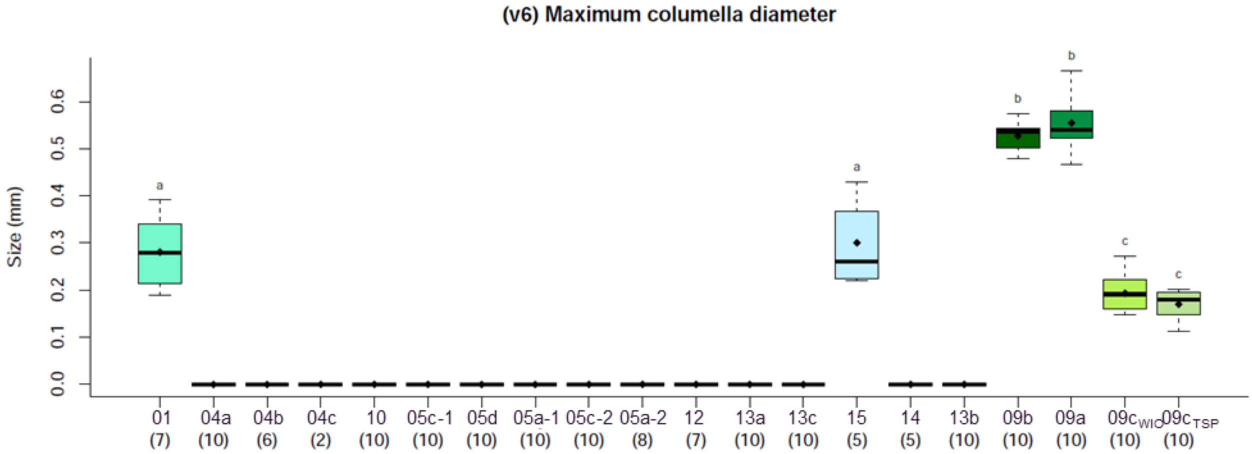
(v6) Maximum columella diameter
Size (mm)
01 (7) 04a (10) 04b (6) 04c (2) 10 (10) 05c-1 (10) 05d (10) 05a-1 (10) 05c-2 (10) 05a-2 (8) 12 (7) 13a (10) 13c (10) 15 (5) 14 (5) 13b (10) 09b (10) 09a (10) 09cWIO (10) 09cTSP (10)

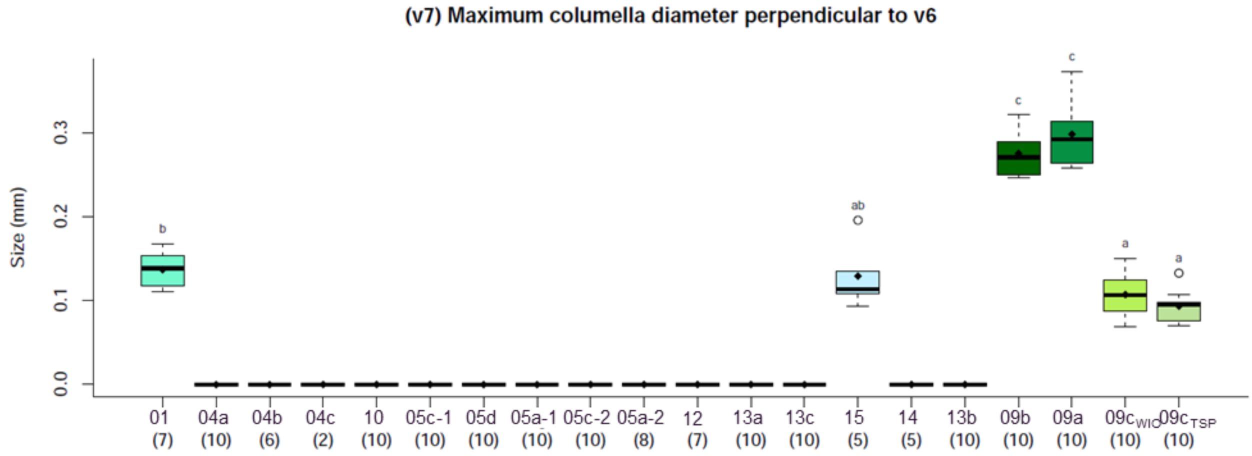
(v7) Maximum columella diameter perpendicular to v6
Size (mm)
01 (7) 04a (10) 04b (6) 04c (2) 10 (10) 05c-1 (10) 05d (10) 05a-1 (10) 05c-2 (10) 05a-2 (8) 12 (7) 13a (10) 13c (10) 15 (5) 14 (5) 13b (10) 09b (10) 09a (10) 09cWIO (10) 09cTSP (10)

b)

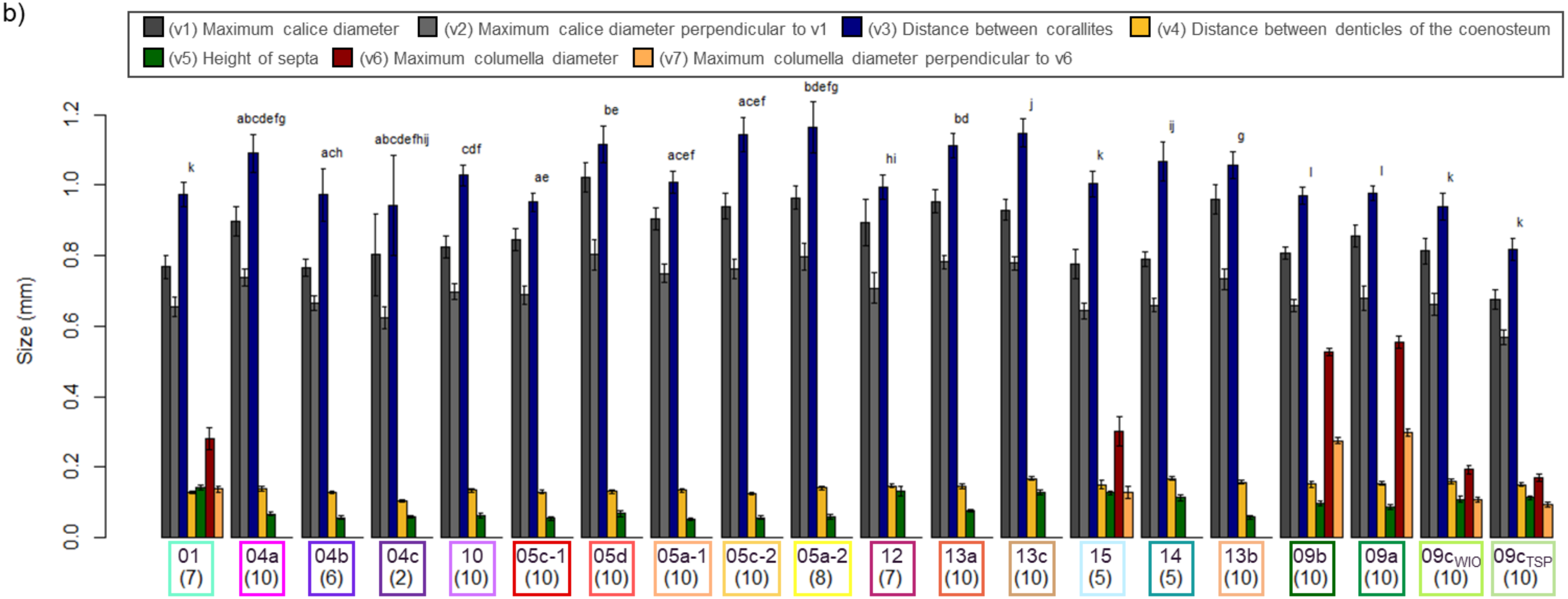

(v1) Maximum calice diameter
(v2) Maximum calice diameter perpendicular to v1
(v3) Distance between corallites
(v4) Distance between denticles of the coenosteum
(v5) Height of septa
(v6) Maximum columella diameter
(v7) Maximum columella diameter perpendicular to v6
Size (mm)
0.0
0.2
0.4
0.6
0.8
1.0
1.2
k
abcdefg
ach
abcdefhij
cdf
ae
be
acef
acef
bdefg
hi
bd
j
k
ij
g
l
l
k
k
01 (7)
04a (10)
04b (6)
04c (2)
10 (10)
05c-1 (10)
05d (10)
05a-1 (10)
05c-2 (10)
05a-2 (8)
12 (7)
13a (10)
13c (10)
15 (5)
14 (5)
13b (10)
09b (10)
09a (10)
09c$_{WIO}$ (10)
09c$_{TSP}$ (10)

**Fig. S10** Plots of the first three principal components (PC) from the factorial analysis of mixed data (FAMD) with the nine morphological variables from all *Pocillopora* specimens examined ($N = 170$). Individuals are coloured according to the genomic species hypotheses (GSHs) previously identified, and those from the same GSH are enclosed by dashed polygons. Specimens from Schmidt-Roach et al. (2014) are represented by triangles and coloured according to the most probable GSH identification (based on mitochondrial open reading frame haplotypes).

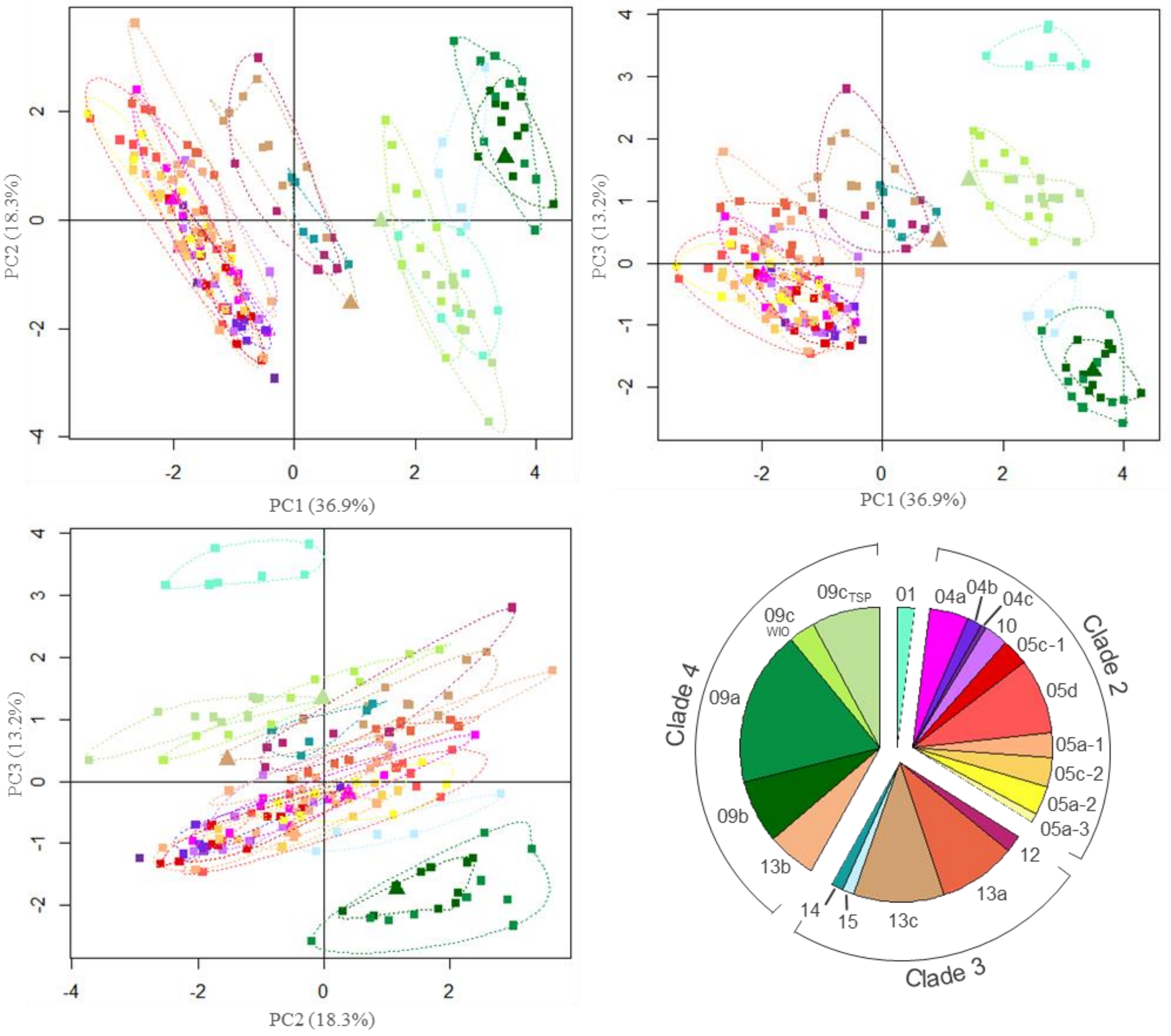
