## Appendix 4: Symbiodiniaceae Analyses for "From Genomics to Integrative Taxonomy? The Case Study of *Pocillopora* Corals"

---

### **Appendix 4** Symbiodiniaceae Analyses.

#### ***Characterisation of Associated Symbiodiniaceae***

Symbiodiniaceae communities were characterised for a subset of colonies [ca. 15 per genomic species hypothesis (GSH) when available; including three replicates] by high-throughput sequencing the ribosomal RNA internal transcribed spacer 2 (ITS2). Fragments of ca. 350 bp were amplified using the itsD (Pochon et al. 2001) and its2rev2 (Stat et al. 2009) primers tagged with internal indexing tails (see Table S2 in Appendix 1 for the PCR conditions). The resulting PCR products were sent to the plateforme iGenSeq (ICM, Paris, France) for PE250 sequencing with an Illumina MiSeq platform (Illumina, San Diego, CA).

Sequence reads were processed with the SAMBA v3.0.1 (Standardized and Automated MetaBarcoding Analyses) workflow (https://github.com/ifremer-bioinformatics/samba), developed by SeBiMER (Ifremer Bioinformatics Core Facility, Ifremer, France) and implemented in NextFlow v20.04.1 (Di Tommaso et al. 2017). Briefly, reads were quality controlled and checked using a custom python script, then primers were trimmed with cutadapt v2.1 (Martin 2011), implemented in QIIME 2 v2019.10.0 (Bolyen et al. 2019). Paired reads were merged with the R v4.0.4 (R Core Team 2021) library *'DADA2'* (Callahan et al. 2016) and chimeric sequences were removed. Then, operational taxonomic units (OTUs) were identified following a distribution-based clustering (Preheim et al. 2013) of the amplicon sequence variants (ASVs) with the dbOTU3 algorithm from QIIME 2. Finally, the resulting OTUs were taxonomically assigned by querying a custom reference database of Symbiodiniaceae ITS2 adapted from the one available in SymPortal (downloaded on 13/01/2022; Hume et al. 2019). Taxonomic affiliations of the OTUs were confirmed by reconstructing the phylogenetic relationships among them using MAFFT v7.713 (Katoh and Standley 2013) to produce sequence alignment and FastTree v2.1.11 (GTR+CAT model; Price et al. 2009) to compute tree with the approximately maximum-likelihood (ML) method. The final OTU table was produced in a standard BIOM format for subsequent analyses.

OTU table, sample metadata, and taxonomic data were imported into R using the *'phyloseq'* library (McMurdie and Holmes 2013) to facilitate downstream analyses that were performed with the R library *'vegan'* (Oksanen et al. 2020). OTUs and individuals with less than 10 and 500 sequences, respectively, were removed to reduce possible sequencing errors. Then, alpha diversity metrics (Chao1 and Shannon) were computed at the OTU level and compared using non-parametric permutational ANOVA performed with the R library *'RVAideMemoire'* (Hervé 2021), with the GSHs or the localities as factor. Finally, a nonmetric multidimensional scaling (NMDS) using Bray and Curtis (1957) dissimilarity index was performed to assess community similarity.

### References

Bolyen E., Rideout J.R., Dillon M.R., Bokulich N.A., Abnet C.C., Al-Ghalith G.A., Alexander H., Alm E.J., Arumugam M., Asnicar F., Bai Y., Bisanz J.E., Bittinger K., Brejnrod A., Brislawn C.J., Brown C.T., Callahan B.J., Caraballo-Rodríguez A.M., Chase J., Cope E.K., Da Silva R., Diener C., Dorrestein P.C., Douglas G.M., Durall D.M., Duvallet C., Edwardson C.F., Ernst M., Estaki M., Fouquier J., Gauglitz J.M., Gibbons S.M., Gibson D.L., Gonzalez A., Gorlick K., Guo J., Hillmann B., Holmes S., Holste H., Huttenhower C., Huttley G.A., Janssen S., Jarmusch A.K., Jiang L., Kaehler B.D., Kang K.B., Keefe C.R., Keim P., Kelley S.T., Knights D., Koester I., Kosciolek T., Kreps J., Langille M.G.I., Lee J., Ley R., Liu Y.-X., Loftfield E., Lozupone C., Maher M., Marotz C., Martin B.D., McDonald D., McIver L.J., Melnik A.V., Metcalf J.L., Morgan S.C., Morton J.T., Naimey A.T., Navas-Molina J.A., Nothias L.F., Orchanian S.B., Pearson T., Peoples S.L., Petras D., Preuss M.L., Pruesse E., Rasmussen L.B., Rivers A., Robeson M.S., Rosenthal P., Segata N., Shaffer M., Shiffer A., Sinha R., Song S.J., Spear J.R., Swafford A.D., Thompson L.R., Torres P.J., Trinh P., Tripathi A., Turnbaugh P.J., Ul-Hasan S., van der Hooft J.J.J., Vargas F., Vázquez-Baeza Y., Vogtmann E., von Hippel M., Walters W., Wan Y., Wang M., Warren J., Weber K.C., Williamson C.H.D., Willis A.D., Xu Z.Z., Zaneveld J.R., Zhang Y., Zhu Q., Knight R., Caporaso J.G. 2019. Reproducible, interactive, scalable and extensible microbiome data science using QIIME 2. Nat. Biotechnol. 37:852–857.

Bray J.R., Curtis J.T. 1957. An ordination of the upland forest communities of southern Wisconsin. Ecol. Monogr. 27:326–349.

Callahan B.J., McMurdie P.J., Rosen M.J., Han A.W., Johnson A.J.A., Holmes S.P. 2016. DADA2: high-resolution sample inference from Illumina amplicon data. Nat. Methods. 13:581–583.

Di Tommaso P., Chatzou M., Floden E.W., Barja P.P., Palumbo E., Notredame C. 2017. Nextflow enables reproducible computational workflows. Nat. Biotechnol. 35:316–319.

Hervé M. 2021. RVAideMemoire: testing and plotting procedures for biostatistics. R package version 0.9-64.

Hume B.C.C., Smith E.G., Ziegler M., Warrington H.J.M., Burt J.A., LaJeunesse T.C., Wiedenmann J., Voolstra C.R. 2019. SymPortal: a novel analytical framework and platform for coral algal symbiont next-generation sequencing ITS2 profiling. Mol. Ecol. Resour. 19:1063–1080.

Katoh K., Standley D.M. 2013. MAFFT multiple sequence alignment software version 7: improvements in performance and usability. Mol. Biol. Evol. 30:772–780.

Martin M. 2011. Cutadapt removes adapter sequences from high-throughput sequencing reads. EMBnet J. 17:10–12.

McMurdie P.J., Holmes S. 2013. phyloseq: an R package for reproducible interactive analysis and graphics of microbiome census data. PLoS ONE. 8:61217.

Oksanen J., Blanchet F.G., Friendly M., Kindt R., Legendre P., McGlinn D., Minchin P.R., O'Hara R.B., Simpson G.L., Solymos P., Stevens M.H.H., Szoecs E., Wagner H. 2020. vegan: community ecology package. R package verson 2.5-7.

Pochon X., Pawlowski J., Zaninetti L., Rowan R. 2001. High genetic diversity and relative specificity among *Symbiodinium*-like endosymbiotic dinoflagellates in soritid foraminiferans. Mar. Biol. 139:1069–1078.

Preheim S.P., Perrotta A.R., Martin-Platero A.M., Gupta A., Alm E.J. 2013. Distribution-based clustering: using ecology to refine the operational taxonomic unit. Appl. Environ. Microbiol. 79:6593–6603.

Price M.N., Dehal P.S., Arkin A.P. 2009. FastTree: computing large minimum evolution trees with profiles instead of a distance matrix. Mol. Biol. Evol. 26:1641–1650.

R Core Team. 2021. R: a language and environment for statistical computing. R Foundation for Statistical Computing, Vienna, Austria. URL: https://www.R-project.org/: .

Stat M., Pochon X., Cowie R., Gates R. 2009. Specificity in communities of *Symbiodinium* in corals from Johnston Atoll. Mar. Ecol. Prog. Ser. 386:83–96.

**Fig. S11** Phylogenetic relationships among Symbiodiniaceae operational taxonomic units (OTUs): maximum-likelihood (ML) phylogenetic tree of the 534 identified OTUs. Main clades are indicated.

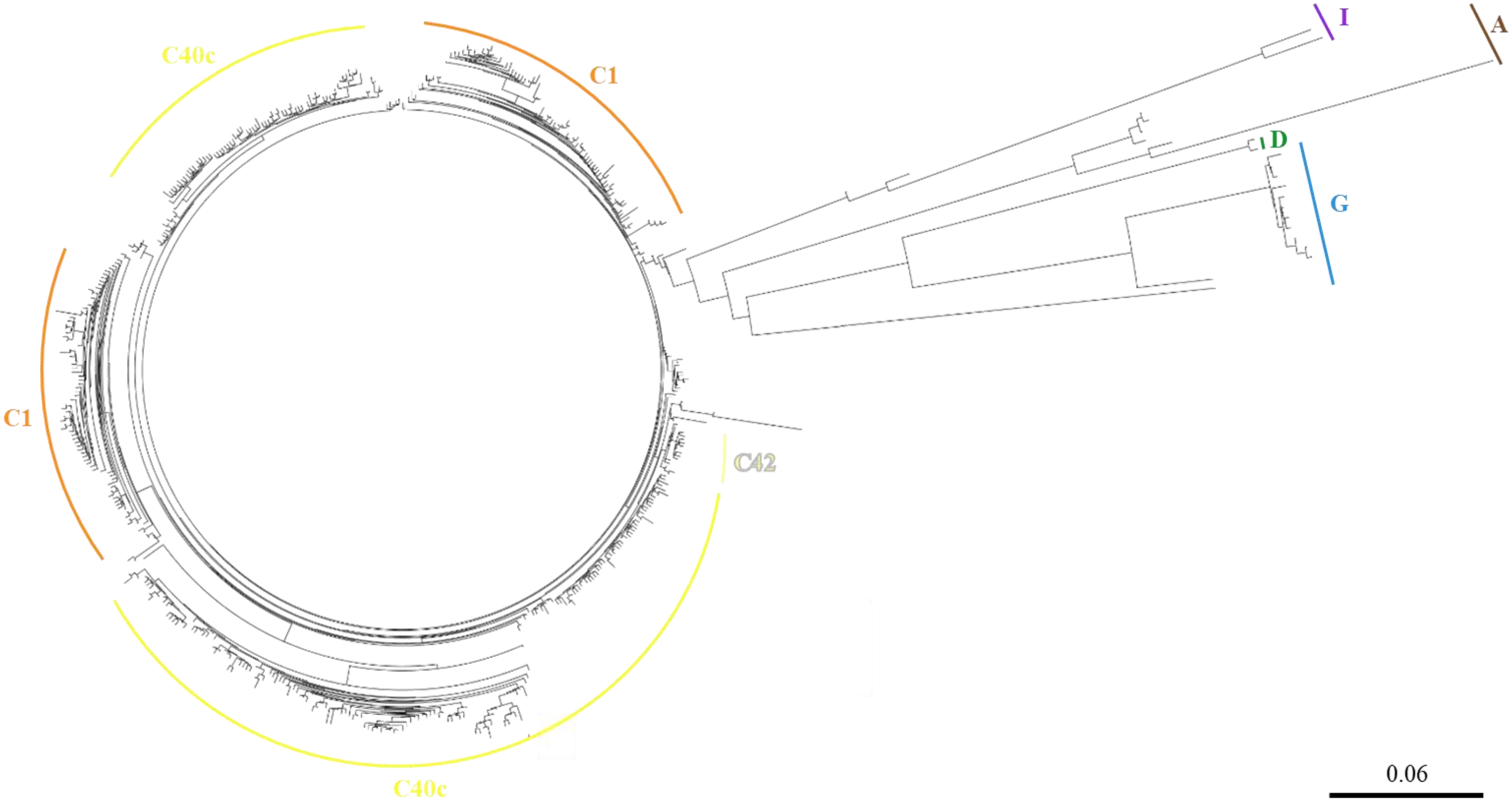

**Fig. S12** Symbiodiniaceae alpha diversity index [(a,c) Shannon and (b,d) Chao1] within samples grouped (a,b) per genomic species hypothesis (GSH) and (c,d) per sampling localities (indicated below; number of samples in parentheses). Diamonds represent means and letters above each box denote significance groups according to pairwise permutational t tests [i.e., boxes sharing the same letter are not significantly different ($P < 0.05$)].

MAY: Mayotte, JDN: Juan de Nova Island, EUR: Europa Island, MADne, MADnw and MADsw: northeastern, northwestern and southwestern Madagascar, respectively, REU: Reunion Island, ROD: Rodrigues Island, TRO: Tromelin Island, CHE: Chesterfield Islands, NCAw and NCAe: western and eastern Grande Terre (New Caledonia), respectively, LOY: Loyalty Islands (New Caledonia), and PYF: French Polynesia (Bora-Bora, Moorea and Tahiti).

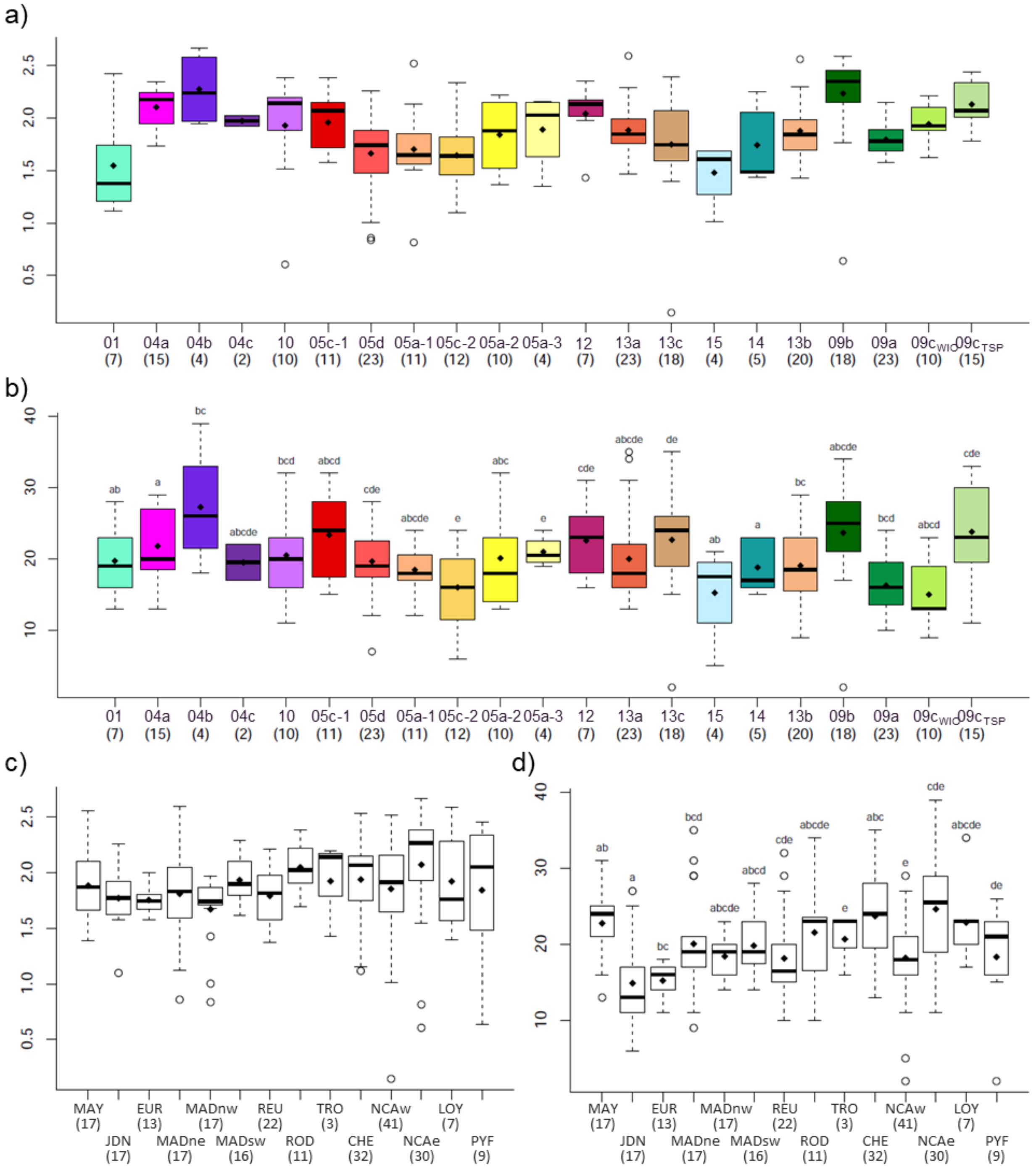

**Fig. S13** Proportion of Symbiodiniaceae most represented taxa for each sample, sorted (a) per genomic species hypothesis (GSH) and (b) per sampling locality (indicated below; number of samples in parentheses).

MAY: Mayotte, JDN: Juan de Nova Island, EUR: Europa Island, MADne, MADnw and MADsw: northeastern, northwestern and southwestern Madagascar, respectively, REU: Reunion Island, ROD: Rodrigues Island, TRO: Tromelin Island, CHE: Chesterfield Islands, NCAw and NCAe: western and eastern Grande Terre (New Caledonia), respectively, LOY: Loyalty Islands (New Caledonia), and PYF: French Polynesia (Bora-Bora, Moorea and Tahiti).

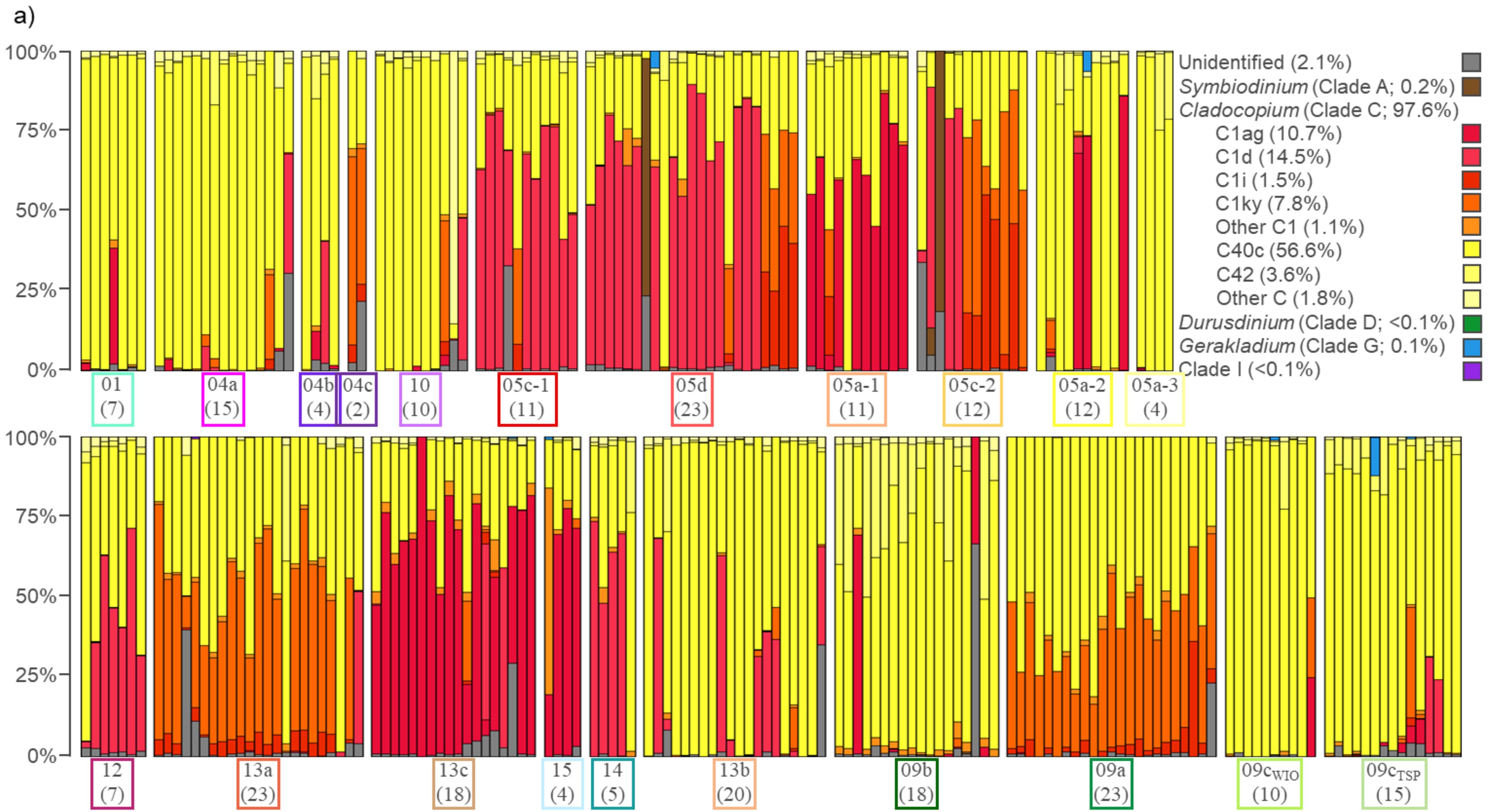

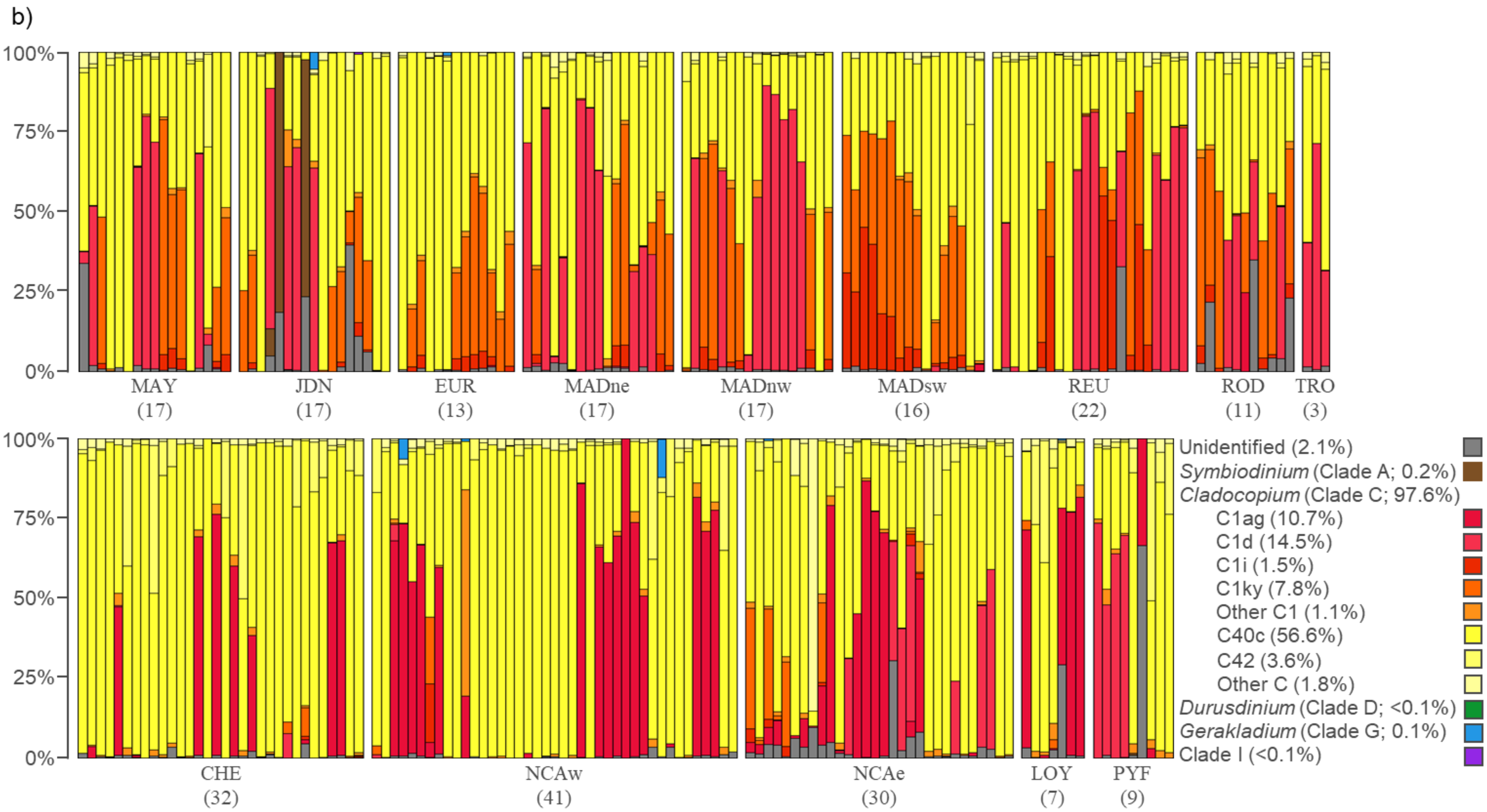

b)
100%
75%
50%
25%
0%
MAY (17)
JDN (17)
EUR (13)
MADne (17)
MADnw (17)
MADsw (16)
REU (22)
ROD (11)
TRO (3)
CHE (32)
NCAw (41)
NCAe (30)
LOY (7)
PYF (9)
Unidentified (2.1%)
Symbiodinium (Clade A; 0.2%)
Cladocopium (Clade C; 97.6%)
C1ag (10.7%)
C1d (14.5%)
C1i (1.5%)
C1ky (7.8%)
Other C1 (1.1%)
C40c (56.6%)
C42 (3.6%)
Other C (1.8%)
Durusdinium (Clade D; <0.1%)
Gerakladium (Clade G; 0.1%)
Clade I (<0.1%)

**Fig. S14** *Pocillopora* associated Symbiodiniaceae communities. Plots of the first three principal components (PC) from the nonmetric multidimensional scaling (NMDS) using the Bray and Curtis (1957) dissimilarity index between all pairs of samples, coloured (a) by genomic species hypothesis (GSH) and (b) by locality. Dashed squares indicate dominant Symbiodiniaceae taxa and polygons frame 75% of the individuals from a single GSH.

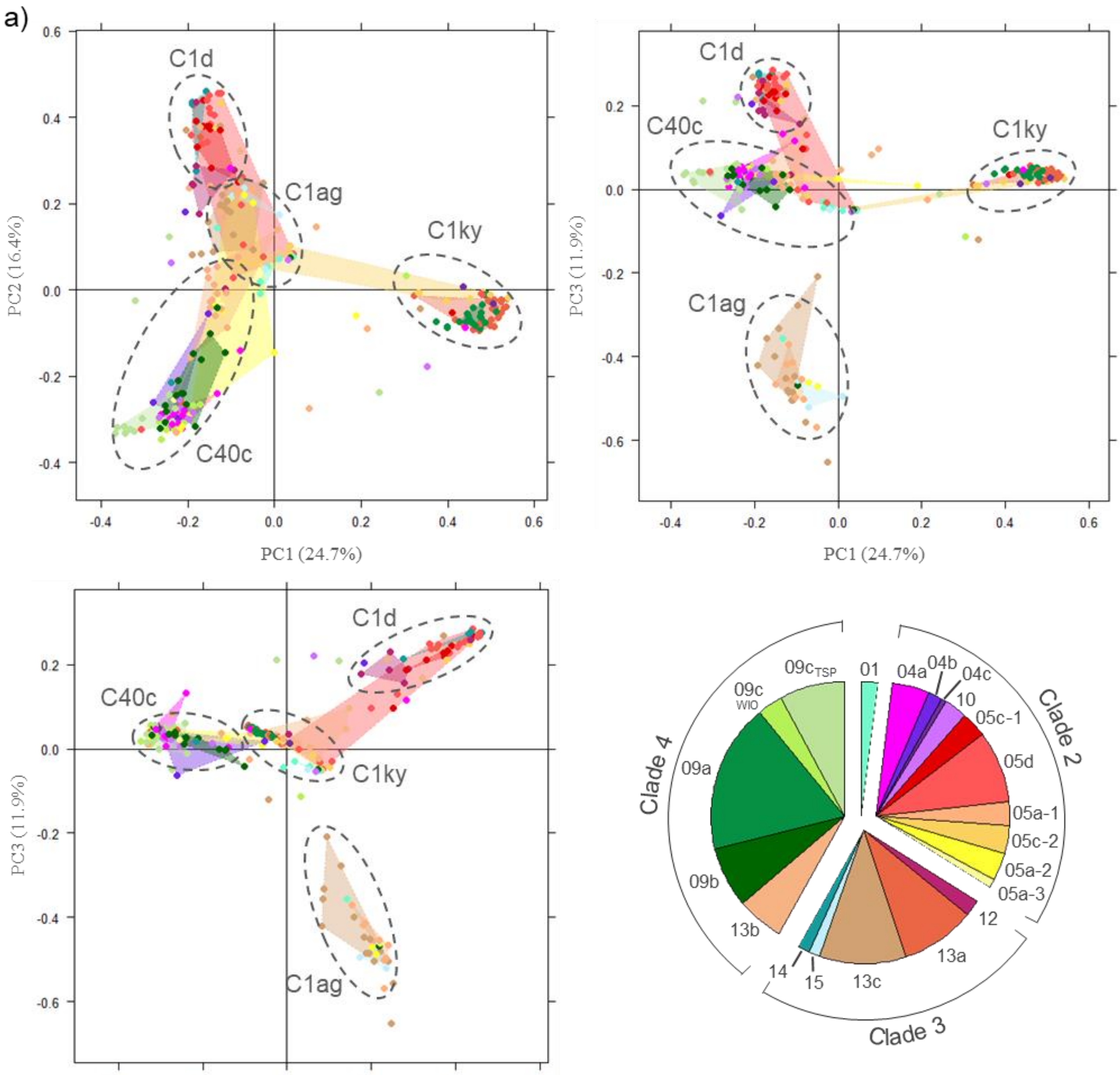

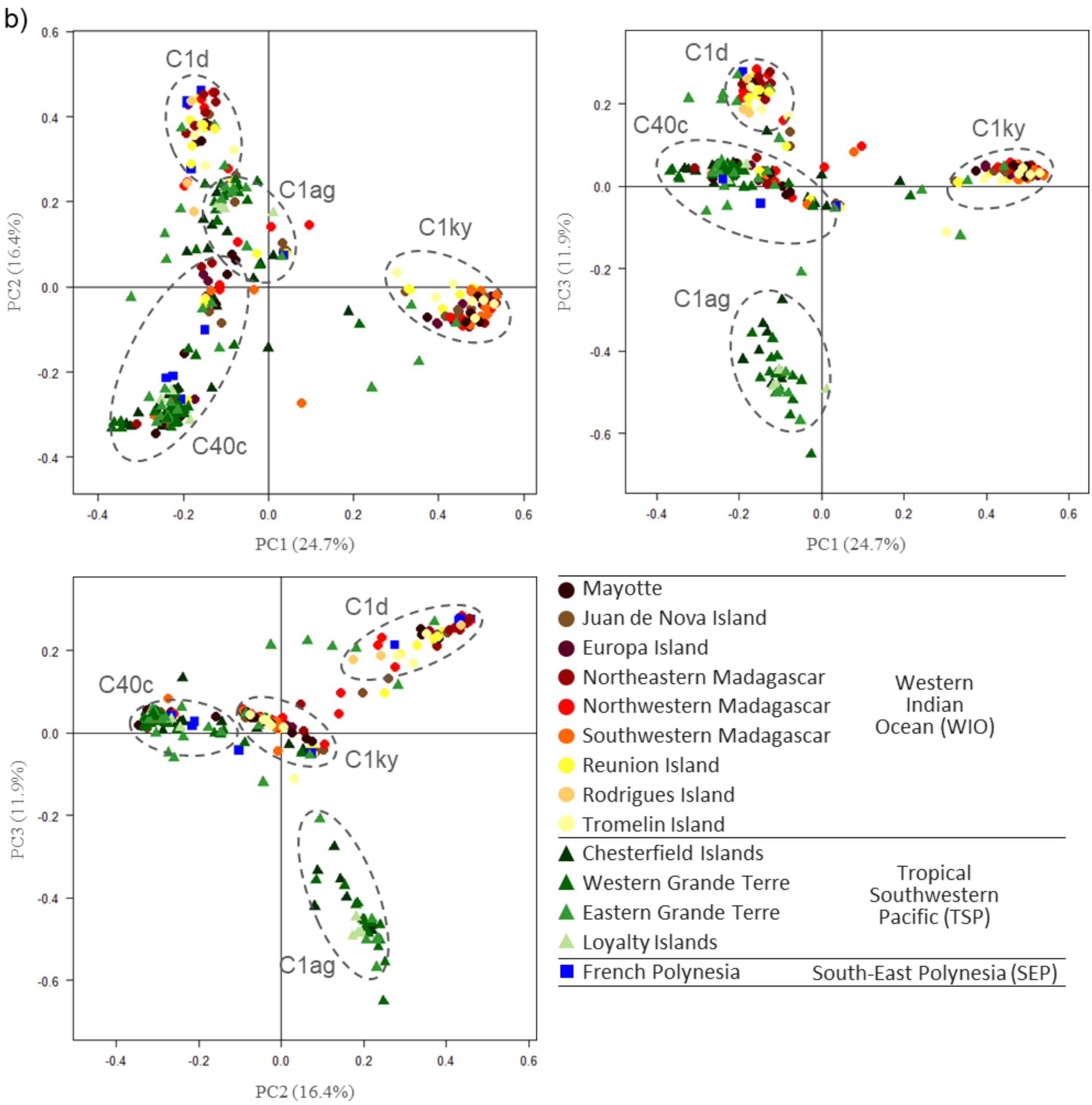
