## Appendix 5: Illustration of the Pocillopora Genomic Species Hypotheses for "From Genomics to Integrative Taxonomy? The Case Study of *Pocillopora* Corals"

#### **Appendix 5** Illustration of the *Pocillopora* Genomic Species Hypotheses (GSHs).

### GSH01

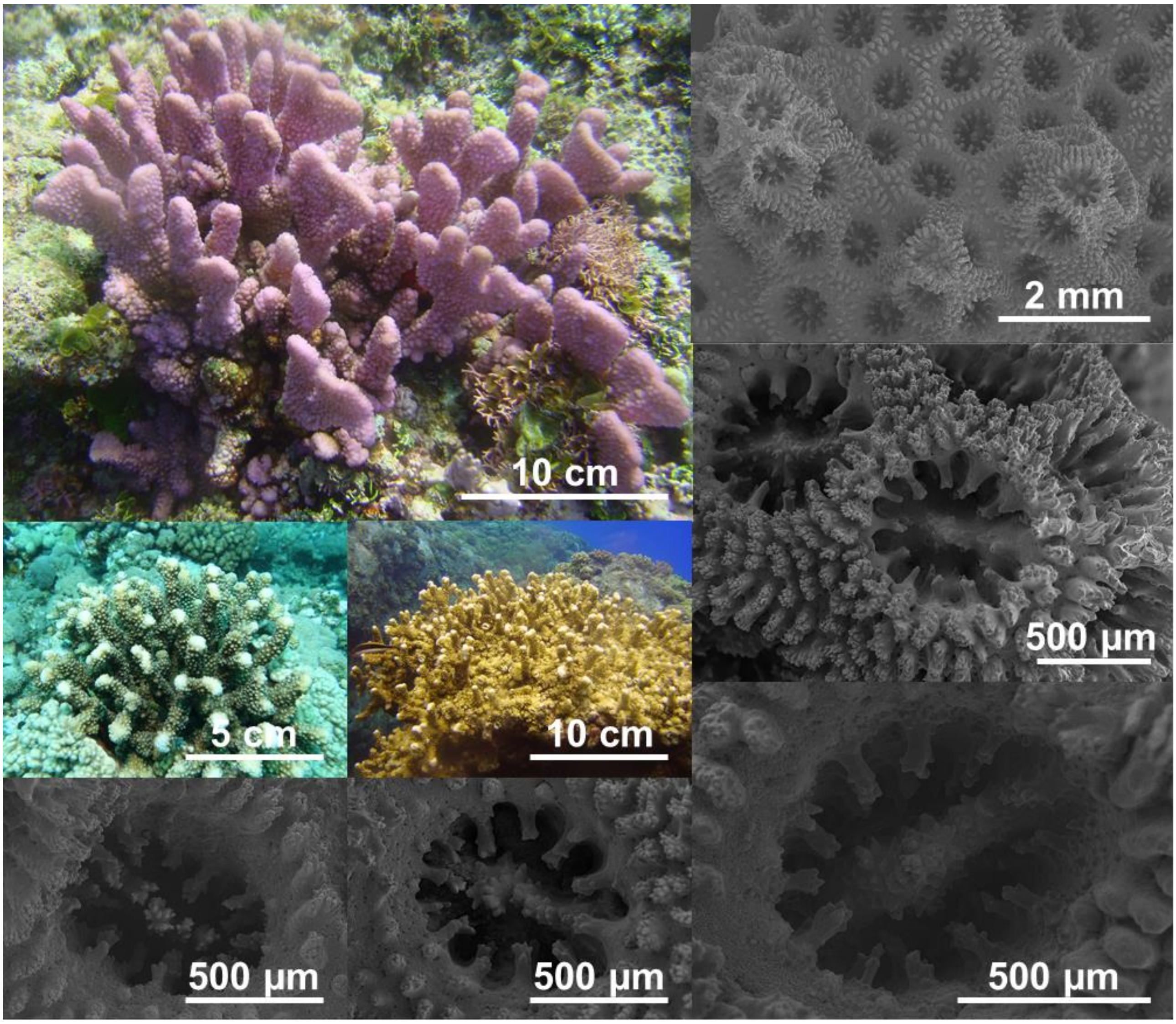

### GSH04a

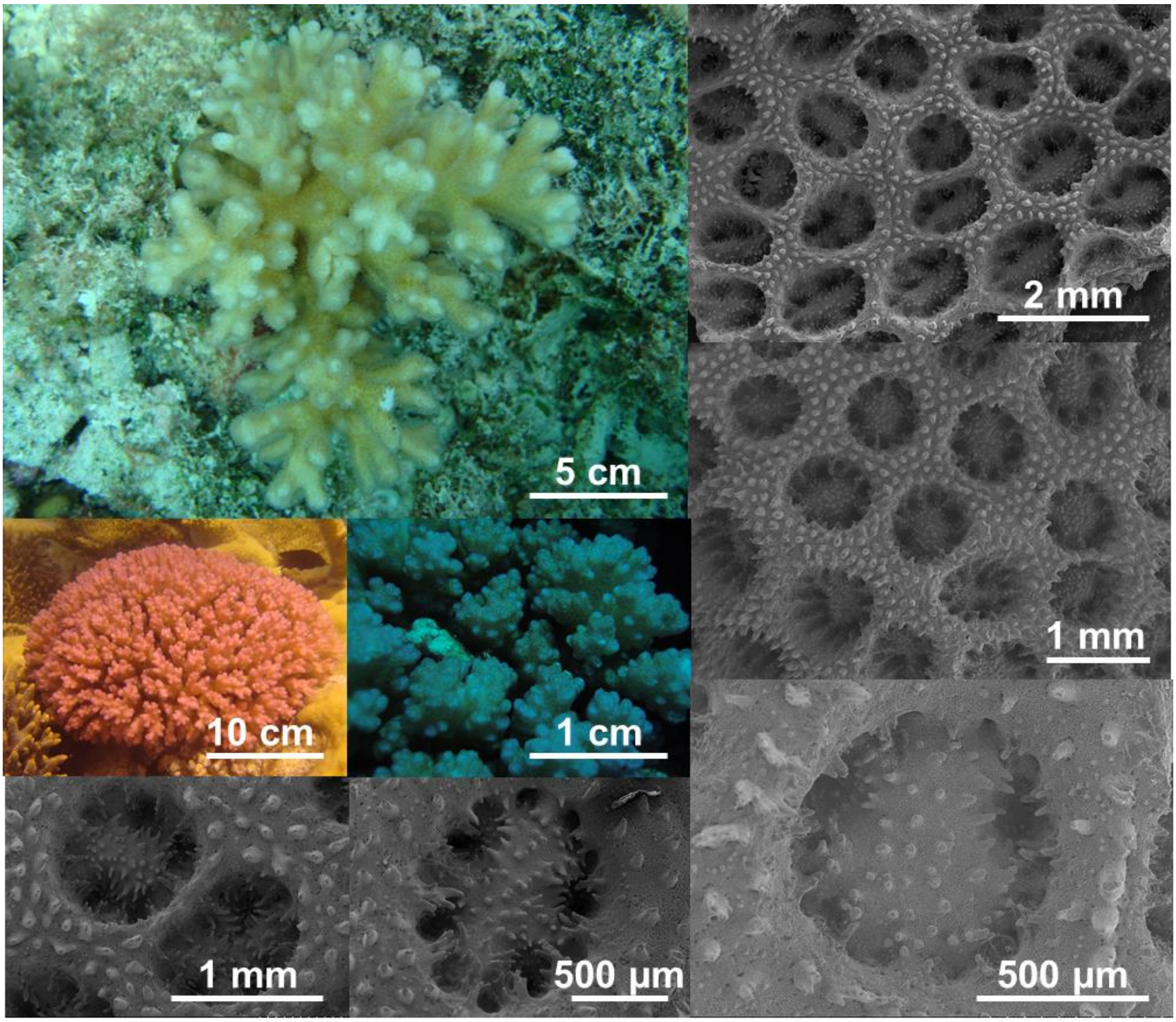

### GSH04b

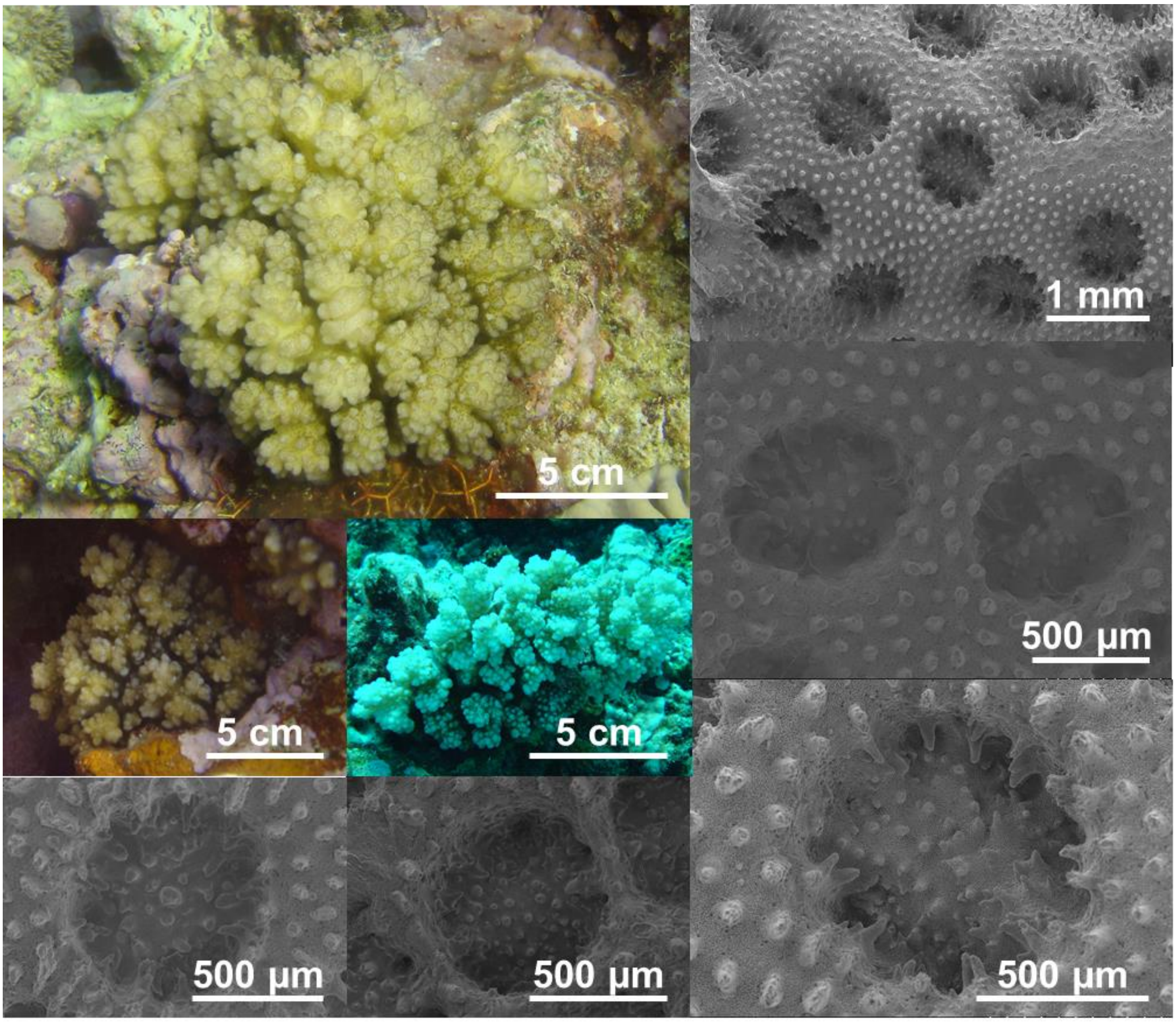

### GSH04c

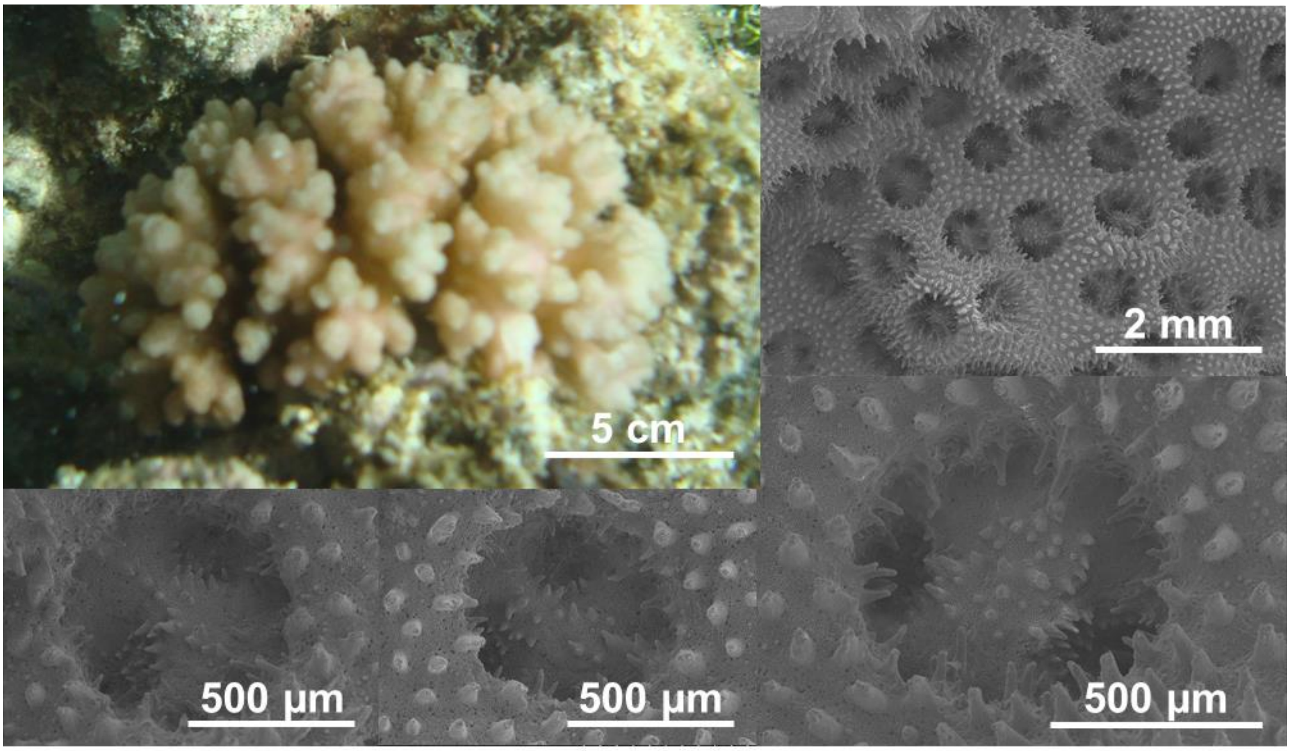

### GSH05a-1

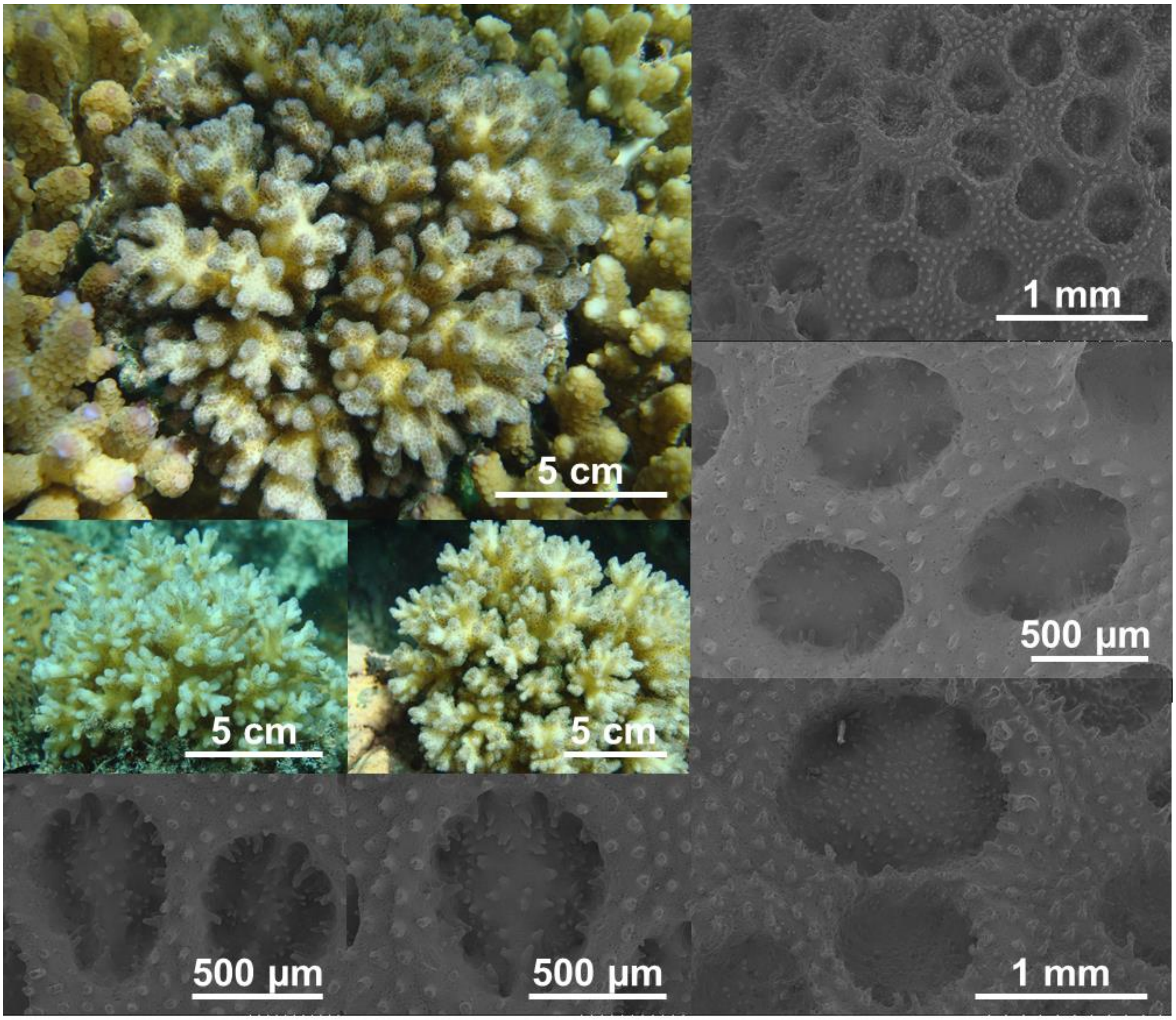

### GSH05a-2

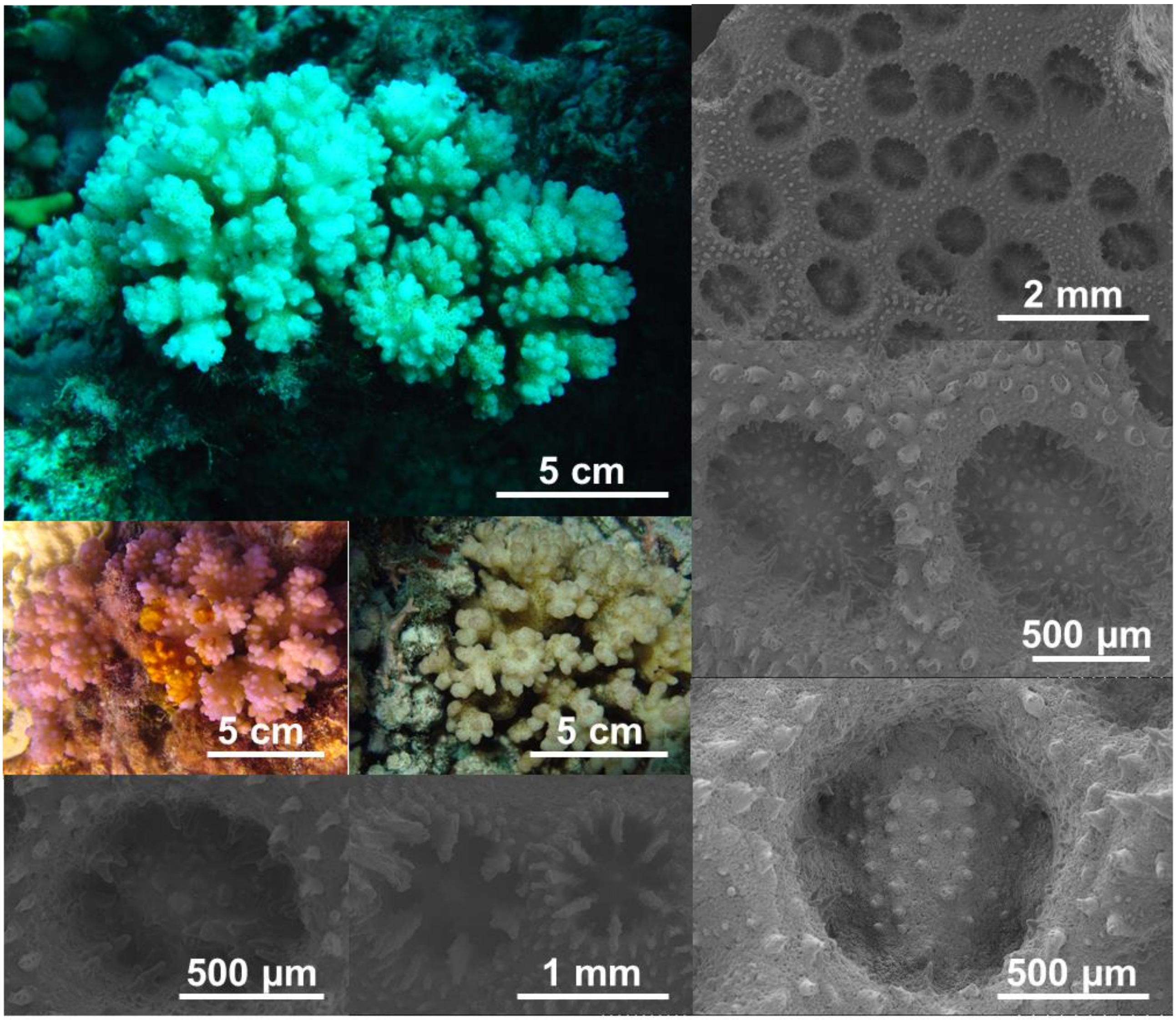

### GSH05a-3

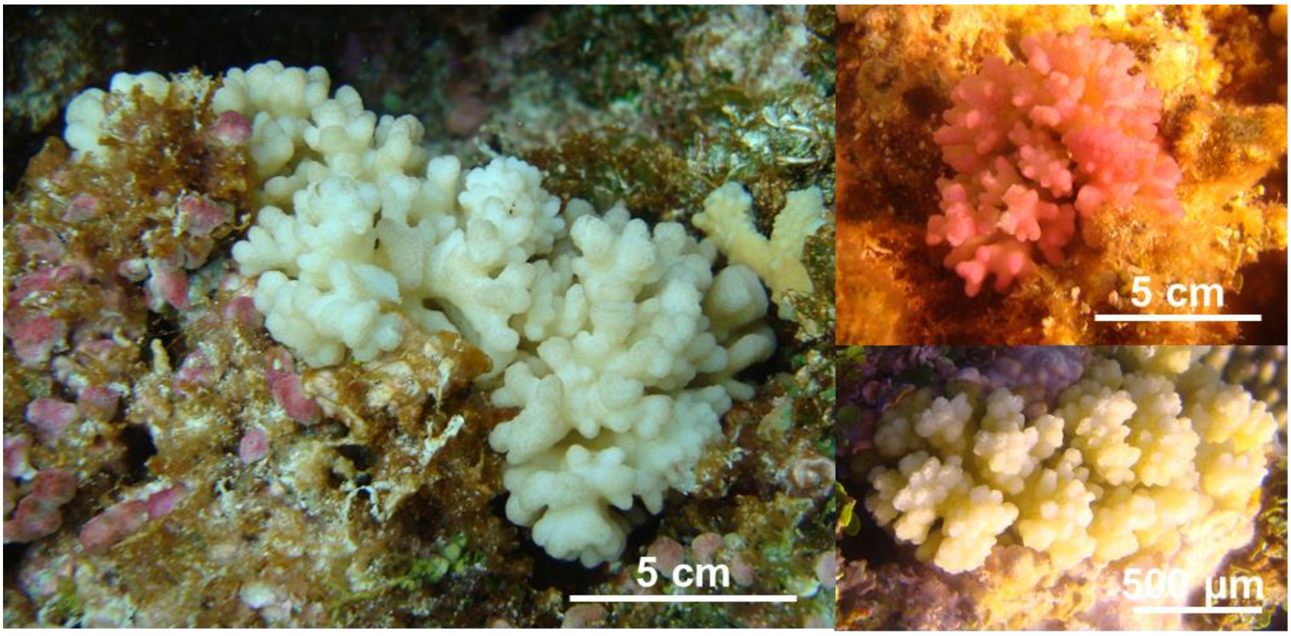

### GSH05c-1

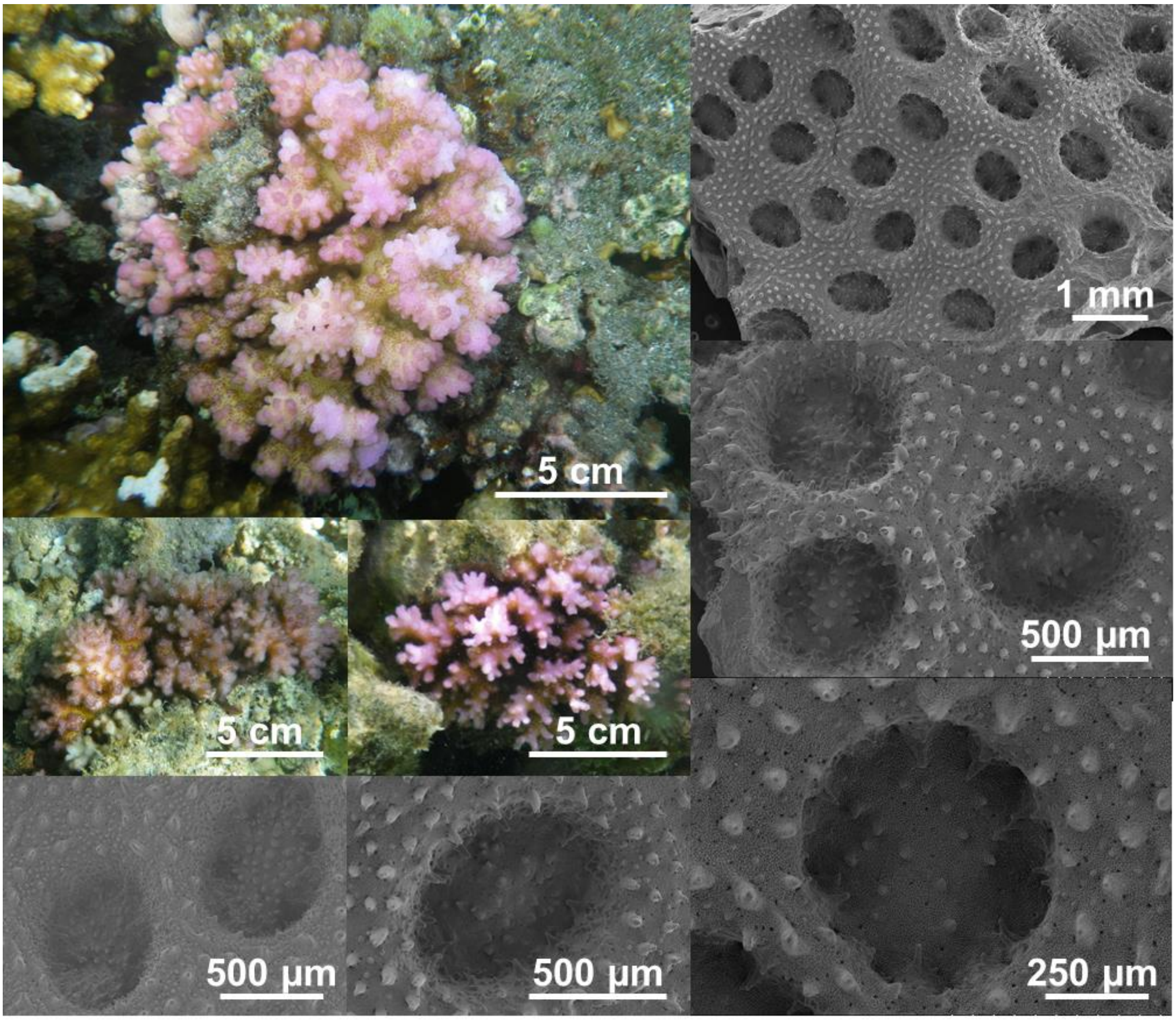

### GSH05c-2

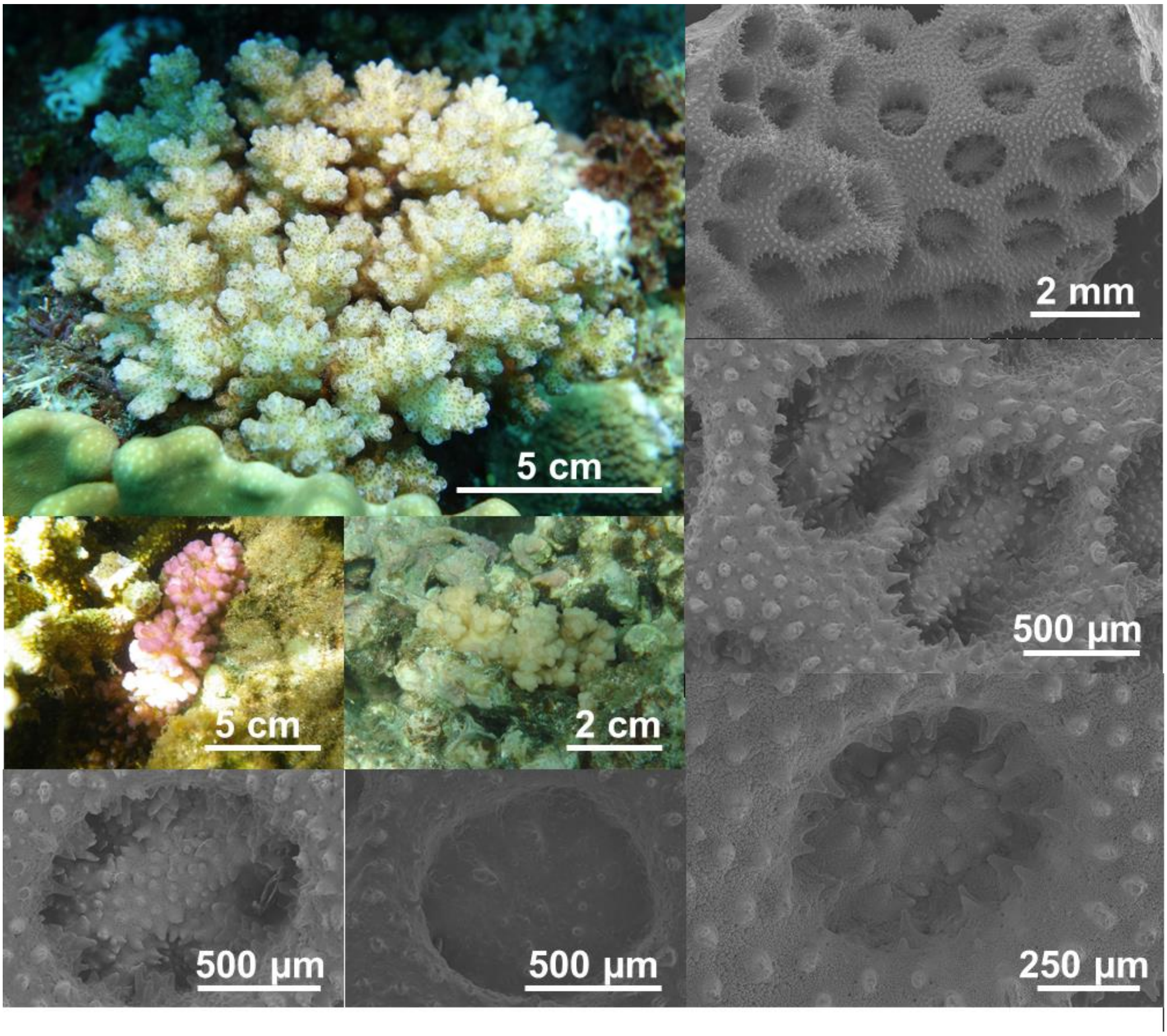

### GSH05d

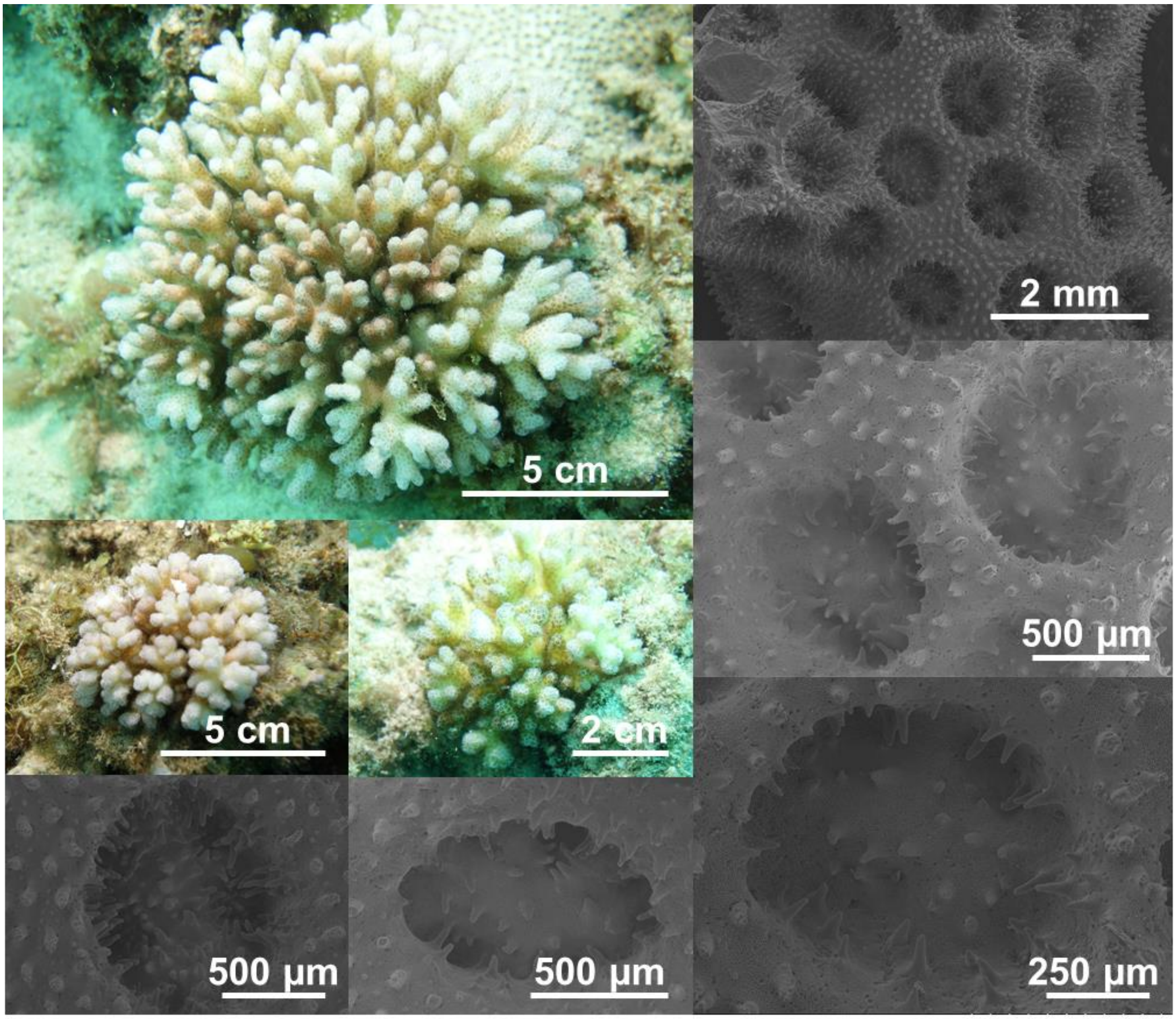

### GSH09a

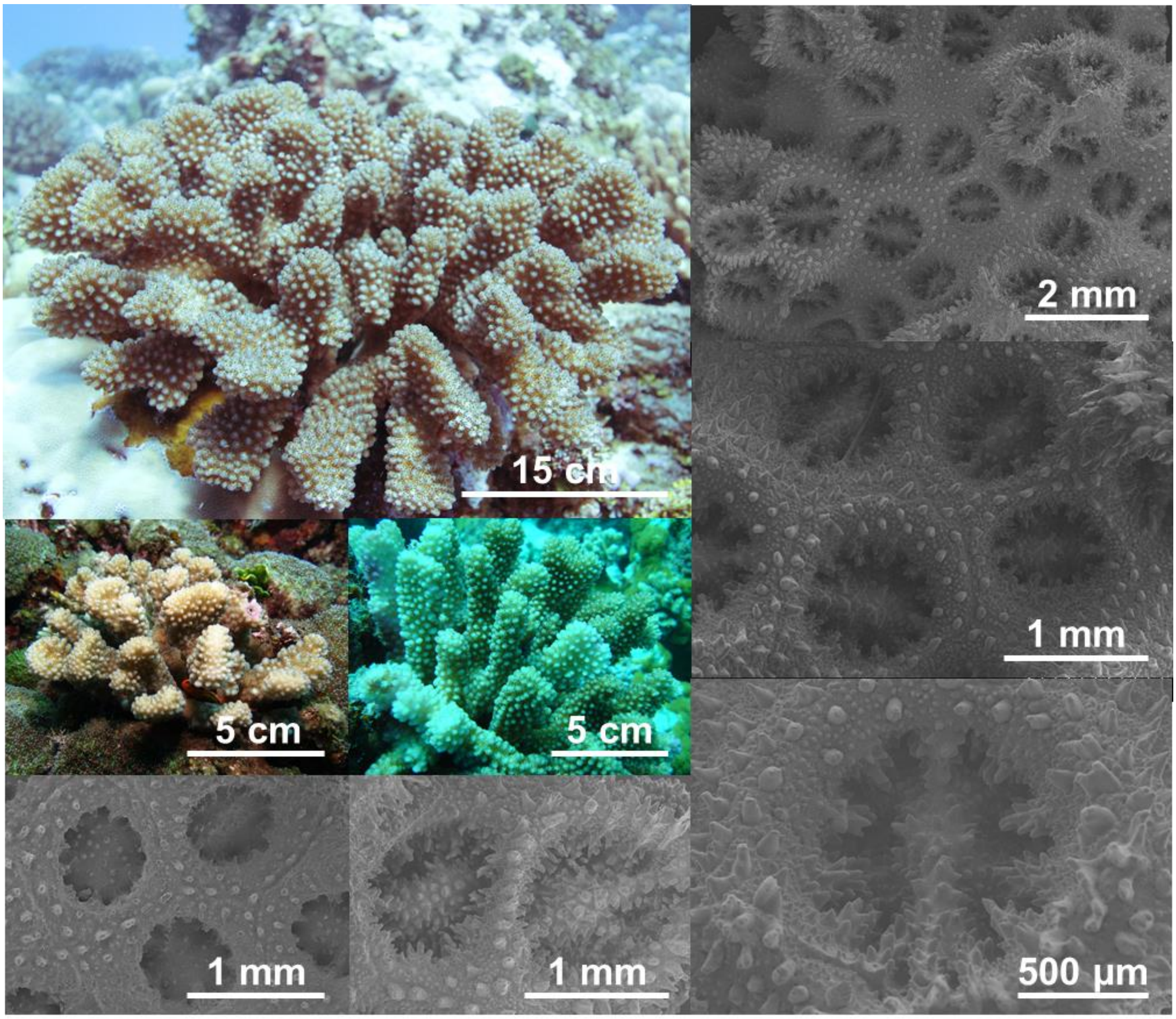

### GSH09b

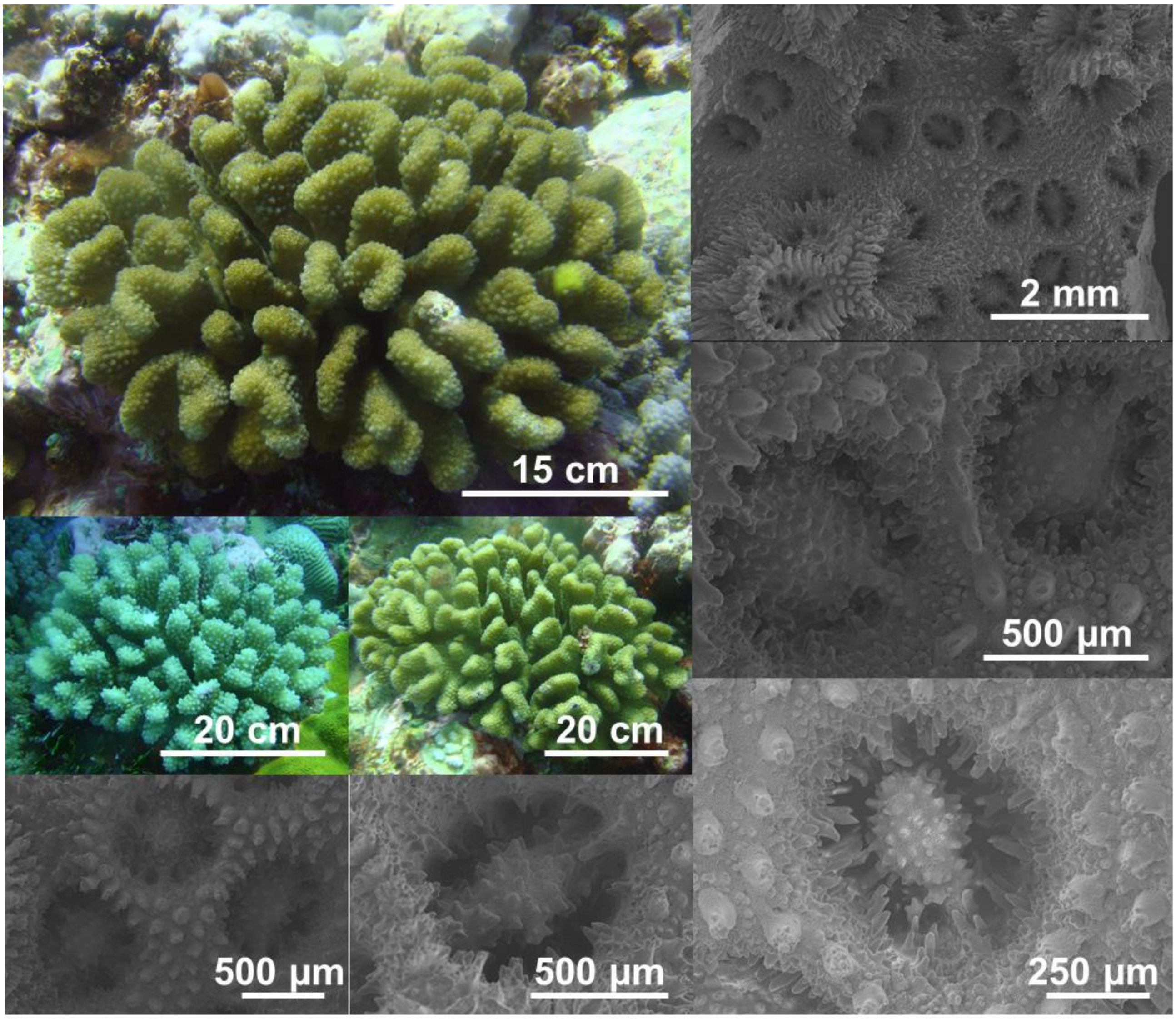

### GSH09c$_{WIO}$

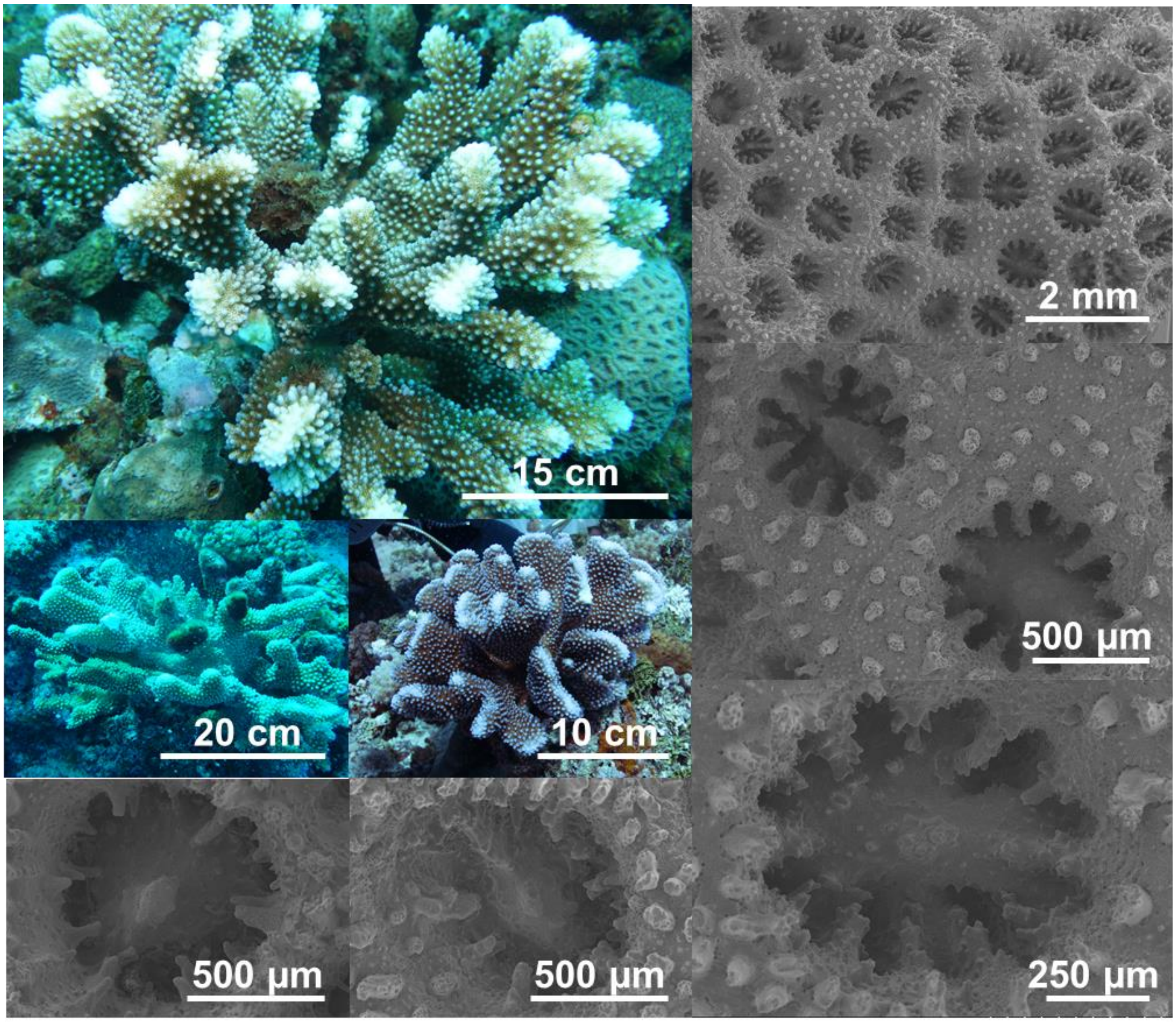

### GSH09c$_{TSP}$

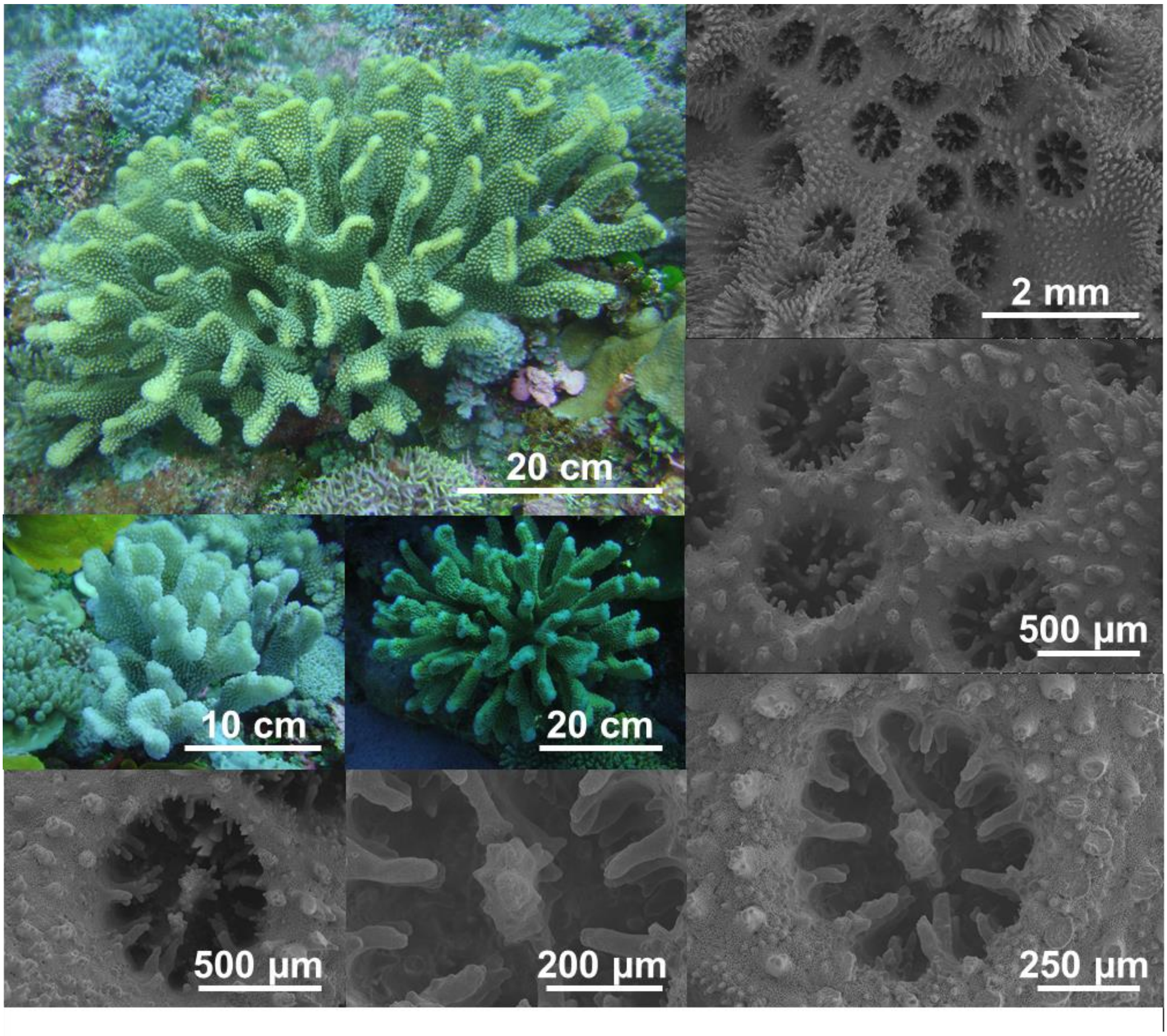

### GSH10

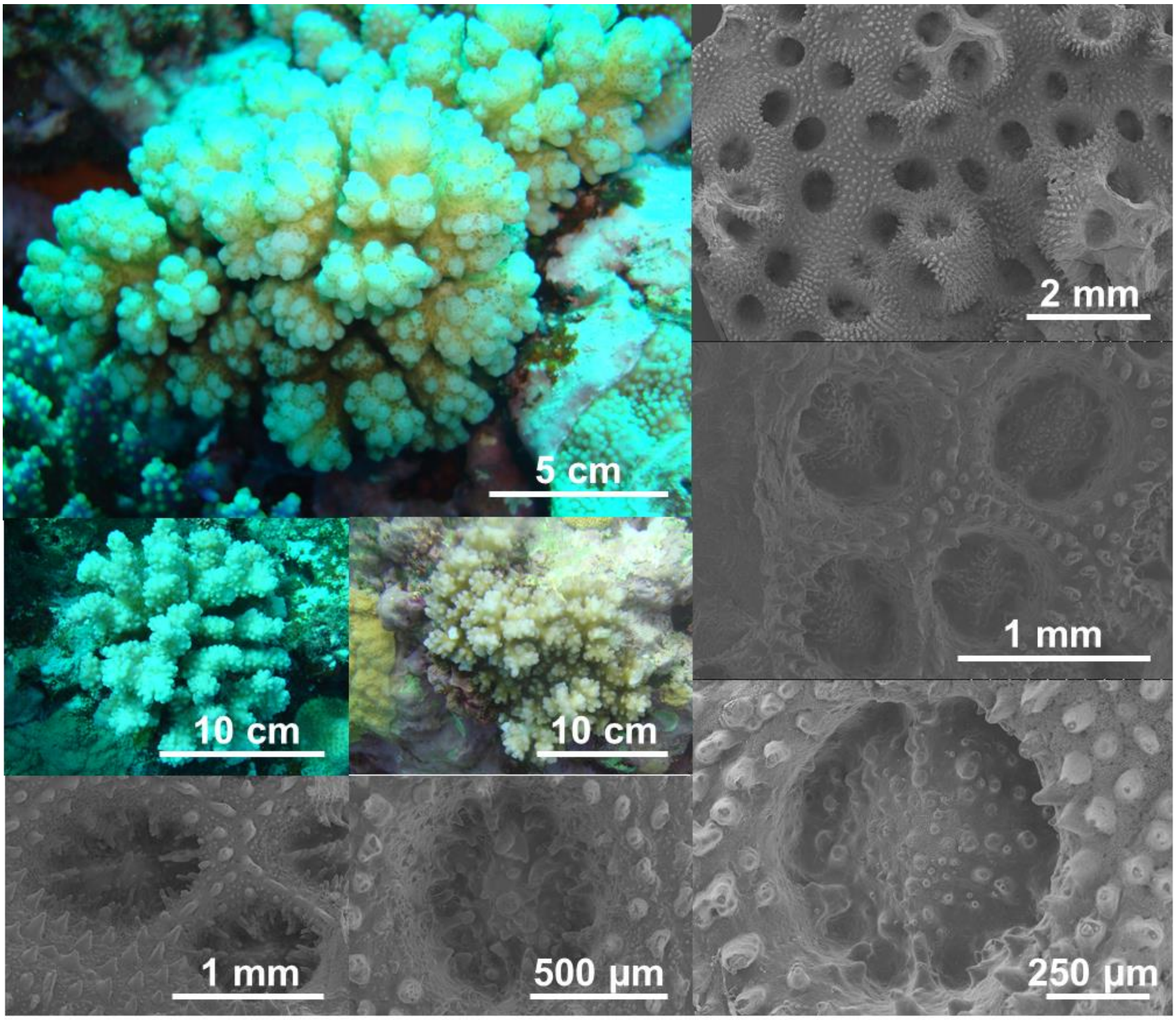

### GSH12

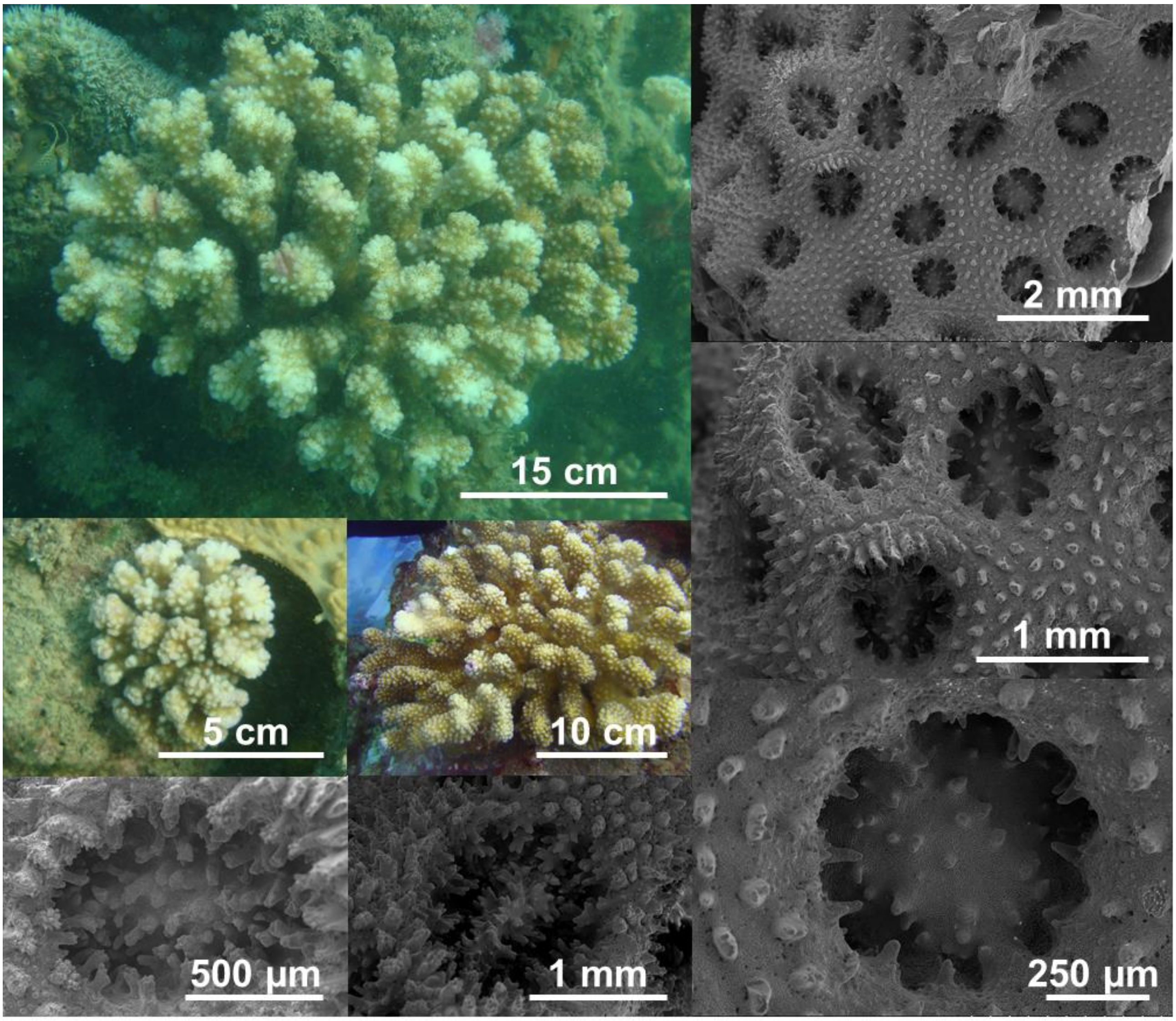

### GSH13a

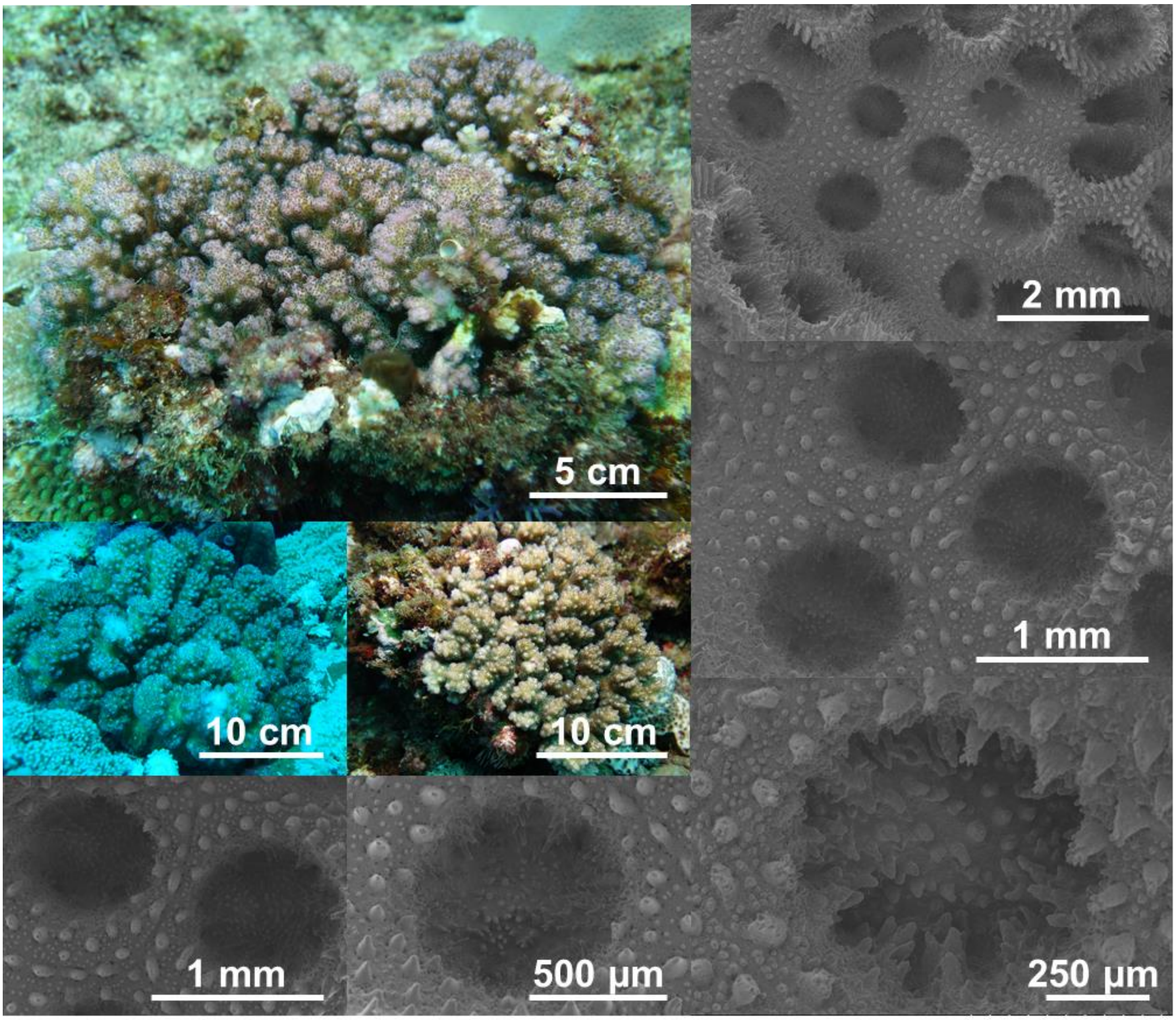

### GSH13b

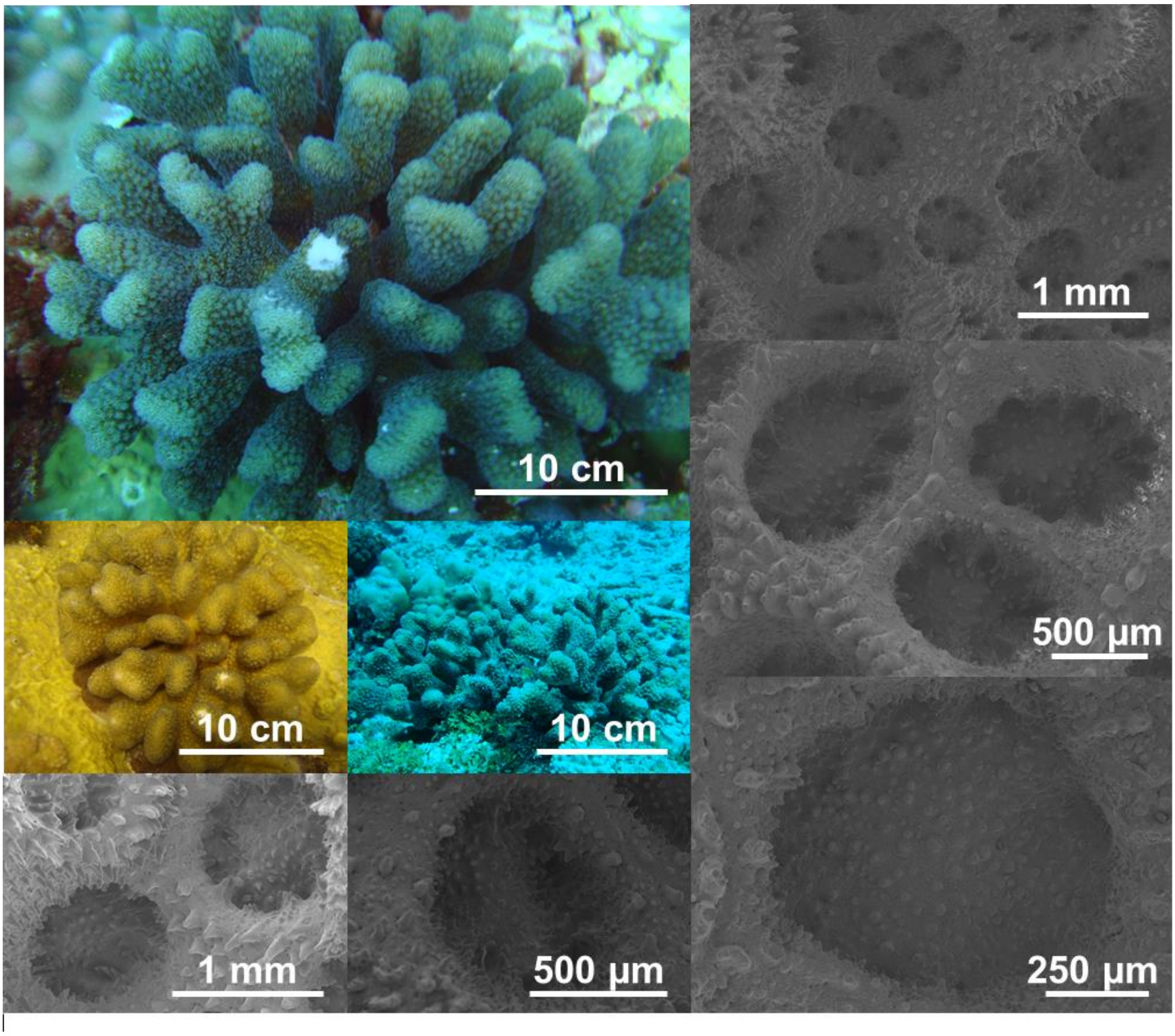

### GSH13c

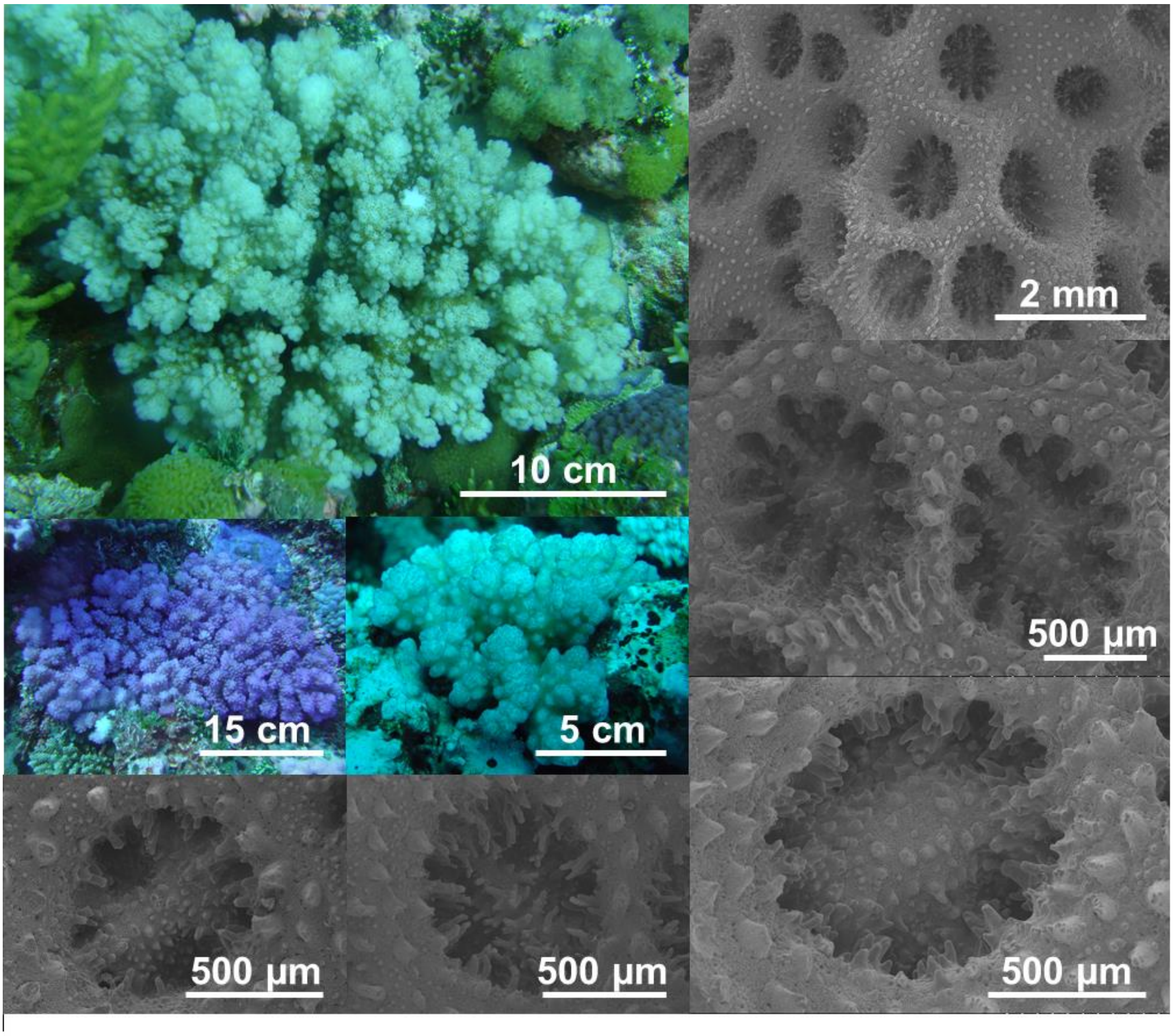

### GSH14

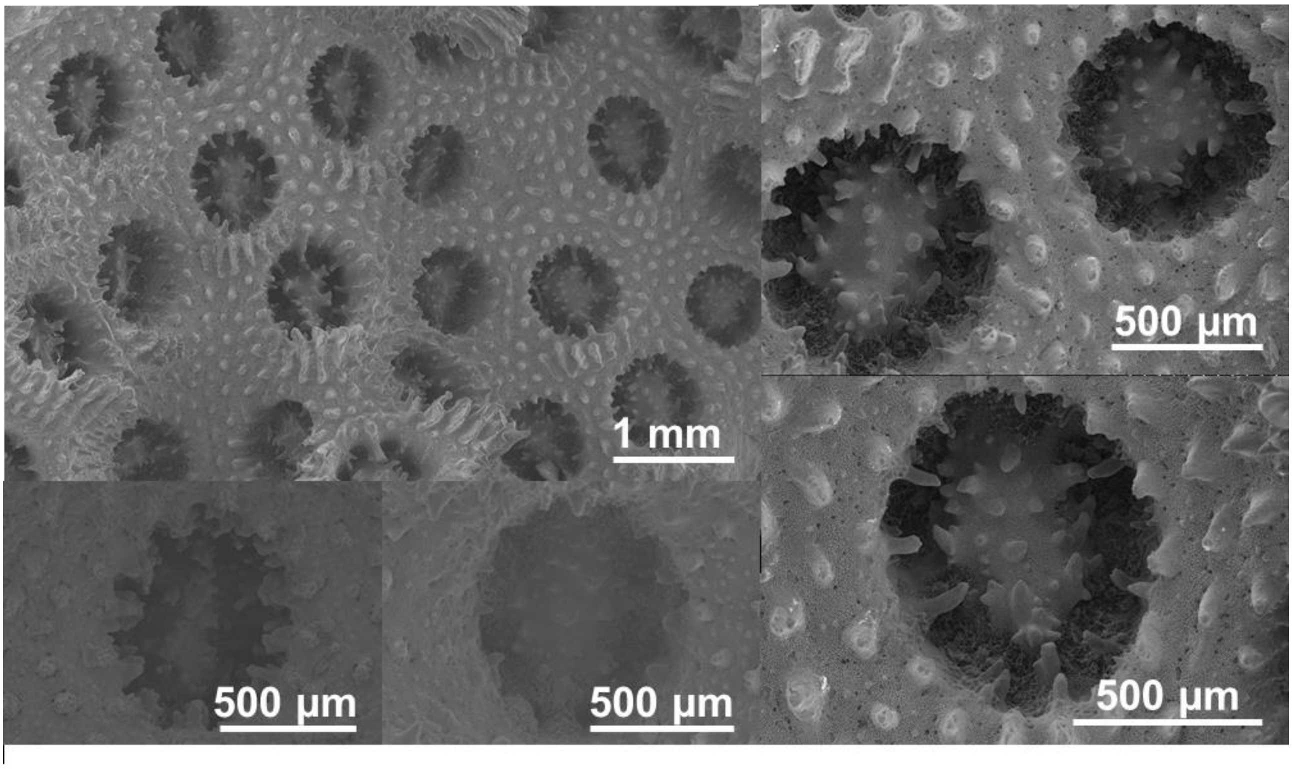

### GSH15

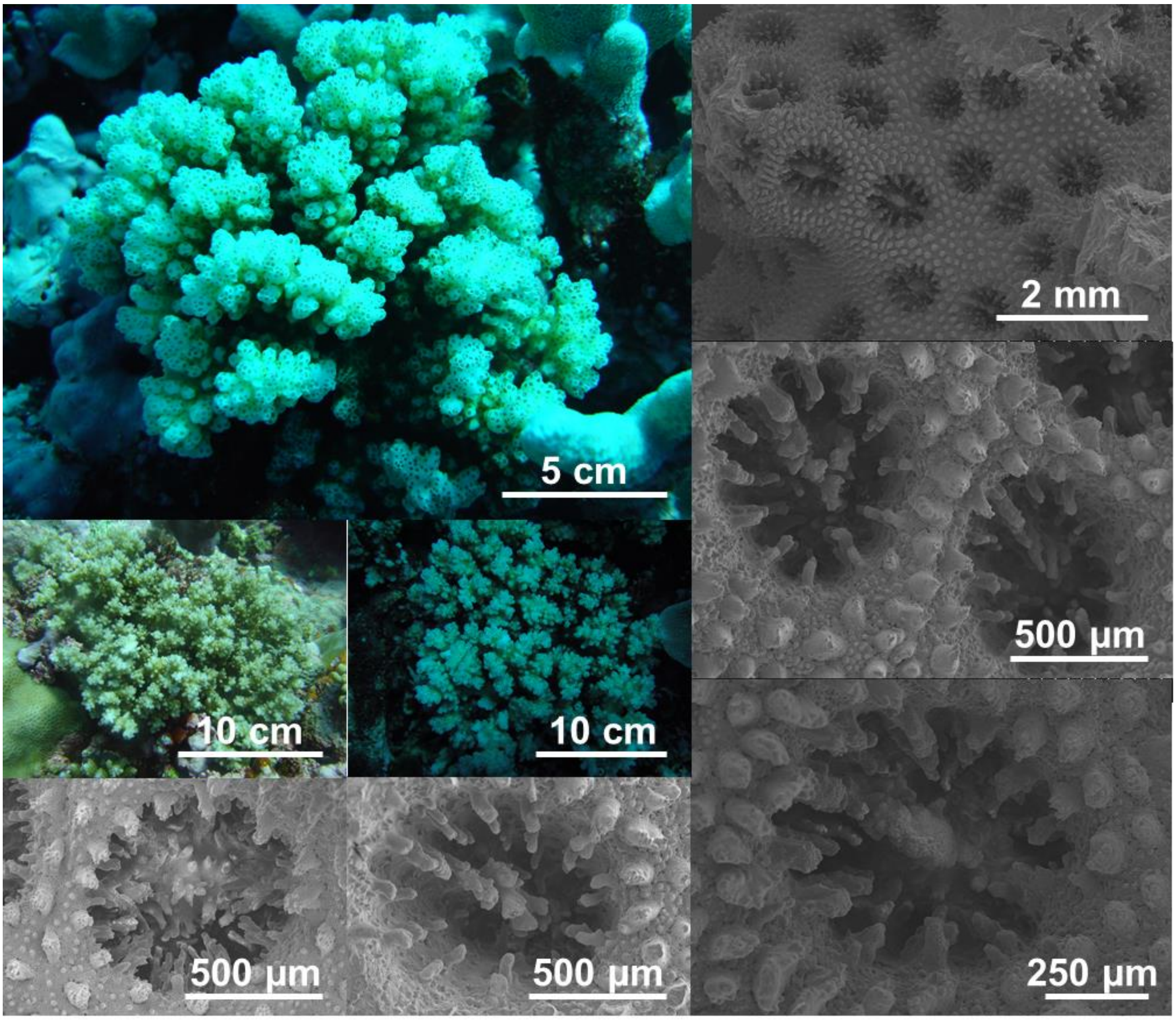
